# The Prevalence of Killer Yeasts in the Gardens of Fungus-Growing Ants and the Discovery of a Novel Killer Toxin named Ksino

**DOI:** 10.1101/2024.10.14.618321

**Authors:** Rodolfo Bizarria, Jack W. Creagh, Renato A. Corrêa dos Santos, Lily LGivens, Sarah A. Coss, Tanner J. Badigian, Adrian V. Chavez, Rim T. Tekle, Noah Fredstrom, F. Marty Ytreberg, Maitreya J. Dunham, Andre Rodrigues, Paul A. Rowley

## Abstract

Killer toxins are proteinaceous antifungal molecules produced by yeasts, with activity against a wide range of human and plant pathogenic fungi. Fungus gardens of attine ants in Brazil were surveyed to determine the presence of killer toxin-producing yeasts and to define their antifungal activities and ecological importance. Our results indicate that 10 out of 59 yeasts species isolated from fungal gardens are killer yeasts. Killer yeasts were less likely to inhibit the growth of yeasts isolated from the same environment but more effective at inhibiting yeast isolated from other environments, supporting a role for killer yeasts in shaping community composition. All killer yeasts harbored genome-encoded killer toxins lacked cytoplasmic toxin-encoding elements (i.e., double-stranded RNA satellites and linear double-stranded DNAs). Of all the killer yeasts associated with attine ants, *Candida sinolaborantium* (strain LESF 1467) showed a broad spectrum of antifungal activities against 39 out of 69 57% of yeast strains tested for toxin susceptibility. The complete genome sequence of *C. sinolaborantium* LESF 1467 identified a new killer toxin, Ksino, with similarities in primary sequence and tertiary structure to the *Saccharomyces cerevisiae* killer toxin named Klus. Surveys of publicly available genome databases identified homologs of Ksino in the genomes of yeast strains of *Saccharomycetes* and *Pichiomycetes*, as well as other species of Ascomycota and Basidiomycota filamentous fungi. This demonstrates that killer yeasts can be widespread in attine ant fungus gardens, possibly influencing fungal community composition and the importance of these complex microbial communities for discovering novel antifungal molecules.

**Importance:** Attine ants perform essential ecosystem services through the harvesting of substrates for fungiculture. The cultured fungi are a food source for attine ants. Characterizing antifungal toxin-producing yeasts (killer yeasts) is vital to understanding how they might protect gardens from invasion by unwanted fungal species. This study describes a new toxin named Ksino from the yeast *Candida sinolaborantium*, a member of a new group of toxins found across many different species of fungi. This work supports the role of killer yeasts in the ecology of fungicultures and competition between fungi. The observed high prevalence of killer yeasts in fungal gardens also enables the discovery of novel antifungal molecules with the potential to be applied against disease-causing fungi.

## 1. Introduction

Killer yeasts were first described in *Saccharomyces cerevisiae* in the 1960s (Makower and Bevan, 1963; Woods and Bevan, 1968), after which many strains and species of yeasts and yeast-like fungi were also observed to secrete antifungal killer toxins (Woods and Bevan, 1968; Golubev et al., 2002; Pfeiffer et al., 2004; Koltin and Day, 1976; Castillo and Cifuentes, 1994). Killer toxins can have a range of antifungal activities ranging from narrow to broad specificities, affecting cells from close or distantly related fungal species (Boynton, 2019; Golubev, 1998; Golubev, 2006). The prevalence of killer yeasts strains in natural environments is reported between 5% to 33% for positive killer phenotypes (Philliskirk and Young, 1975; Stumm et al., 1977; Starmer et al., 1987; Abranches et al. 1997, Abranches et al. 1998; Trindade et al. 2002; Pieczynska et al., 2013; Wojcik and Kordowska-Wiater, 2015) but can also be as high as 59% in yeast associated with wine production (Hidalgo and Flores, 1994). Killer toxins are known to be active against human and plant pathogenic yeasts (Middelbeek et al. 1980; Hodgson et al. 1995; Buzzini et al., 2001; Giovati et al., 2018) and can control spoilage organisms that are important for agriculture and food industry (Walker et al., 1995; Chessa et al., 2017). The rise in acquired drug resistance and the emergence of drug-resistant fungal pathogens justifies the search for novel antifungal killer toxins from diverse natural environments, including insect-associated fungal communities.

Killer yeast communities have been previously reported in fungus-growing ant colonies (Hymenoptera: tribe Attini: subtribe Attina, hereafter called ‘attine ants’). Attine ants established a long-term obligate mutualism with basidiomycete fungi in the *Agaricaceae* and *Pterulaceae* families, using them as the primary source of food for their colonies (Branstetter et al., 2017; Nygaard et al., 2016; Schultz and Brady, 2008, Möller, 1893; Weber, 1972). According to their cultivars and foraging substrates, each species approaches fungiculture differently, and these interactions are considered a model for studying the evolution of mutualisms (Schultz and Brady, 2008, Schultz et al., 2015). Moreover, attine ants are key ecological agents in the Neotropics as dominant herbivores, promoting nutrient cycling, soil fertility, and seed dispersal (Corrêa et al., 2010; Meyer et al., 2011, Farji-brener and Werenkraut, 2015; Sternberg et al., 2007; Leal and Oliveira, 1998). The foraging behavior of attine ants also contributes to the presence of a large and complex community of environmental yeasts in fungus gardens (Craven et al. 1970; Fisher et al., 1996; Pagnocca et al., 2008; Arcuri et al., 2014; Bizarria Jr. et al., 2022). Fungus gardens are likely an important source of new killer toxins due to the high concentration of simple sugars (Silva et al., 2003, Silva et al., 2006), a large density of yeasts and filamentous fungi (Fisher et al., 1996; Craven et al., 1970; Pagnocca et al., 1996; Rodrigues et al., 2008; Bizarria Jr. et al., 2022), and the hypothesized role on garden “immunity” by yeasts (Rodrigues et al. 2009) – all key elements for species competition (Boynton, 2019). Killer yeasts are primarily associated with the basidiomycete fungus farmed by attine ants, the bodies of ants, and the leaves foraged by the ants (Carreiro et al. 2002; Robledo-Leal et al. 2016; Bizarria Jr. et. al. 2023). In prior studies, killer positive phenotypes have been identified in 5% to 11% of the interactions between yeasts derived from attine ant fungus gardens (Carreiro et al. 2002; Bizarria Jr. et. al. 2023).

Killer toxins can modify the community composition of yeasts as an outcome of competition or cooperation between killer yeasts, killer toxin-susceptible yeasts, and killer toxin-resistant yeasts that are determined by spatial distribution, pH, and ploidy (Károlyi et al., 2005; McBride et al. 2008; Wloch-Salamon et al. 2008). Killer toxin production also plays a potential role in yeast dispersal, resource consumption among species, and invade or prevent invasions from competitors (Boynton, 2019; Buser et al., 2021; Pintar and Starmer, 2003; Ganter and Starmer, 1992; Travers-Cook et al., 2023). The coexistence of the different phenotypes is known to be affected by environmental parameters, including nutrient availability (Pintar and Starmer, 2003; Wloch-Salamon et al. 2008), cell density (Greig and Travisano 2008; Vadasz et al. 2003), pH, and temperature (Bussey et al. 1988; McBride et al. 2008; Magliani et al. 1997), which can reflect the production and distribution of killer toxins in spatially structured environments (Boynton, 2019; Giometto et al. 2021; Kerr et al. 2002; Libberton et al. 2015; Wloch-Salamon et al. 2008). On the other hand, killer toxins are often reported to play a role in interference competition in environments that enforce the interaction between toxin-producing and neighboring susceptible cells (such as well-mixed habitats like laboratory co-cultures and wine fermentation, changing the community structure and function, as an outcome of competition or cooperation between killer yeasts, killer toxin-susceptible yeasts, and killer toxin-resistant yeasts (Bussey et al., 1988; Hidalgo and Flores, 1994; Pieczynska et al., 2016; Boynton, 2019).

Killer toxins are encoded by different genetic elements, including chromosomal genes, extrachromosomal elements (such as non-autonomous double-stranded RNA (dsRNA) satellites that are maintained by mycoviruses), and autonomous linear double-stranded DNAs (dsDNA) (Bevan et al., 1973; Somers and Bevan, 1969; Magliani et al., 1997; Rowley, 2017). DsRNA satellites have been previously found to be associated with *Hanseniaspora uvarum*, *Pichia kluyveri*, *Saccharomyces* spp., *Torulaspora delbrueckii*, and *Zygosaccharomyces bailii* (Ivannikova et al 2006, Ivannikova et al 2007; Naumov et al., 2009; Schmitt and Neuhausen, 1994; Ramírez et al., 2015; Rodríguez-Cousiño et al., 2017; Zorg et al., 1988), while dsDNA plasmids have been identified in the genera *Babjevia, Debaryomyces*, *Kluyveromyces*, *Millerozyma*, and *Pichia* (Klassen and Meinhardt 2007; Schaffrath et al. 2018; Klassen et al. 2017). The association of the killer phenotype with the presence of dsRNA satellites has been most thoroughly studied in *S. cerevisiae* (Crabtree et al., 2023; Vijayraghaven et al., 2023). For example, in a survey of 1,270 *S. cerevisiae* isolates, the frequency of killer yeasts was 50%, and 60% of these killer yeasts had dsRNA satellites (Crabtree et al., 2023). Killer toxin production by the remaining 40% of killer yeasts was closely correlated with the chromosomal killer toxin gene *KHS1*. Considering the many killer yeasts described, only a few studies have defined chromosomal genes that encode the toxins responsible for the observed antifungal effects. Chromosomally encoded killer toxins and their homologs, including K1-like (KKT killer toxins), KHR, KHS, SMKT, KP4-like, PMKT, and other *Pichia* toxins, have been described in many different species of fungi (Goto et al., 1990; Goto et al., 1991; Suzuki and Nikkuni, 1994; Belda et al., 2017, Fredericks et al. 2021, Brown 2011, Lu and Faris 2019). Despite the prevalence of genomic-encoded toxins, some dsRNA satellite-encoded toxins were observed to have genomic homologs in different Saccharomycotina lineages (Frank and Wolfe, 2009; Fredericks et al., 2021). However, the frequency of killer yeasts and the prevalence of dsRNA satellites and dsDNA extrachromosomal elements in different lineages of Saccharomycotina and yeast-like fungi is still understudied.

In this study, we explore the nature and prevalence of the killer phenotype in different Saccharomycotina lineages associated with attine ant fungus gardens, which are habitats that are rich in yeasts, shedding light on their putative ecological role in this environment (Bizarria Jr. et al. 2023). It was revealed that fungus gardens harbor diverse killer yeasts with narrow and broad antifungal activities. Unlike the well-studied killer yeasts of the *Pichia* and the *Saccharomyces* genera, attine ant-associated killer yeasts were devoid of cytoplasmic toxin-encoding elements such as linear dsDNA plasmids and dsRNA satellites (Magliani et al., 1997; Marquina et al., 2002). Therefore, to identify the gene(s) responsible for killer toxin production, a genome mining approach was used by determining the whole genome sequence of the killer yeast *Candida sinolaborantium* that displayed a broad spectrum of antifungal activities against many species of yeasts, including human pathogens. This approach identified a new killer toxin named Ksino. This novel killer toxin has homologs in both yeasts and filamentous fungi and shares a similar antifungal activity and predicted structure to an *S. cerevisiae* killer toxin named Klus. Collectively, it was found that fungus gardens of attine ants can be used to explore the ecological role of killer yeasts and discover new killer toxins for possible future application as natural antifungals.

## 2. Results

### Fungus gardens of attine ants harbor killer yeasts

To assess the diversity of yeasts associated with fungus-growing ants and their potential for antifungal toxin production, fungus gardens of four ant fungiculture systems were surveyed from 28 nests from four cities in Brazil (Fig. 1A; Table S1). The ant species included the leaf-cutting ant *Acromyrmex coronatus*, the non-leaf-cutting ant *Mycetomoellerius tucumanus*, and the lower attines *Mycetophylax* aff. *auritus* and *Mycocepurus goeldii* that cultivate Agaricaceae fungi, and *Apterostigma goniodes* that cultivates Pterulaceae fungi (Fig. 1B). Ants were selected to broadly represent different fungiculture systems, foraging and preparation behaviors, and the yeast microbiota associated with fungus gardens. Yeasts were isolated from fungus gardens by culturing on synthetic growth media. Microsatellite amplification, sequencing of the D1/D2 domain of the large subunit ribosomal RNA gene, and phylogenetic reconstruction were used for taxonomy. The 180 isolated yeast strains comprised 59 species belonging to eight families and six orders from the Saccharomycotina subphylum previously characterized in a large study of yeasts and yeast-like fungi associated with fungus gardens of attine ants (Fig. 1C; Table S1) (Bizarria Jr. et al., 2023). Of the yeasts isolated, 91 were associated with *Acromyrmex coronatus*, 13 with *Mycetomoellerius tucumanus*, 42 with *Mycetophylax* aff. *auritus*, 25 with *Mycocepurus goeldii*, and nine with *Apterostigma goniodes* (Table S1) (Bizarria Jr. et al., 2023).

**FIG 1.**
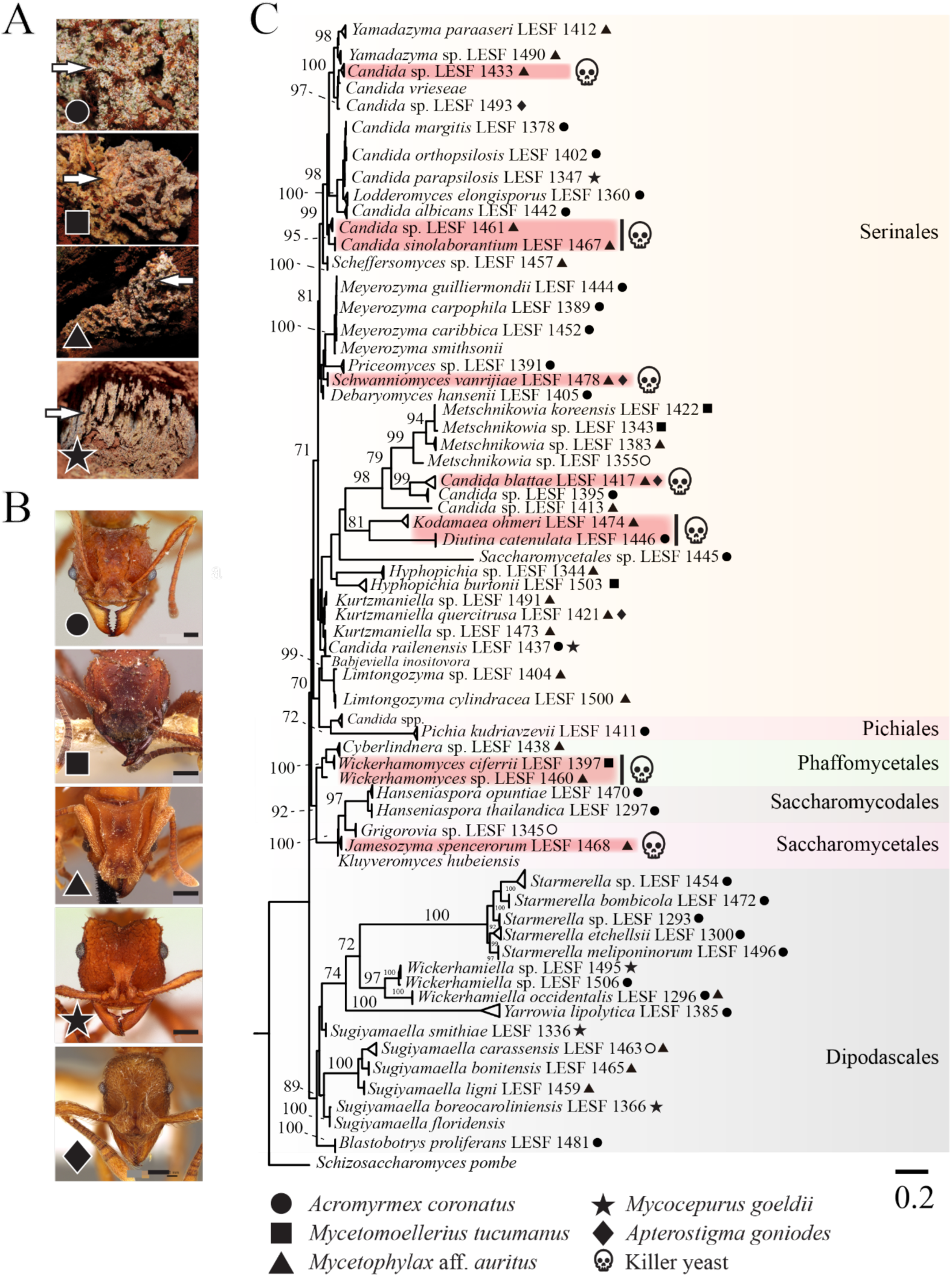
Fungus gardens of different attine ants harbor different killer yeasts. **A.** Images of fungus gardens of different ant species from which yeasts were isolated (arrowheads). The gardens of the leaf-cutting ant *Acromyrmex coronatus* (circle), the non-leaf-cutting ant *Mycetomoellerius tucumanus* (square), the lower attines *Mycetophylax* aff. *auritus* (triangle), and *Mycocepurus goeldii* (star). The fungus garden of *Apterostigma goniodes* is not represented. **B.** Representative images of attine ant species: *Acromyrmex coronatus* (CASENT0173791, April Nobile) (circle); *Mycetomoellerius tucumanus* (CASENT0909391, Will Ericson) (square); *Mycetophylax auritus* (CASENT0901666, Ryan Perry) (triangle), *Mycocepurus goeldii* (CASENT0173988, April Nobile) (star), and *Apterostigma goniodes* (CASENT0922040, Michele Esposito (diamond). The scale bar on all images represents 0.2 mm. Photos of ants were downloaded from “www.antweb.org.” **C**. Phylogenetic relationship of yeasts isolated from fungal gardens as inferred by maximum likelihood and the rRNA gene. Symbols in tree leaves indicate the isolation source of yeasts, as depicted in panels A and B. Red shading and skulls denote the species confirmed as killer yeasts. The phylogenetic clades were highlighted by order following the updated classification for the Saccharomycotina proposed by Groenewald et al., (2023) and www.mycobank.org. *Schizosaccharomyces pombe* was positioned as an outgroup. The number on the branches are ultrafast bootstrap values (values higher than 70 are shown), and the scale bar denotes the number of nucleotide substitutions per site.

From the 59 yeast species identified that were associated with attine ant gardens, ten species had isolates that were able to produce killer toxins (10 out of 59; 17% of killer yeasts) (Fig. 1C). Killer toxin production was scored by the absence of growth around a killer yeast or the staining of the lawn yeast with the redox indicator methylene blue (as an indicator of cell death) at pH 4.6. This pH was used as many previously described killer toxins are active in these conditions and it is close to the pH observed in fungus gardens where these yeasts were isolated (pH 5.1-5.4). The identified killer yeasts were effective at inhibiting strains that were either previously determined to be susceptible to killer toxins (Fredericks et al., 2021), human pathogenic yeasts, or yeasts randomly selected from fungus gardens and other sources (Fig. 2A and S1; File S1). From the attine ant gardens, *Candida blattae*, *Candida sinolaborantium*, *Candida* sp. (closest to *Candida temnochilae*), *Candida* sp. (closest to *Candida membranifaciens*), *Diutina catenulata*, *Jamesozyma spencerorum*, *Kodamaea ohmeri*, *Schwanniomyces vanrijiae*, *Wickerhamomyces ciferrii*, and *Wickerhamomyces* sp. (closest to *Wickerhamomyces rabaulensis*) were identified as killer yeasts. These yeasts had unique antifungal activities when compared to other previously identified *Saccharomyces* killer toxins (K1, K1L, K2, K21, K28, K45, K62, K74, and Klus), as has been observed in a previous screen for killer yeasts of *S. cerevisiae* (Crabtree et al., 2023). Killer yeasts were only absent from the fungus gardens of *M. goeldii*, but were recovered from the fungus gardens of *Acromyrmex coronatus* (7% killer yeasts, 6 out of 91 isolates), *Mycetomoellerius tucumanus* (46%, 6 out of 13), *Mycetophylax* aff. *auritus* (43%, 18 out of 42), and *Apterostigma goniodes* (22%, 2 out of 9) (Fig. 2B). Thirty-two out of 180 (18%) yeasts from attine ants were also capable of inhibiting the growth of susceptible yeasts, including human pathogenic yeasts such as *Candida albicans*, *Candida glabrata*, and *Candida auris* (Fig. S1). The attine ant-associated killer yeasts *J. spencerorum* LESF 1468 and *S. vanrijiae* LESF 1521 could only inhibit the growth of pathogenic yeasts (Fig. S1, File S1). Many killer yeasts from the same species had similar antifungal activities as demonstrated by their phenotypic clustering (see *Candida* sp. closest to *C. temnochilae*, *C. blattae*, *D. catenulata*, *S. vanrijiae*, *Wickerhamomyces* sp. closest to *W. rabaulensis*, and *W. ciferrii*) (Fig. 2A). Specifically, *Candida blattae* killer yeasts had similar antifungal activities even when isolated from the fungicultures of different ant species (compare *Candida blattae* isolates from *Mycetophylax* aff. *auritus* and *Apterostigma goniodes*) (Fig. 2A). However, there were isolates of *S. vanrijiae* killer yeasts from the fungiculture of *Mycetophylax* aff. *auritus* with strikingly different spectra of antifungal activities, which suggests that isolates of the same species were not always clonal. *Candida sinolaborantium* (strain LESF 1467) had the broadest range of antifungal activity, inhibiting 57% of all 69 lawn strains tested (Fig. 2A and S1, File S1). Comparing interactions between ant-associated killer yeasts and susceptible lawn strains from fungus gardens and other environmental sources found that growth inhibition was more prevalent for yeasts from different sources (Chi-squared test, p-value < 0.01) (Fig. 2C; Table S2). Specifically, only 1% (6 of 647) of interactions between ant-associated yeasts from the same type of fungiculture resulted in growth inhibition. In contrast, 4% (301 of 8,353) of interactions between ant-associated killer yeasts and yeasts from fungus gardens and other environmental sources isolation sources resulted in growth inhibition.

**FIG 2.**
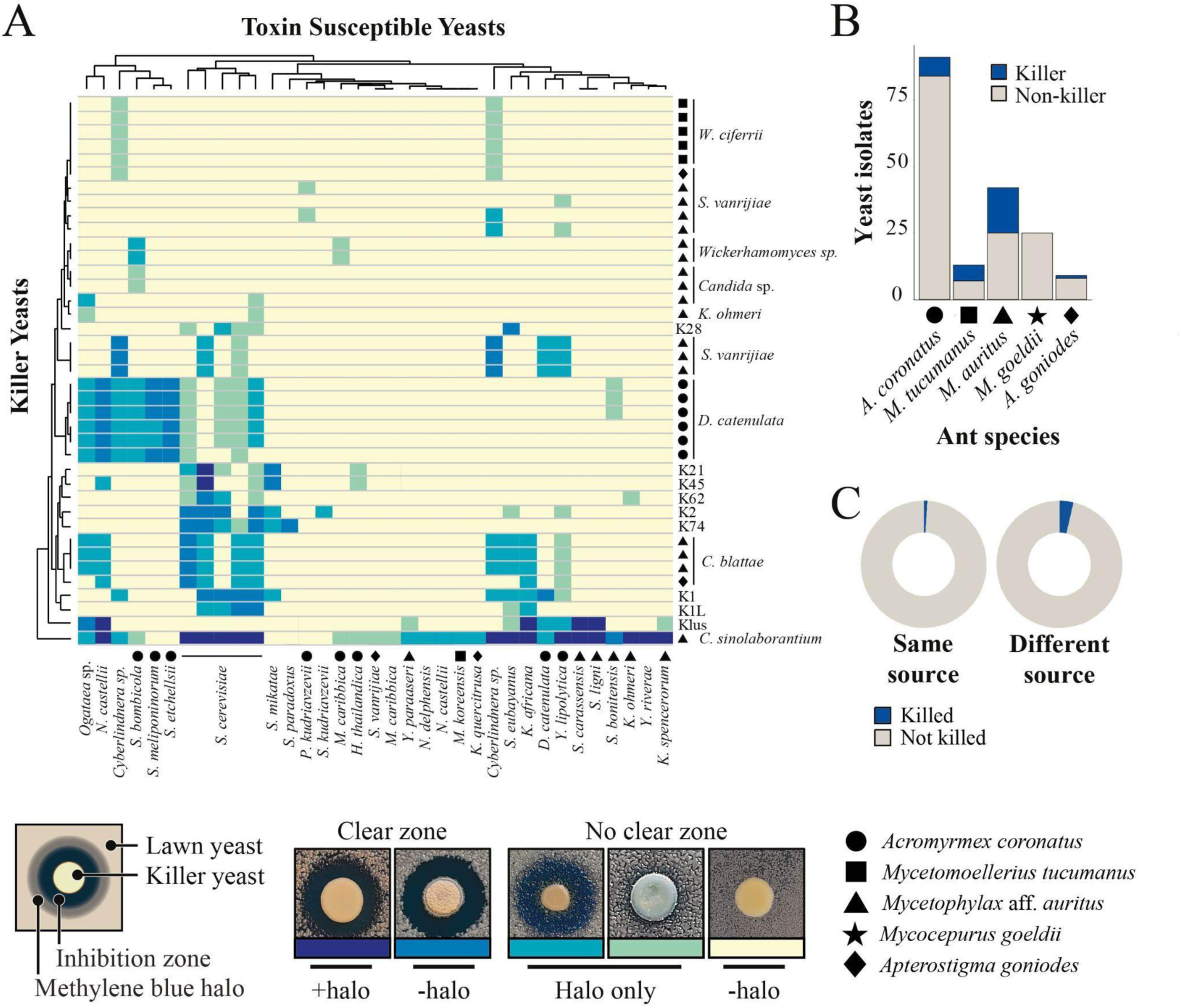
Killer yeasts from fungus gardens have unique spectra of antifungal activity. **A.** Heatmap with the interactions between canonical *Saccharomyces* killer yeasts (K1, K1L, K2, Klus, K21, K28, K45, K62, K74), killer yeasts associated with ants, and killer toxin susceptible strains. Origins of the ant-associated killer yeasts; *Acromyrmex coronatus* (circle); *Mycetomoellerius tucumanus* (square); *Mycetophylax* aff. *auritus* (triangle), *Apterostigma goniodes* (diamond). Killer toxin activity was qualitatively assessed based on the presence and size of growth inhibition zones and/or methylene blue staining around killer yeasts (representative images of these phenotypes are shown). Darker colors on the heatmap represent a more prominent killer phenotype, with yellow indicating no observable killer phenotype. Clusters on the dendrograms connecting individual killers or susceptible yeasts indicate similar susceptibilities to killer toxins or antifungal activities. **B.** Number of killer yeasts and non-killers associated with the different attine ant fungicultures. **C**. Number of killer toxin-positive and negative interactions between yeast from attine ant environment and susceptible strains (Table S1.), growth inhibition by killer yeasts is different among the same or different fungicultures and locations.

The ability to produce killer toxins is often associated with the presence of linear dsDNA plasmids or mycovirus-associated dsRNA satellites. We assayed all attine ant-associated killer yeasts for the presence of these genetic elements using solvent extraction of nucleic acids, cellulose chromatography, and gel electrophoresis (Crabtree et al., 2019). In particular, the use of cellulose chromatography selectively enriches dsRNAs derived from mycoviruses and satellites, reducing the false negative rate compared to solvent extraction alone and without the bias of reverse-transcriptase PCR (Crabtree et al., 2023). However, we did not find that any of the isolated attine ant-associated strains possess dsRNAs (i.e., viruses and satellites) or dsDNA cytoplasmic elements (i.e., plasmids). Double-stranded RNA viruses have been found in many different species of yeasts without satellite dsRNAs (Crucitti et al., 2021; Lee et al., 2022; Taylor et al., 2013); thus we broadened the survey to include all attine-associated yeasts but again did not identify any dsRNA viruses. This suggested that the killer toxins associated with yeasts from fungus gardens are genome-encoded and that dsRNA viruses, dsRNA satellites, and DNA plasmids were not observed in yeasts associated with attine ant species in the geographical area sampled.

### Characterization of the killer yeast *Candida sinolaborantium*

*C. sinolaborantium* was identified as a killer yeast with a broad antifungal spectrum that could inhibit 39 of the 69 yeast strains tested (Fig. 2 and S1). In contrast, the next most potent killer yeasts could only inhibit 11 strains tested. *C. sinolaborantium* was also able to cause large zones of growth inhibition in many different species of yeasts that were resistant to other killer yeasts (Fig. 2A). After several days of incubation, we observed that *K. ohmeri* produced a striking white halo between a zone of growth inhibition and methylene blue staining when challenged by the killer yeast *C. sinolaborantium* (Fig. 3A). This phenotype was unique to the pairing of *C. sinolaborantium* and *K. ohmeri*, and was distinct from the growth inhibition caused by the killer toxin K62 (the only other known killer toxin capable of inhibiting *K. ohmeri*) (Fig. 2A). The white halo also appeared to be raised from the surface of the agar plate (Fig. S2). When cells in this area (+halo) were observed under a microscope, they appeared to be elongated compared to cells of *K. ohmeri* outside of the halo (-halo) (Fig. 3A). The killer toxin (or toxins) secreted by *C. sinolaborantium* were precipitated by ethanol and remained active when incubated at room temperature, but were inactivated by heating to 60°C and 98°C (Fig. 3B). The inhibition of *K. ohmeri* by *C. sinolaborantium* was optimal at pH 5 (Fig. 3C, Table S3), with loss of killing at pH >5.5. The optimal temperature for killing *K. ohmeri* by *C. sinolaborantium* was 20°C. Using *C. castelli* as a sensitive lawn indicated a wider optimum temperature (17-25°C) and pH (3.5-5.0) (Fig. 3D, Table S3 and S4). In both cases the there was no killer activity at 35°C (Fig. 3C and 3D, Table S4). Further analysis of the white halo of *K. ohmeri* revealed an 8-fold increase in cell elongation in the white halos (+halo) when compared to untreated cultures of *K. ohmeri* (-halo) (Fig. 3E and Table S5). Cells of *K. ohmeri* from the white halo or untreated culture were seeded on agar to measure viability by colony-forming units. Cells isolated from the white halos showed a 2.1-fold reduction in colony-forming units compared to *K. ohmeri* isolated away from halos (Fig. 3F and Table S6). Ultrafiltration of culture media was used to determine the approximate molecular weight of the antifungal molecules produced by *C. sinolaborantium*. Antifungal activity was observed for fractions that enriched molecules >100 kDa and >30 kDa. Raw filtrates and concentrated ethanol-precipitated fractions were used to challenge a panel of yeasts previously shown to be susceptible to toxins produced by *C. sinolaborantium* (Fig. 2). The spent culture medium showed less antifungal activity compared to the precipitated medium, with the latter capable of inhibiting 21/29 susceptible yeasts that were inhibited in killer assays on agar (Fig. S3). Antifungal activities were similar before and after passage through a filter with a molecular weight cut-off (MWCO) of 100 kDa and were mostly retained in the MWCO 30 kDa filter. Only weak antifungal activity was detectable for the fraction that passed through the MWCO 30 kDa filter (Fig. S3). The appearance of white halos was observed using fractions captured by filters with MWCO 100 kDa and MWCO 30 kDa when tested against *K. ohmeri* but not with MWCO 5 kDa. Together, these data show that *C. sinolaborantium* is a killer yeast with a broad spectrum of antifungal activities, with similar temperature and pH optima of other previously described yeast killer toxins, but with the ability to induce cell elongation in *K. ohmeri*. Fractions from ultrafiltration are consistent with the antifungal activities of *C. sinolaborantium,* suggesting either the production of multiple chromosomally encoded killer toxins with similar antifungal properties or perhaps large oligomers or complexes of a single >30 kDa killer toxin.

**FIG 3.**
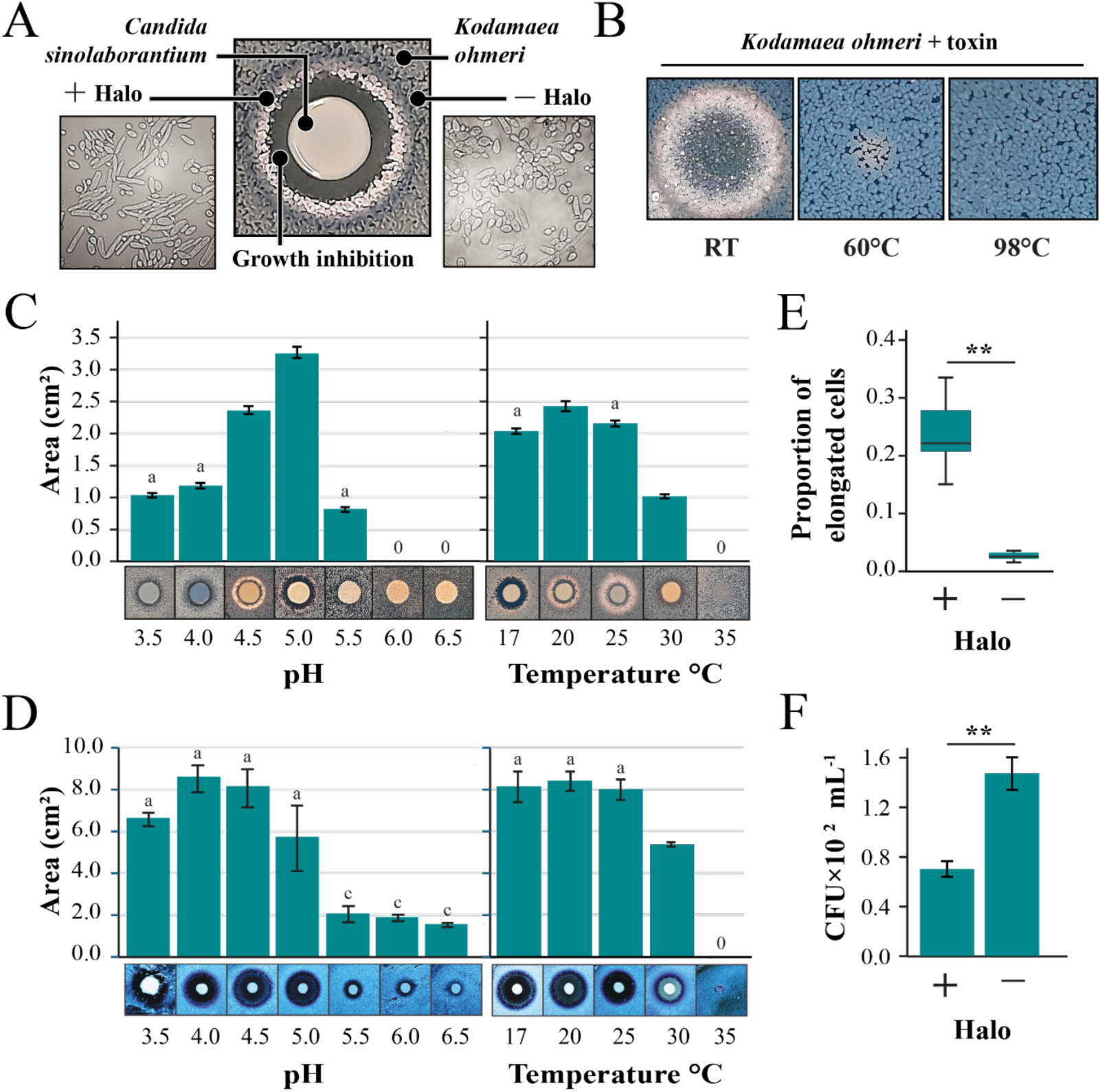
The antifungal activities of *Candida sinolaborantium*. **A**. Growth phenotypes associated with the co-culture of the killer yeast *C. sinolaborantium* with *K. ohmeri* at room temperature (RT) and pH 4.6. Cell morphology of *K. ohmeri* inside (+Halo) and outside (-Halo) the white halo around *C. sinolaborantium* was evaluated by microscopy. **B**. The antifungal activity of protein precipitates derived from the spent growth medium of *C. sinolaborantium* and the heat stability of these activities at 60°C and 98°C. Killer yeast activity of *C. sinolaborantium* under different pH and temperature conditions against **C.** *K. ohmeri* and *C. castelli*. For all pH and temperature data, the means were significantly heterogeneous (one-way anova, P<0.01). Means with the same letter are not significantly different from each other (Tukey–Kramer test, P>0.05). **E.** Proportion of elongated to normal cells of *K. ohmeri* inside (+Halo) and outside (-Halo) the white halo around *C. sinolaborantium*, as depicted in Fig. 3A. **F.** The same growth areas as shown in **E.** but colony forming units (CFU) were measured to indicate *K. ohmeri* cell survival. Asterisks indicate different means for the Welch Two Sample t-test (p-value < 0.01). All error bars are standard error (n>3).

### Ksino: a novel killer toxin produced by *C. sinolaborantium* with structural homology to the Klus killer toxin

To determine the gene or genes responsible for killer toxin production by *C. sinolaborantium*, the genome sequence of the yeast was determined (Table S7). The resulting 11.21 Mb assembly resulted in a BUSCO completeness (Complete and single-copy BUSCOs) of around 98% using the saccharomycetes_odb10 dataset (Seppey et al., 2019), with GC content of 49%, an N50 of 155,979 bp, and a total of 5,951 protein-coding genes (Table S7). To identify genome-encoded killer toxins in the *C. sinolaborantium* proteome, BLASTp was performed using a database of known killer toxins from yeasts (Table S8). This approach identified 12 toxin-like candidates, all with a sequence alignment length greater than 100 amino acids and an identity of at least 24% compared to any known killer toxin (Table S9). One *C. sinolaborantium* protein had a 28% identity and 42% similarity over an alignment length of 151 amino acids with the *S. cerevisiae* killer toxin Klus and was named Ksino (Genbank Accession: XAT76252) (Table S9) (Rodríguez-Cousiño et al. 2011).

The sequence similarity between Ksino and Klus prompted a more detailed investigation into whether these killer toxins share similar antifungal properties. Although we had previously failed to detect growth inhibition of *K. ohmeri* by *S. cerevisiae* killer toxin Klus (Fig. 3), we noted the appearance of white halos instead of a zone of growth inhibition or methylene blue staining after prolonged co-culture (Fig. 4A). The appearance of white halos was similar to those produced by *C. sinolaborantium* and was dependent on the presence of the Mlus satellite dsRNA that encodes the Klus killer toxin gene (Fig. 4A). Analysis of the cells within the white halo found a significant proportion of elongated cells, as previously observed upon exposure of *K. ohmeri* to Ksino (Fig. 4B). However, unlike the Ksino halos, Klus did not cause a significant loss of cell viability (Fig. 4C).

**FIG 4.**
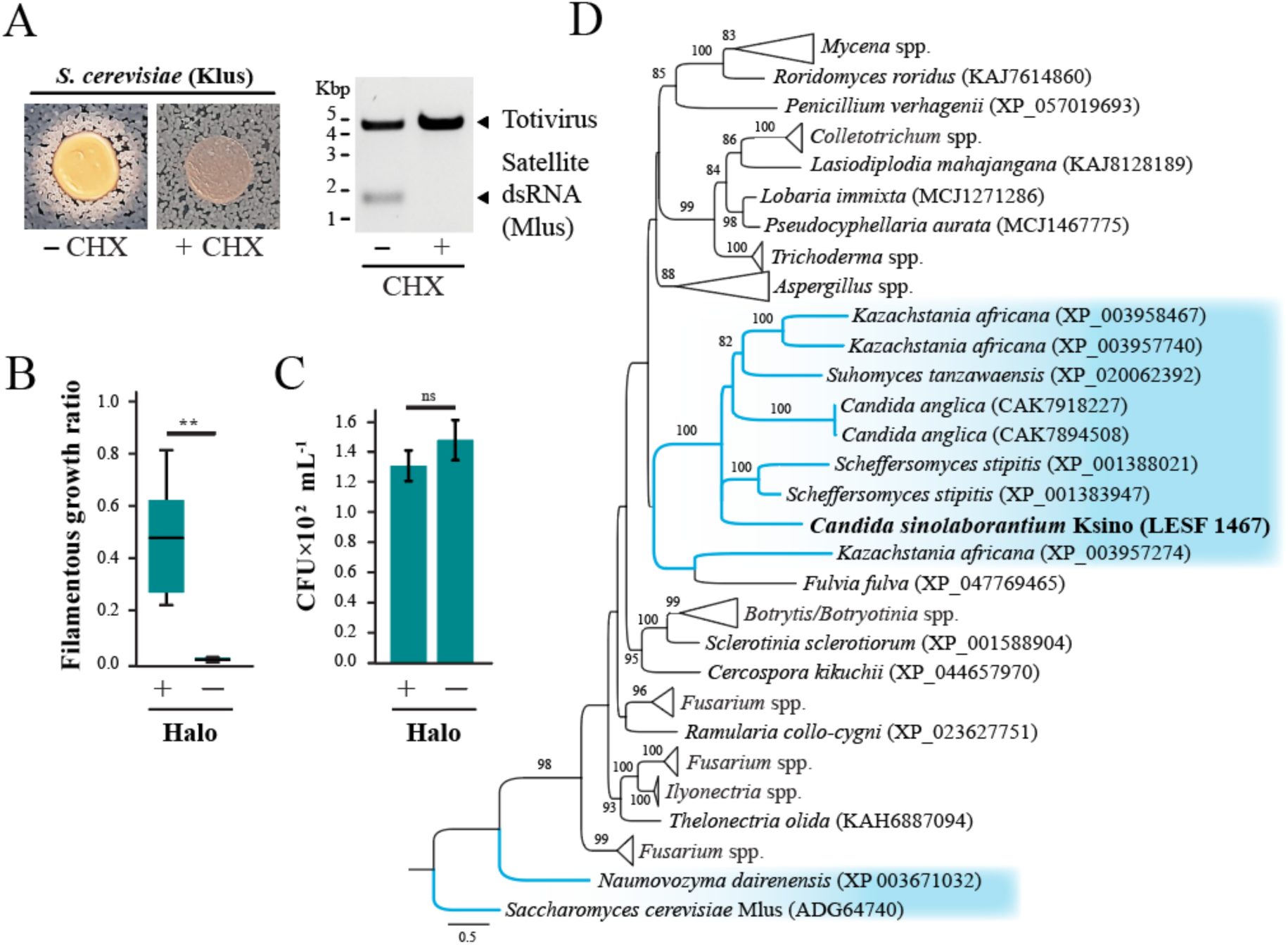
Ksino from *C. sinolaborantium* defines a novel family of killer toxins. **A.** The phenotypes associated with the co-culture of the Klus-encoding killer yeast *S. cerevisiae* strain DSM-70459 with *K. ohmeri* (+Mlus). Treatment with cycloheximide (CHX) causes the loss of the Mlus satellite (-Mlus) and the loss of the white halo phenotype. The ratio of **B.** elongated and **C.** viable *K. ohmeri* cells in the white halo around Klus-producing *S. cerevisiae* (+Halo) and in axenic culture (-Halo). Asterisks indicate different means for Welch Two Sample t-test (p-value <0.01), ns = not significant. **D.** A maximum likelihood phylogeny of Ksino homologs identified in diverse fungi. The Ksino from *C. sinolaborantium* is highlighted in bold, and all Saccharomycotina yeasts are highlighted in blue. The dsRNA-encoded killer toxin Klus is represented as an outgroup. Bootstrap values higher than 70 are shown. All error bars are standard error (n>3).

To identify other possible Ksino-homologous proteins in different organisms, we performed a BLASTp search across the NCBI database, resulting in 84 protein sequences from various Ascomycota and Basidiomycota fungi. Ksino homologs were found in *Saccharomycetes, Pichiomycetes* yeasts, and other filamentous fungi with identity and alignment lengths higher than 30% and 109 amino acids, respectively (Fig. 4D and Table S11). Phylogenetic analysis of homologs, with Klus as an outgroup, indicates that proteins with the highest identity of Ksino (35-42% identity) were all uncharacterized genes of the Saccharomycotina yeasts from the lineages of *Debaryomycetaceae* and the *Saccharomycetaceae* (Fig. 4D). Saccharomycotina species with Ksino-like genes included *Candida anglica*, *Kazachstania africana*, *Scheffersomyces stipitis*, and *Suhomyces tanzawaensis*, with both *K. africana* and *S. stipitis* (Basionym: *Pichia stipitis*) being previously reported as killer yeasts (Antunes and Aguiar, 2012; Fredericks et al. 2021). Filamentous fungi encode the majority of Ksino homologs (75 out of 84), including species belonging to the genera *Fusarium*, *Mycena*, and *Aspergillus* (Fig. 4D and Table S11), known as plant and human pathogens (*Fusarium* spp.), saprophytic and biotrophic plant associated fungi (*Mycena* spp.), and fungi species that are ubiquitous in the environment (*Aspergillus* spp.) (Summerell et al. 2010, Harder et al. 2023; Thoen et al. 2020; Houbraken et al. 2020; Jurjevic et al. 2012; Peterson et al. 2001).

The Ksino open reading frame was determined to be 202 amino acids long, comparable to Klus at 242 amino acids. Linear representation of the secondary structure revealed a similar pattern of alpha helices and beta sheets (Fig. 5A; Fig. S5). There was also evidence of potential post-translational modification sites, including N-terminal signal sequence cleavage sites predicted by psipred (Buchan et al., 2013) and dibasic motifs (i.e., KR) in the unstructured region before helix 1 (Fig. S5). Dibasic motifs in killer toxins are known to be proteolytically cleaved by the Golgi-specific protease Kex2 (Julius et al., 1984). As Ksino has no sequence homology to a protein of known structure, AlphaFold2 was used to generate tertiary structure models of Ksino and Klus (Jumper et al., 2021). Ksino and Klus had global LDDT scores of 63.6 and 65.1, respectively, and per residue, LDDT was greater than 80 in regions of secondary structure for both proteins (Fig. 5A and S5). Molecular dynamics simulation improved the bond angles of the alphafold model to Ramachandran favored regions in models of Ksino (77% to 91%) and Klus (73% to 89%) (Table S10, Fig. S6). Ksino and Klus simulations reached a stable RMSD of around 1.6 nm after approximately 20 ns of simulation (Fig. S6). Molprobity scores, analogous to structure resolution, were determined to be 1.54 for Ksino and 1.86 for Klus (Studer et al., 2019). Overlay of the final energy-minimized structures of Ksino with Klus compared to the original AlphaFold2 models yielded an RMSD of 2.6 Å (Fig. 5B). The predicted organization of the Ksino structure consists of a discontinuous five-stranded antiparallel beta-sheet packed against a pair of antiparallel alpha helices (Fig. 5B and S5). The tertiary structure of Klus follows the same overall organization but with one less beta-turn. Ksino has eight cysteine residues, compared to the seven in Klus (Fig. S5). The predicted structures have three intramolecular disulfide bonds. In Ksino, C105-C127 is positioned to crosslink the C-terminus of helix 1 to the beta-sheet 2, similar to C141-C162 in Klus (Fig. S5). The remaining predicted disulfide bonds are unique to both structural models (Fig. S5). These simulations collectively suggest that the Klus and Ksino proteins share significant structural homology despite having low amino acid identity.

**Figure 5.**
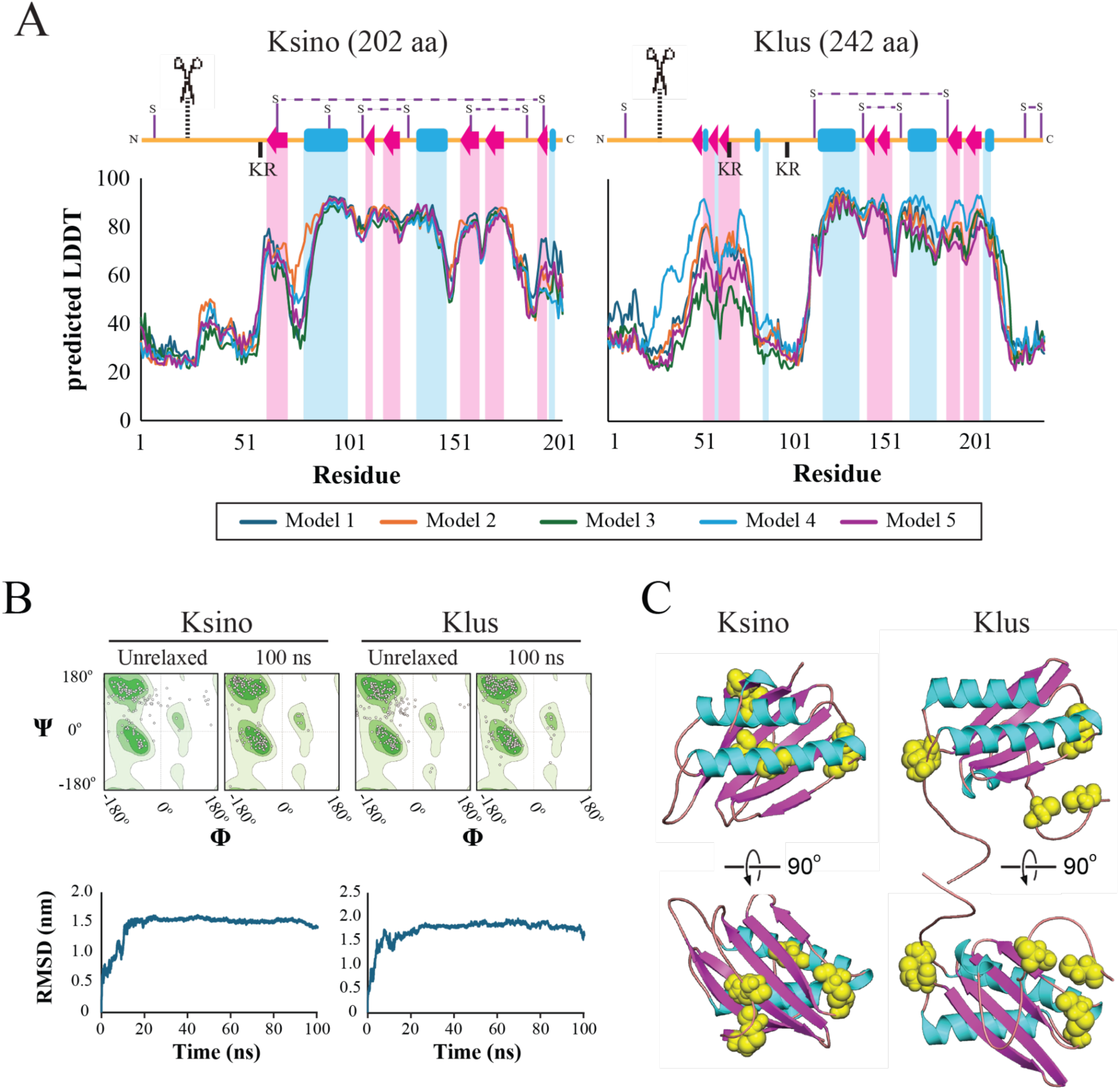
Molecular modeling of Ksino indicates structural homology to Klus. pLDDT confidence per residue and predicted disulfide bond arrangement of AlphaFold2 models of Ksino and Klus. A. Secondary structure representations aligned with per-residue pLDDT (predicted local distance difference test) scores from AlphaFold2. Alpha helices are represented as blue rectangles, beta sheets as red arrows, and unstructured regions as orange lines. Predicted disulfide linkages are indicated as horizontal dashed lines. Scissors with vertical dashed lines represent predicted signal sequence cleavage sites. B. Ramachandran plots of general residues (non-proline/glycine) generated by SWISS structure assessment tool demonstrating improved Ramachandran favored after performing 100 ns simulation. The unrelaxed structure represents AlphaFold2’s raw output (Ramachandran et al., 1963). **B.** Full protein RMSD over 100 ns molecular dynamics simulation. GROMACS was used to generate alignments of each snapshot back to the structure at 0 ns. C. Ksino and Klus AlphaFold2 models with cysteine residues highlighted as yellow spheres to illustrate predicted disulfide linkage arrangement. The cartoon helices and beta sheets are colored to match panel A.

The molecular weight of Ksino was predicted to be 21.5 kDa, which appeared to be smaller than the predicted molecular weight determined by ultrafiltration (Figure S3). To determine if Ksino was was expressed by *C. sinolaborantium*, total RNA was extracted from laboratory cultured yeast cells in order to assay for the presence of Ksion transcript by reverse transcriptase PCR (RT-PCR). This approach was successful in detecting the presence of Ksino RNA transcripts (Figure S4). To confirm that Ksino is an active killer toxin, the gene was amplified by PCR and cloned into an *S. cerevisiae* expression vector under the control of a galactose-inducible promoter. *C. sinolaborantium* belongs to the Serinales order and uses the CUG codon to encode serine (Groenewald et al 2023; Krassowski et al., 2018). Ksino has one CUG codon that will introduce a leucine at position 186 when expressed by *S. cerevisiae*. This mutation is predicted to have a small destabilizing effect on folding compared to the wild type based on FoldX 5.0 and molecular dynamics simulation (estimated ΔΔG_Folding_ = 1.0 kcal/mol) (Fig. 5). Therefore, residue 186 was recoded for serine in *S. cerevisiae* (L186S). When the strains were grown on dextrose or galactose containing media, no growth inhibition of *K. ohmeri* was observed during co-culture with a laboratory strain of *S. cerevisiae* transformed with the wild type or L186S expression vectors (Fig. SXX). However, challenging the strain *C. castelli* Y-17070 with Ksino wild type or L186S on galactose triggered large zones of inhibition and methylene blue staining, indicating that Ksino is a novel killer toxin (Figure 6A). Growth of *S. cerevisiae* under different temperatures and pH identified that Ksino has an optimum antifungal activity at pH4-4.5 (Figure 6B and Table S13) and 20-25°C (Figure 6C and Table S14), which was narrower than the killer phenotype of *C. sinolaborantium* (Figure 3). The inability of Ksino expressed by *S. cerevisiae* to prevent the growth of *K. ohmeri* and differences in the optimal conditions for antifungal activity on *C. castelli*, indicated that Ksino may be one of many antifungal molecules produced by *C. sinolaborantium*, or that expression of Ksino by *S. cerevisiae* alters the toxins spectrum of activity (Fig. 4F and S8).

**Figure 6.**
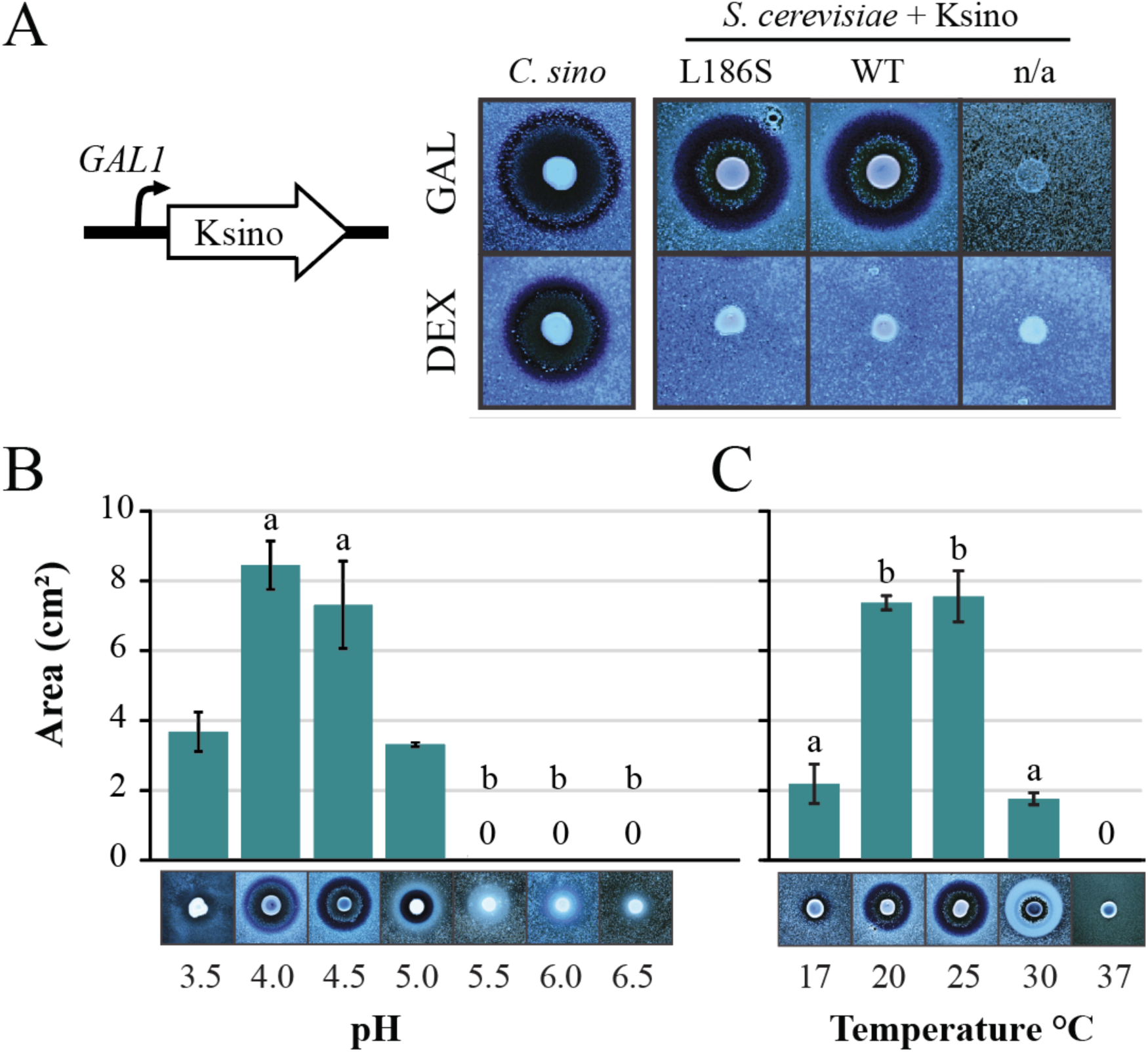
Ksino is an antifungal killer toxin when expressed by *S. cerevisiae*. **A**. Recombinant expression of Ksino (L186S) by *S. cerevisiae* using galactose induction with a lawn of *C. castelli* Y-17070 (pH 4.5). Dextrose was used as a non-induced control. n/a indicates an empty vector negative expression control. Zones of growth inhibition were compared to the killer phenotype of *C. sinolaborantium*. B. Killer yeast activity under different **B**. pH and **C**. temperature conditions as measured by zone of growth inhibition and methylene blue staining. Representative images of the killer phenotype are displayed under each histogram. The means were significantly heterogeneous (one-way anova, P<0.01) for both the pH and temperatures tested. Means with the same letter are not significantly different from each other (Tukey–Kramer test, P>0.05). All error bars are standard error (n=3).

## 3. Discussion

The large number of killer yeasts in fungicultures suggests they might play a role in attine ant fungus gardens by suppressing fungal competitors or allelopathy effects on complex dynamics of microbial interactions. Different studies have revealed that yeasts are abundant in attine ant environments that likely represent differences in the foraging composition and substrate preparation behaviors observed among the different fungicultures (Bizarria Jr. et al., 2022, de Fine Licht and Boomsma, 2010; Ronque et al., 2019). Yeasts from fungus gardens of the leaf-cutting ant *Atta texana* were observed to suppress the growth of fungal garden contaminants, especially *Escovopsis* and *Syncephalastrum* (Rodrigues et al. 2009). These antifungal properties were hypothesized to contribute to fungal garden “immunity,” whereby the resident fungi within a fungiculture prevent invasion by other fungal species. This would require interplay between the cleaning behaviors of attine ants in their gardens to remove unwanted fungi and other microbial interactions within attine ant colonies, i.e., allelopathy. Our study indicates that fungus gardens of attine ants harbor killer yeasts with similar prevalence to other surveys (Philliskirk and Young, 1975; Pieczynska et al., 2013; Starmer et al., 1987; Stumm et al., 1977; Trindade et al. 2002; Wojcik and Kordowska-Wiater, 2015; Abranches et al. 1997, Abranches et al. 1998). This expands the previous findings regarding the prevalence of killer yeasts in laboratory-reared attine ant colonies of *Atta sexdens* (Carreiro et al., 2002). Moreover, the prevalence of killer yeasts among the different ant species, such as *Mycetophylax* aff. *auritus* (43%, eight nests sampled) and *Mycetomoellerius tucumanus* (46%, four nests sampled) were considerably higher than in other surveyed fungicultures.

Killer yeasts from attine ant colonies were more likely to inhibit the growth of susceptible strains from different fungicultures. This supports previous observations of killer and non-killer yeast interactions whereby local selective pressure for killer toxin resistance allows the coexistence of killer and non-killer yeasts in a given niche (Starmer et al. 1987; Ganter and Starmer, 1992, Abranches et al. 1997, Trindade et al. 2002). The susceptibility of yeast populations that have never been exposed to a specific killer toxin would greatly depend on the standing genetic variation (perhaps in the cell wall and membrane toxin receptors) among different lineages of yeasts dictated by the population’s evolutionary history. Our findings support the hypothesis that yeasts contribute to fungal garden “immunity” since killer yeasts are expected to prevent the invasion of other fungal species into the fungus garden. The coexistence of killer and non-killer yeast species could be an outcome of microhabitats in the spatially structured environment of the fungus garden, which may harbor different populations of yeasts and allow the colonization by killer yeasts (Giometto et al. 2021, Wloch-Salamon et al. 2008).

Saccharomycotina yeast species isolated in attine ant gardens have been previously found to secrete killer toxins. Among the yeast species, the ability to produce toxins was observed for *Diutina catenulata* (Basionym: *Candida catenulata*; Stopiglia et al., 2014; Fuentefria et al., 2007), *Kodamaea ohmeri* (Synonym: *Pichia ohmeri*; Fuentefria et al., 2006, Fuentefria et al., 2008; Coelho et al., 2009), *Schwanniomyces vanrijiae* (Madbouly et al., 2020), and *Wickerhamomyces ciferrii* (Basionym: *Hansenula ciferrii*; Nomoto et al., 1984). The exception was *Jamesozyma spencerorum* (Synonym: *Kazachstania spencerorum* nom. inv.), which, to our knowledge, is the first report of a killer phenotype displayed by this species, although toxin production has been previously reported in the genus, specifically *K. africana*, *K. exigua* and *K. unispora* (Perez et al., 2016; Stopiglia et al., 2014; Fredericks et al., 2021). These yeast species have also been identified in substrates that are commonly foraged or associated with attine ants, including plants (*Schwanniomyces vanrijiae*, Madbouly et al., 2020; *Kodamaea ohmeri*, Fuentefria et al., 2006), insects and their environments (*Candida blattae*, Nguyen et al., 2007; *Candida sinolaborantium*, Suh et al., 2005; Guamán-Burneo et al., 2015; *Schwanniomyces vanrijiae*, Maksimova et al., 2016; *Kodamaea ohmeri*, Benda et al., 2008; Amos et al., 2019; Suh and Blackwell, 2005), and soil (*Jamesozyma spencerorum*, Synonym : *Kazachstania spencerorum* nom. inv.; Vaughan-Martini et al., 1995; Kurtzman, 2003; Kurtzman and Robnett, 2003; Liu et al., 2024).

Importantly, killer yeast species appear to be unique between colonies of the same ant species (Bizarria Jr. et al., 2023). This observed diversity could be due to the foraging behavior of ants, whereby sampling of the surrounding environment would introduce different fungal species to the fungiculture of the same species of ants in different geographical locations (Bizarria Jr. et al., 2023). However, despite the potential role of killer yeasts in garden immunity, whether the occurrence of killer yeasts in a fungiculture is acquired by chance, inherited from the environment, or inherited during garden propagation remains unknown (Bizarria Jr. et al., 2022). Niche construction by the fungal cultivar species could also play a role in yeast diversity in the fungus garden environment or by other features such as the pH of the fungus garden. Interestingly, the pH observed in the different fungus gardens (pH 5.1-5.4) is consistent with the optimal antifungal activities of *C. sinolaborantium* (pH 3.5-5.0) and Ksino (pH 4.0-4.5). However, the association between killer yeasts and fungicultures still requires robust empirical evidence to reject the stochastic nature of killer yeast colonization. Future yeast isolation will provide new directions to clarify whether killer yeast occurrence is modulated by ant foraging choice or fungus garden features such as pH and fungal cultivar species.

Our survey of the encoding elements of killer toxins indicated that the 180 isolated strains of Saccharomycotina yeasts were devoid of cytoplasmic killer toxin-encoding elements such as dsRNA satellites and dsDNA linear plasmids. The frequency of dsRNA elements associated with *Saccharomyces* yeasts has been observed to range from 10% to 51% in wild and domesticated strains (Pieczynska et al., 2013; Crabtree et al., 2023). It is expected that more of these genetic elements will be found in other Saccharomycotina lineages, given their discovery in non-*Saccharomyces* yeasts and the frequency of genome-integrated molecular fossils of plasmid and virus sequences in different lineages (Frank and Wolfe, 2009; Liu et al., 2010; Satwika et al., 2012; Myers et al., 2020; Villan Larios, et al. 2023; Taylor and Bruenn, 2009, Lee et al., 2022). Most of the yeast isolated from fungicultures belong to the Serinales, where the CUG codon is translated into serine instead of leucine in the universal genetic code (Groenewald et al 2023). The rewiring of the fungal genetic code has been suggested to prevent the invasion of viruses and plasmids by stopping the faithful translation of viral proteins (Krassowski et al., 2018). Alternatively, RNAi might also contribute to limiting viral infections in these yeasts, even though, in some cases, yeast lineages maintain viruses that suppress RNAi (Drinnenberg et al., 2009; Drinnenberg et al., 2011; Lee et al., 2022; Segers et al., 2006; Segers et al., 2007; Rodriguez Coy et al., 2022). RNAi and genome recoding could be possible reasons for the absence of cytoplasmic elements in the yeasts associated with attine ant fungiculture. However, this still requires a more in-depth investigation to exclude the stochastic nature of virus acquisition by fungi and sampling biases.

The lack of extrachromosomal elements in killer yeasts associated with attine ant fungiculture suggests that killer toxin genes are genome-encoded. Our survey indicates that the killer toxins from different Saccharomycotina lineages associated with fungus gardens differ in their spectrum of antifungal activity from canonical *Saccharomyces* killer toxins (K1, K2, K28, etc.). In particular, toxins secreted by *Candida sinolaborantium* had the broadest activity, including toward infectious human pathogens (e.g., *Candida albicans*, *C. glabrata*, and *C. auris*). A unique response to the killer activity of *C. sinolaborantium* was also observed in *K. ohmeri*, where cells exposed to the killer yeast grew as filaments. Usually, dimorphism in yeasts has been demonstrated to be triggered by nitrogen starvation (Gimeno et al., 1992), different carbon sources, and temperature (Rupert and Rusche, 2022). However, to our knowledge, this is the first case of this phenotype associated with exposure to a killer yeast. The broad spectrum antifungal activities of *C. sinolaborantium* and the ability to fractionate antifungal activities by ultrafiltration indicates the likely presence of multiple unidentified chromosomally encoded toxins. After determining the genomic sequence of *C. sinolaborantium*, the new killer toxin Ksino was discovered due to its sequence homology to the canonical *Saccharomyces* killer toxin Klus. However, the killer activity of *C. sinolaborantium* was not fully reproduced when Ksino was heterologously expressed by *S. cerevisiae* and was unable to inhibit or cause filamentouse growth in *K. ohmeri*. Instead robust growth inhibition was only observed after challenging *C. castelli.* These data also supports the expression of multiple killer toxins by *C. sinolaborantium* or that ectopic heterologous expression by *S. cerevisiae* is inadequate due to potential deficiencies in extracellular export, maturation, or Ksino’s innate toxicity.

Many proteins from different Ascomycete and Basidiomycete fungi lineages were identified in our search for Ksino-like proteins, including filamentous fungi and yeast (from Saccharomycetes and Pichiomycetes). The Ksino-like proteins from yeasts of the Saccharomycotina subphylum (*Candida anglica*, *Kazachstania africana*, *Scheffersomyces stipitis*, and *Suhomyces tanzawaensis*) share a closer evolutionary origin to Ksino and therefore might represent bonafide killer toxins. Despite our approach to identifying killer toxins being limited by primary sequence similarity to known toxins, it has successfully identified Ksino. This combination of genome sequencing, sequence homology searches, and structural modeling could be used more widely on the many killer yeasts that are devoid of cytoplasmic extrachromosomal elements to identify new killer toxins (Boynton, 2019; Crabtree et al. 2023).

In summary, this study demonstrates that (i) Fungus gardens of attine ants can harbor killer yeasts, an underappreciated environment for killer yeasts. (ii) DsRNA and dsDNA extrachromosomal elements are lacking from yeasts associated with attine ant fungicultures. (iii) Genome mining can discover new chromosomally encoded killer toxins, specifically Ksino secreted by *Candida sinolaborantium*. (iv) Ksino has sequence and structural homology to the *Saccharomyces* killer toxin Klus and is a member of a new family of killer toxins in Ascomycete and Basidiomycete fungi.

## Data Availability

Data supporting the results in the paper are available in the Supplementary Material. The GenBank accession number for the Open reading frames of the Ksino killer toxin of *Candida sinolaborantium* LESF 1467 is PP790515. The sequenced genome of *Candida sinolaborantium* can be accessed through BioProject PRJNA916362, BioSample SAMN32422847, Assembly JAQRFW000000000. The raw sequence reads are also available through the NCBI Sequence Read Archive (SRR23081998). Gene prediction and annotation are provided in https://figshare.com/s/1f31cd262fc9de969d38 (10.6084/m9.figshare.25895278).

## Acknowledgments

We would like to thank Conselho de Gestão do Patrimônio Genético (CGEn) for providing permits for access to genetic heritage: #AE012D8, #A877F11 and #AA39A6D. We also thank Amanda Viana and Luciana Simão Carneiro for preparing and preserving yeast cultures. We also would like to thank the ARS Culture Collection (NRRL), Westerdijk Fungal Biodiversity Institute (WI-KNAW), and Dr. Roland Klassen for providing yeast cultures used in this study. We also thank Dr. Primrose Boynton for the critical reading of this manuscript. Photos of ants in Fig. 1 were accessed from “www.antweb.org”: CASENT0173791 *Acromyrmex coronatus* (Photographer: April Nobile); CASENT0909391 *Mycetomoellerius tucumanus* (Photographer: Will Ericson); CASENT0901666 *Mycetophylax auritus* (Photographer: Ryan Perry); CASENT0173988 *Mycocepurus goeldii* (Photographer: April Nobile); CASENT0922040 *Apterostigma goniodes* (Photographer: Michele Esposito).

## Authors’ contributions

RB wrote the manuscript supervised by PAR and with inputs from AR, JWC, and MY; RB conducted cross-interaction killer assays and dsRNA extractions. JWC performed protein structure predictions and analysis. TJB conducted cross-interactions with pathogenic yeasts. RACS performed genome assemblage and annotation. SAC performed killer assays for Ksino characterization. SAC performed microscopic evaluations and killer assays. RB and RTT performed dsDNA extractions. NF and MJD performed genome sequencing. PAR and AR supervised and obtained funding for the project. AR is responsible for permit applications. All authors discussed and reviewed the manuscript and approved the final version.

## Supplementary figures

**FIG S1.**
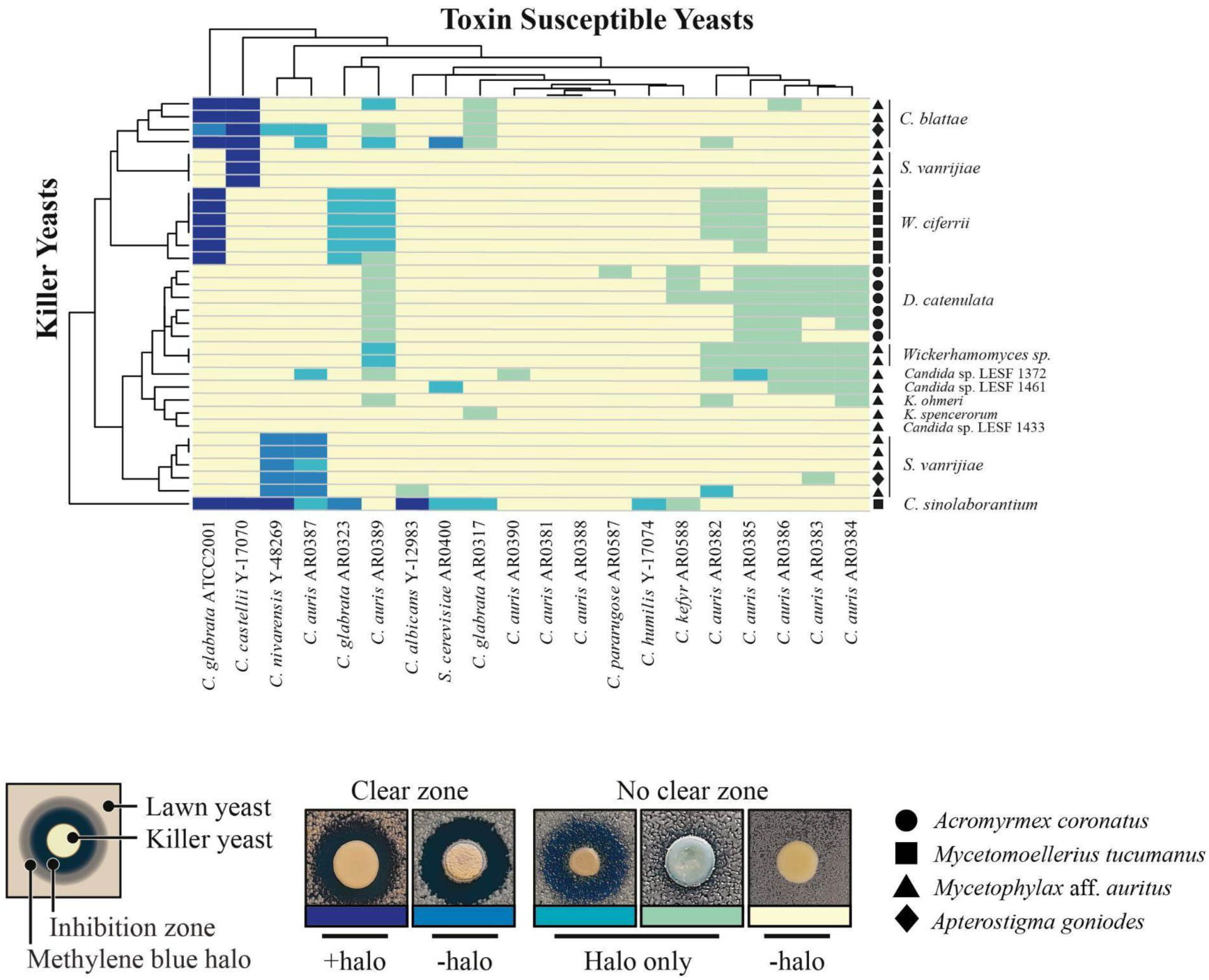
Killer yeasts from attine ant gardens can inhibit the growth of opportunistic human pathogens. A heatmap showing interactions between killer yeasts associated with attine ants and opportunistic human pathogens. Origins of the ant-associated killer yeasts; *Acromyrmex coronatus* (circle); *Mycetomoellerius tucumanus* (square); *Mycetophylax* aff. *auritus* (triangle), *Apterostigma goniodes* (diamond). Killer toxin activity was qualitatively assessed based on the presence and size of growth inhibition zones and/or methylene blue staining around killer yeasts as diagrammed. Darker colors on the heatmap represent a more prominent killer phenotype, with yellow indicating no observable killer phenotype. Clusters on the dendrograms connecting individual killers or susceptible yeasts indicate similar susceptibilities to killer toxins or antifungal activities. Darker colors on the cluster diagram represent a more prominent killer phenotype, with yellow indicating no observable killer phenotype.

**FIG S2.**
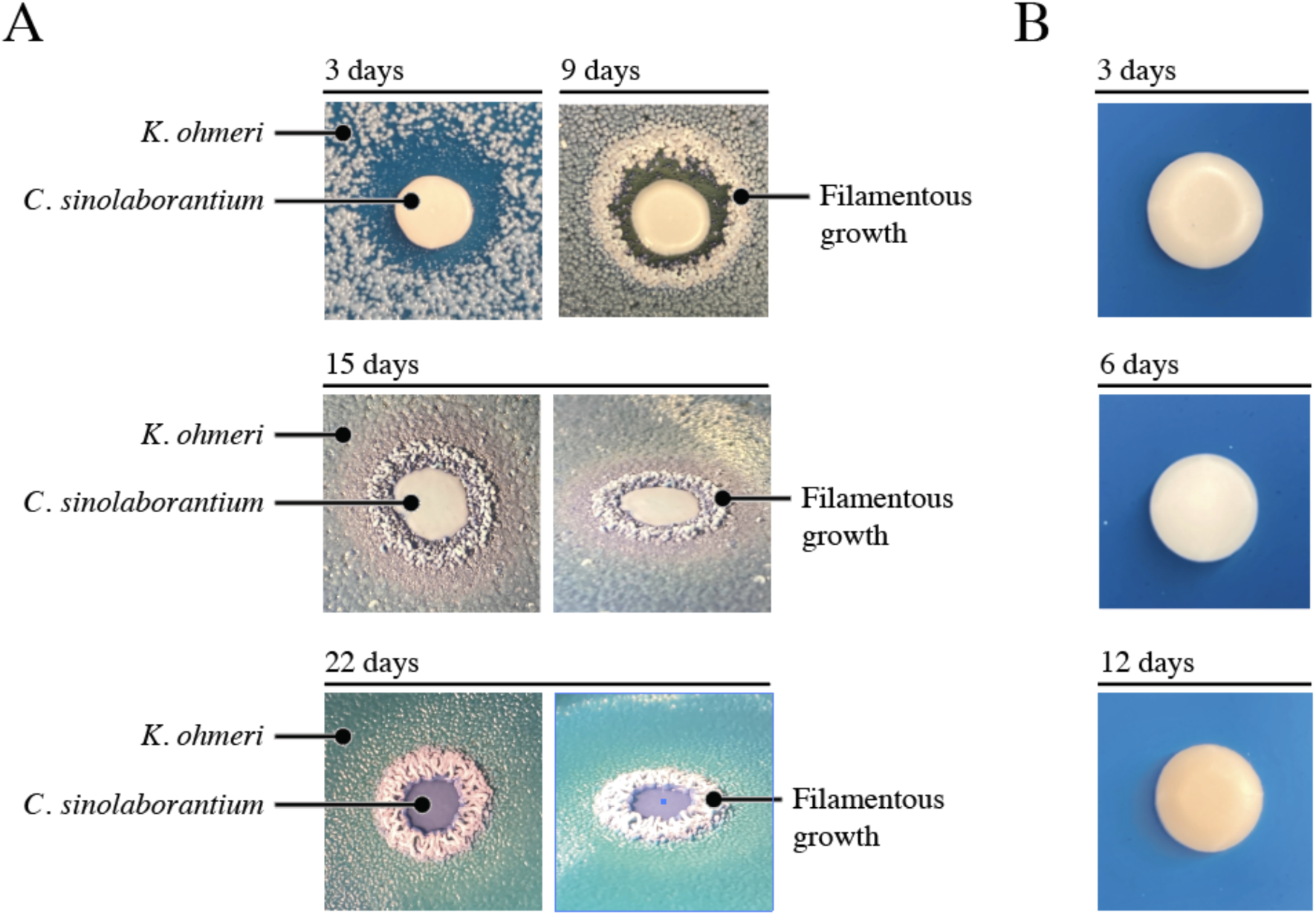
Elongated growth of *K. ohmeri* after coculture with the killer yeast *C. sinolaborantium*. A. Representative images showing the interaction between *K. ohmeri* and the killer yeast *C. sinolaborantium* on YPD agar. After extended incubation times at ∼22°C (15 - 22 days), vertical growth from the surface of the agar was observed. B. Elongated growth phenotypes are not observed when *C. sinolaborantium* is grown in isolation on agar.

**FIG S3.**
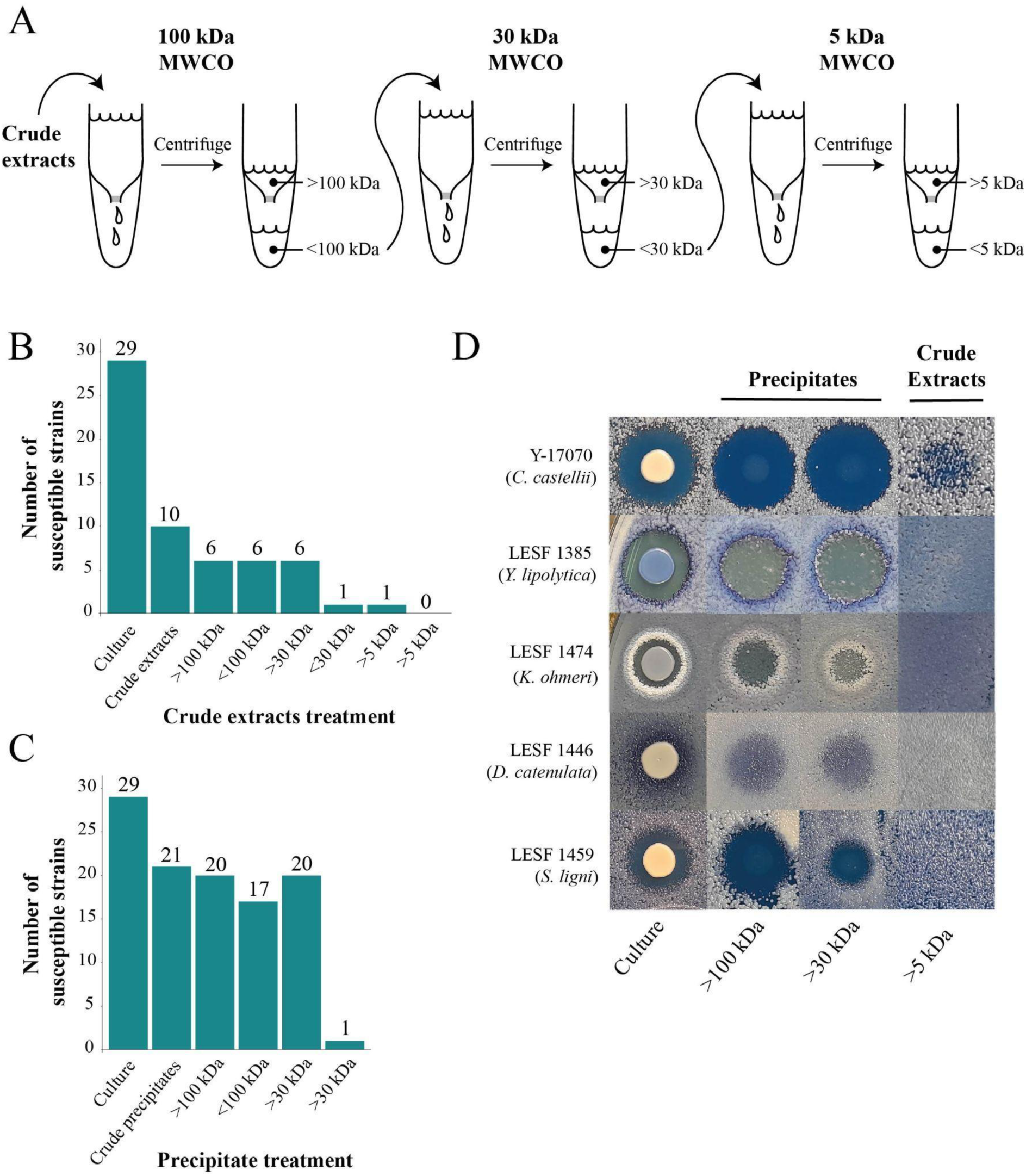
Antifungal activities exhibited by fractionated media after growth of *Candida sinolaborantium*. **A.** The workflow demonstrates the ultrafiltration of YPD-spent growth media to obtain fractions of antifungal molecules. Crude extracts were processed by ultrafiltration with molecular weight cut-off (MWCO) filters of 100 kDa, 30 kDa, and 5 kDa. The resulting protein preparations were used to challenge susceptible yeasts before (>) and after (<) ultrafiltration. Fractions were used **B.** before or **C.** after ethanol-precipitation **D.** Phenotypic examples of antifungal activity detectable in different fractions against representative susceptible yeast species.

**FIG S4.**
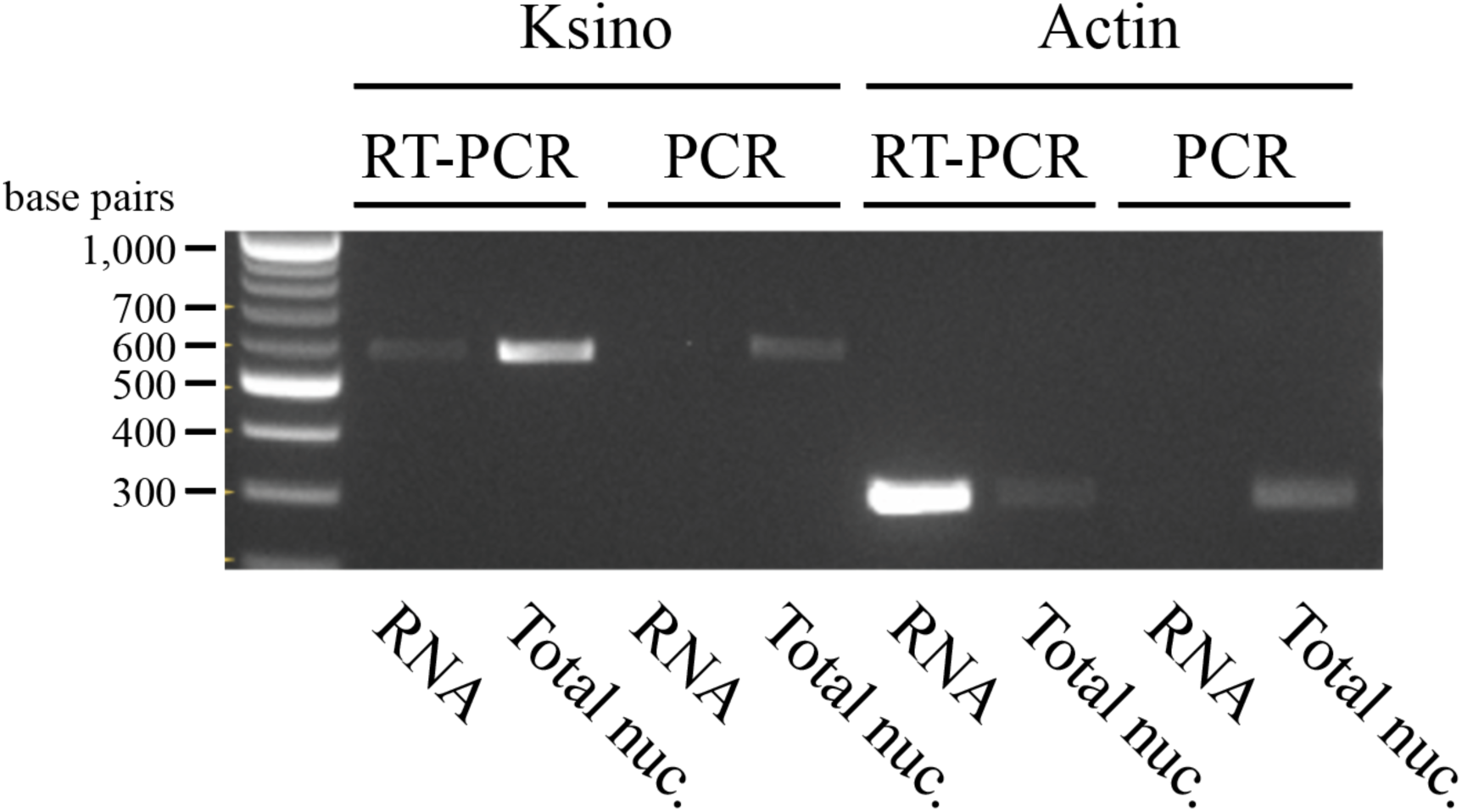
Detection of Ksino expression by RT-PCR. XXXXXX

**FIG S5.**
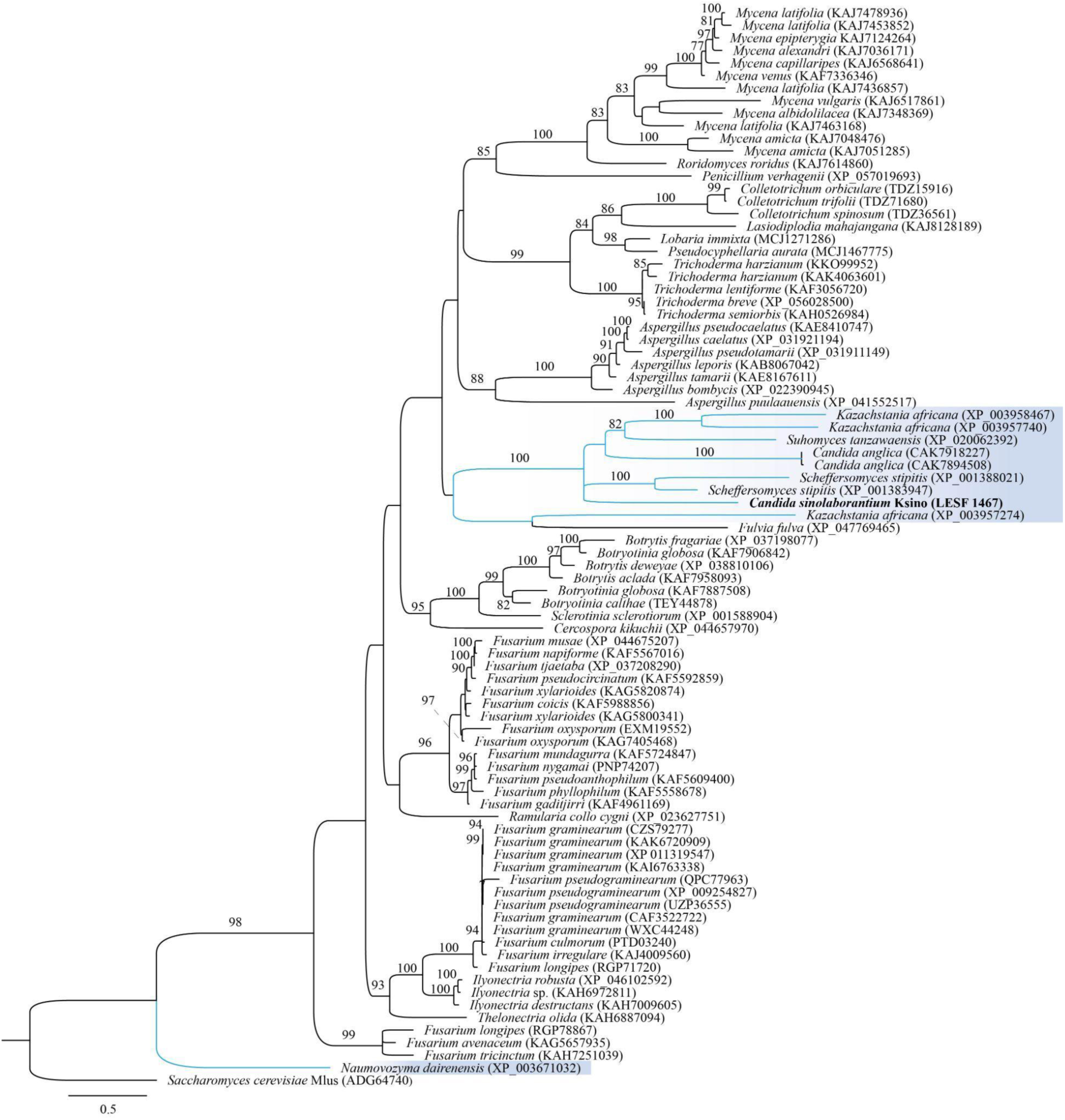
Phylogeny of Ksino homologs identified in diverse fungi. The Ksino-encoding yeast *C. sinolaborantium* is highlighted in bold, and Saccharomycotina in blue. The tree was inferred with maximum likelihood criteria, and numbers on branches are ultrafast bootstrap support values (only values higher than 70 are shown). The dsRNA-encoded killer toxin Klus is represented as an outgroup.

**FIG S6.**
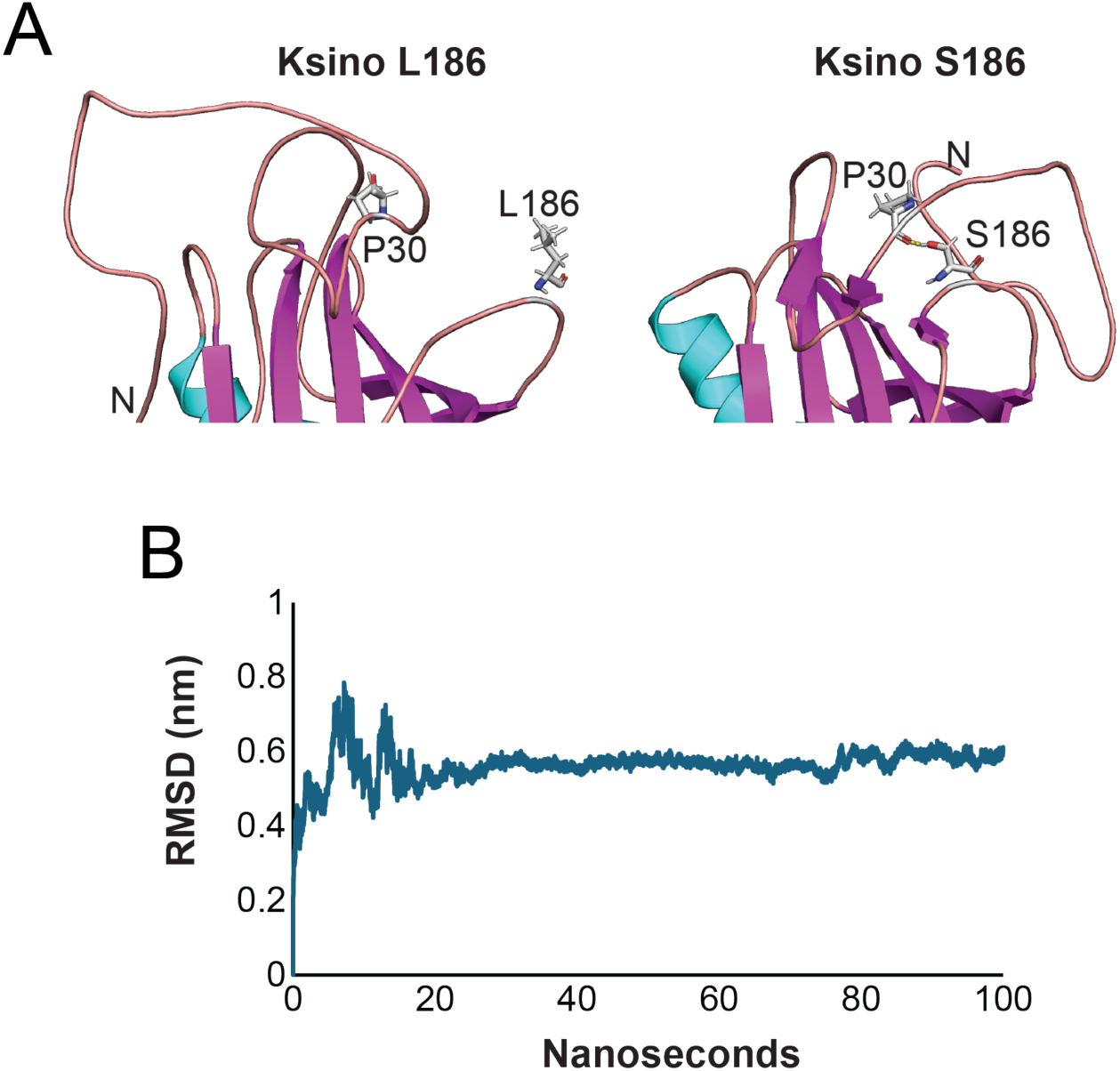
Computational predictions of protein stability with leucine and serine at position 186 in Ksino. **A.** AlphaFold2 models of Ksino wild type S186 and mutant L186 after 100 ns of molecular dynamics simulation. Residues 186 and P30 are shown as sticks. P30 forms a hydrogen bond (1.6 Å) with the hydroxyl of S186 but is too distant to create this bond in L186. FoldX 5.0 analysis revealed a ΔΔG_Folding_ of 1.0 kcal/mol for L186S. **B.** 100 ns MD simulation of Ksino L186 AlphaFold2 model. Unlike wild type and Klus, the RMSD stabilizes around 0.55 nm.

**FIG S7.**
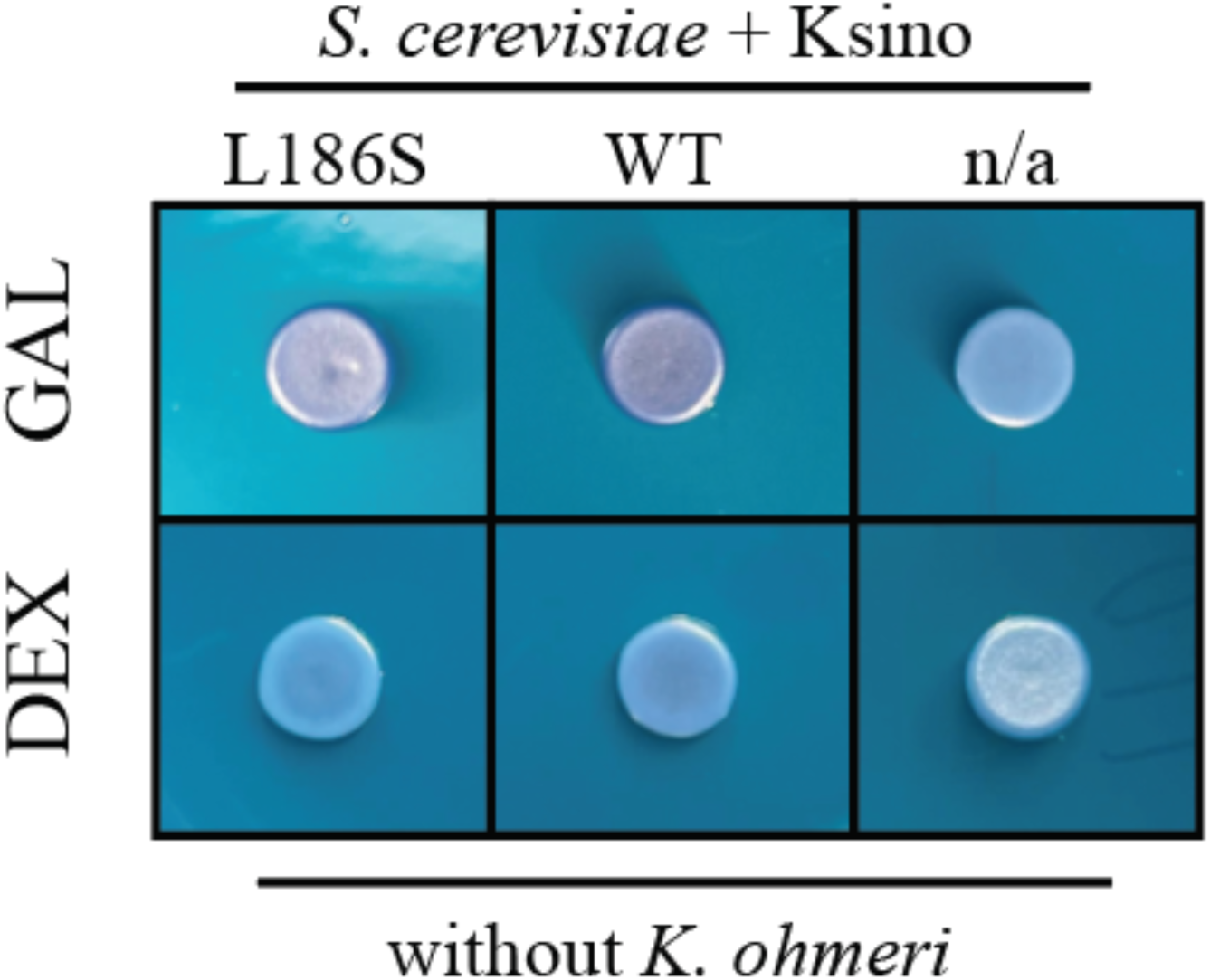
Recombinant expression of Ksino does not cause a white coloration of *S. cerevisiae*. Recombinant expression of Ksino by *S. cerevisiae* using galactose induction without a lawn of *K. ohmeri*. Dextrose was used as a non-induced control. n/a indicates an empty vector negative expression control.

## Supplementary Tables

**Table S1.** Strains and species of yeasts used in the current study.

**Table S2.** The effect of isolation source on killer yeasts activity.

**Table S3.** The effect of pH on the growth inhibition of *Kodamaea ohmeri* LESF 1474 and *C. castelli* by *Candida sinolaborantium* LESF 1467

**Table S4.** The effect of temperature on the growth inhibition of *Kodamaea ohmeri* LESF 1474 and *C. castelli* by *Candida sinolaborantium* LESF 1467.

**Table S5.** The ratio between elongated and yeast morphology of *Kodamaea ohmeri* LESF 1474 cells challenged with killer toxins.

**Table S6.** Cell survival of *Kodamaea ohmeri* LESF 1474 cells against killer toxins.

**Table S7.** Statistics of the assembly, prediction, and annotation of the *Candida sinolaborantium* LESF1467 genome.

**Table S8.** Killer toxins were used as queries for the BLASTp search of the *Candida sinolaborantium* LESF1467 genome.

**Table S9.** Blastp hits of killer toxins towards *Candida sinolaborantium* LESF1467 genome.

**Table S10.** SWISS-MODEL structure assessment scores of Ksino top unrelaxed model and Klus top unrelaxed model compared to models after 100 ns of molecular dynamics simulation. Global pLDDT score is also included in the last row.

**Table S11.** Statistics of Blastp hits of Ksino homologs.

**Table S12.** Nucleotide sequences used in phylogenetic analysis of yeast species delimitation.

**Table S13.** The effect of pH on the growth inhibition of *Kodamaea ohmeri* LESF 1474 and *Candida castyelli* by Ksino expressed by *S. cerevisiae*.

**Table S14.** The effect of temperature on the growth inhibition of *Kodamaea ohmeri* LESF 1474 and *Candida castyelli* by Ksino expressed by *S. cerevisiae*.

**Table S15.** Yeasts used in killer assays as susceptible or killer positive standards.

## Supplementary files

**File S1** - All killer assay data.

**File S2 -** PDB coordinates for the molecular models of Ksino and Klus.

## 4. Materials and Methods

### 4.1. Yeast source and phylogenetic analysis

To investigate the prevalence of killer yeast in the fungus garden of attine ants, a representative collection of 180 yeasts previously characterized by Bizarria Jr. et al. (2023) was examined (Table S1). Briefly, yeasts were obtained by fungus garden suspension in saline solution (0.85% NaCl) supplemented with 0.05% Tween 80. Suspensions were ten-fold diluted and surface spread on nutrient media (Yeast-malt agar and Sabouraud dextrose agar, supplemented with 150 µg mL^−1^ chloramphenicol, and pH adjusted to 4.5; details in Bizarria Jr. et al. 2023). Yeast characterization involved microsatellite amplification analysis (MSP-PCR; Meyer et al. 1993) and sequencing of the D1/D2 region of the large subunit (LSU) ribosomal RNA gene (Kurtzman and Robnett, 1998). Our selection of the 180 *Saccharomycotina* yeasts spanned from fungus gardens of the leaf-cutting ant *Acromyrmex coronatus* (n = 91), the non-leaf-cutting ant *Mycetomoellerius tucumanus* (n = 13), the lower attines *Mycetophylax* aff. *auritus* (n = 42), and *Mycocepurus goeldii* (n = 25), which cultivate *Agaricaceae* fungi; and *Apterostigma goniodes* (n = 9), which cultivates *Pterulaceae* fungi.

To assess the phylogenetic relationship between the yeast strains, we performed a phylogenetic analysis with representative species and their closest relatives (the accession number of sequences used in phylogenetic analysis is listed in Table S12). Sequences were aligned with MAFFT v.7. (Katoh et al. 2019), and nucleotide substitution models were generated using the Bayesian information criterion (BIC) by the standard model generation in IQ-TREE2. SYM+FQ+I+G4 was selected as the nucleotide substitution model. Maximum likelihood (ML) phylogenetic trees were reconstructed with ultrafast bootstrap in IQTREE2 (Nguyen et al. 2015), with 10,000 replicates for ultrafast bootstrap (tree completed after 262 iterations, LogL: -14973.244). The final trees were edited in FigTree v.1.4.3 (Rambaut, 2016).

### 4.2. Killer yeast assays

Killer assays were performed to confirm killer toxin production by yeasts, as described in Crabtree et al. (2019). The 180 yeast strains were seeded onto YPD 4.6 dextrose “killer assay” agar plates (yeast extract 10 g L^−1^, peptone 20 g L^−1^, dextrose 20 g L^−1^, sodium citrate 29.9 g L^−1^, supplemented with 0.003% w/v methylene blue, and pH adjusted to 4.6 with HCl), seeded 10^5^ cells mL^-1^ with one of the 49 selected yeast susceptible strains (i.e. from fungus gardens and sources not related to attine ant environment, Table S13). Nine *Saccharomyces* spp. strains that encode known toxins were used as positive controls for killer toxin productio, including K1, K1L, K21, K2, K28, K45, K62, K74, and Klus (Table S13). Yeasts were also evaluated using 20 pathogenic yeasts as susceptible strains (Table S13). Toxin production was recorded after five days of incubation at room temperature, either by an inhibition zone or a methylene blue stain of susceptible strains. Killer activity severity was scored following Fredericks et al. (2021) by zone of inhibition with or without methylene blue halo or by strong or weak methylene blue halo without inhibition. We applied heatmaps to visualize the species interactions, employing the function heatmap.2 from the gplots package, with the default configuration for dendrogram display (Warnes et al., 2024). To evaluate the effect of the yeast sources (i.e., same or different fungicultures or sites) on killer toxin activity (i.e., killed or non-killed). We selected 9,000 interactions (toward 50 strains that showed susceptibility to killer toxins) and analyzed them with Pearson’s Chi-squared test. All yeasts used in killer assays are listed in Table S15.

### 4.3. Extraction of dsRNA encoding elements

To determine if the yeast strains host dsRNA elements, we extracted the dsRNA elements from all yeast strains following Okada et al. (2015), with modifications as stipulated in Crabtree et al. (2019). The K1 killer yeast *S. cerevisiae* BJH001 was used as a positive control for all assays. Specifically, yeast cultures were grown in yeast-peptone-dextrose broth (YPD: yeast extract 10 g L^−1^, peptone 20 g L^−1^, dextrose 20 g L^−1^), centrifuged at 3000 × g for 5 min, and the broth was removed by aspiration. Cells were resuspended in water, centrifuged at 8,000 × g, and aspirated. A total of 450 µL of 2× sodium chloride-Tris-EDTA buffer (STE: 200 mM NaCl; 20 mM Tris-HCl, pH 8.0; 30 mM EDTA) and 0.5 µL β-mercaptoethanol were added, and cells were disrupted in a cell disruptor (Digital Disruptor Genie, Scientific Industries) for 3 min at 3000 rpm. Then, 50 µL of 10% (w/v) sodium dodecyl sulfate (SDS) solution and 350 µL phenol–chloroform–isoamyl alcohol (25:24:1) were added to crude extracts, which were homogenized and centrifuged at 10,000 × g for 5 min. Then, 500 µL of supernatant was mixed with absolute ethanol (5:1), and the final volume of 600 µL was transferred to cellulose columns (i.e., 0.05 g cellulose D in a 0.6 mL tube bottom punctured and placed in collecting tubes).

Column tubes were centrifuged at 10,000 × g for 30 seconds, and the flow-through was discarded. Then, 350 µL of wash buffer (1× STE containing 16% v/v EtOH) was added to the column, and the tubes were centrifuged at 10,000 × g for 30 seconds, followed by the flow-through discard. Columns were transferred to a collecting tube, adding 350 µL of 1× STE, and tubes were centrifuged at 10,000 × g for 30 seconds. The eluted fraction was recovered, and 1 mL of absolute ethanol with 40 µL of 3 M aqueous sodium acetate (pH 5.2) was added. Tubes were inverted ten times and centrifuged at 20,000 × g for 15 min. The ethanol mix was aspirated without disturbing the precipitated pellet, which was then air-dried and dissolved in 15 µL of molecular-grade water. The presence of dsRNAs was examined (loading 5 µL of each sample) by 0.8% TAE agarose gel electrophoresis, stained with ethidium bromide, at 130 v for 30 min in 1% TAE buffer. As a molecular weight reference, a 1 kb DNA ladder was used. The *Saccharomyces cerevisiae* strain YJM1307, known to host dsRNA totivirus and satellites, was used as positive control.

### 4.4. Extraction of dsDNA linear plasmid

To determine if dsDNA linear plasmids encode the killer toxins, we extracted the dsDNA elements from positive killer yeast, *Kluyveromyces lactis* AWJ137 with the linear DNA plasmids pGKL2 (13.5 kb) and pGKL1 (8.9 kb) and *Pichia acaciae* NRRL Y-18665 with the linear plasmids pPac1-1 (12.6 kb) and pPac1-2 (6.8 kb) (Worsham and Bolen, 1990; Gietz et al., 1995). Cultures were grown in YPD overnight, 200 µL were centrifuged at 3000 × g for 5 min, and the broth was removed by aspiration. Yeast cells were resuspended in 10 µL Zymolase solution (5% of 50 mM Tris-HCl, pH 8; 10% of 5 mM EDTA, and 85% water) and 7.2 µL of 10 mg mL^-1^ of Zymolase (United States Biological Life Sciences, Swampscott, MA), and the suspension was incubated at 30°C for 1 h. After incubation, 1 µL of 10% SDS and 3 µL of proteinase K (New England Biolabs, Ipswich, MA) were added to the suspension, followed by an incubation of 1 h at 65°C. After incubation, 5 µL of loading dye was added, and the suspension was centrifuged at 3000 × g for 5 min. The presence of dsDNA plasmids was examined (loading 10 µL of each sample) by 0.6% TAE agarose gel electrophoresis, stained with ethidium bromide, at 130 v for 1 h in 1% TAE buffer.

### 4.5. Toxin precipitation and killer toxin activity at different temperatures and pH

To investigate the stability of the toxins produced by *C. sinolaborantium* LESF 1467, the isolate extracts on YPD medium (pH 4.6) were filtered, mixed with 1:1 absolute ethanol, centrifuged at 20,000 × g for 20 min, followed by supernatant aspiration and suspension in 10 μL of YPD (pH 4.6). To confirm the killer activity of the precipitates, 4 µL of the suspended proteins were used in killer assays against *Kodamaea ohmeri* LESF 1474. To evaluate the thermostability, the precipitated toxins were screened in killer assays incubating at room temperature or heat-inactivated at 60°C and 98°C for 2 min. For quantitative analysis of killer toxin expression, 2 µl of 10^8^ cells mL^-1^ were plated onto susceptible lawns.The killer activity was also evaluated for activity at different pHs, adjusting the pH to 3.5, 4, 4.5, 5, 5.5, and 6 with HCl 12N, and at various temperatures, incubating at 17, 20, 25, 30, and 35°C. The area of activity on killer assays (i.e., zone of inhibition and cell differentiation of the lawn strain) was measured in cm² after four days with ImageJ 1.53t (Schneider et al., 2012). Since the area data do not satisfy the assumptions of parametric tests, we analyzed variance with Kruskal-Wallis followed by the Wilcoxon rank sum test with Bonferroni correction as a post-hoc for pairwise multiple comparisons. Analyses were performed using the software RStudio v. 2023.12.0.369 (POSIT TEAM, 2023), and R v. 4.3.2 (R CORE TEAM, 2023). Figures were prepared using ggplot2 (Wickham, 2016).

### 4.6. Production of elongated cells by *Kodamaea ohmeri* against killer toxins

We observed that the yeast *K. ohmeri* LESF 1474 produces elongated cells in the presence of *C. sinolaborantium* toxins and their precipitates. We suspended the cells in the white and raised zones of killer assays and on axenic cultures in 1× PBS. Cell morphology was checked under a microscope (Zeiss Primo Star iLED) by counting budding (2 to 17 × 10^6^ cells mL^-1^) and elongated cells (2 to 15 × 10^5^ cells mL^-1^) with a Neubauer chamber. Cell survival was evaluated by adjusting the suspensions to 10^3^ cells mL^-1^ and seeding 100 µL in YPD. Colony-forming units (CFU) were counted after three days of incubation at room temperature. A similar elongated cell phenotype was observed in the presence of *S. cerevisiae* DSM-70459, which secretes the known killer toxin Klus (encoded by dsRNA elements). Microscopic and survival evaluations were also conducted in the presence of *S. cerevisiae* DSM-70459, as described above. To confirm that the elongated cell phenotype is related to the presence of the dsRNA satellites, dsRNA satellites were cured from *S. cerevisiae* by growing on a YPD medium supplemented with 5.0 μM cycloheximide. Loss of killer toxin production and the loss of dsRNAs were assayed as described above. The ratio between normal and elongated cells was compared between *K. ohmeri* in the white halo around *C. sinolaborantium* and *S. cerevisiae* DSM-70459 with axenic cultures surrounding the halo, with ten independent replicates. The data was checked for the assumption of parametric tests (performing Shapiro-Wilk and Bartlett tests, respectively, for normality and homogeneity of variances). The ratio data in both treatments, towards *C. sinolaborantium* and *S. cerevisiae* DSM-70459, were found not to satisfy the assumptions of equal variance (Bartlett test, df = 1, p-value < 0.001), so we conducted the Welch Two Sample t-test to compare means between treatments. The same procedure was performed for cell survival data towards *C. sinolaborantium*, which also did not satisfy the assumptions of equal variance (Bartlett’s K-squared = 4.16, df = 1, p-value = 0.04). Cell survival data towards *S. cerevisiae* DSM-70459 satisfied the assumptions for parametric tests (Bartlett’s K-squared = 0.52, df = 1, p-value = 0.47; Shapiro-Wilk p-value > 0.05), so we conducted the Two Sample t-test to compare means between treatments.

### 4.7. Genomic DNA extraction, sequencing, and annotation

To extract genomic DNA from *C. sinolaborantium*, the yeast was grown in YPD broth; cells were recovered by centrifugation (3000 × g for 5 min), washed, and suspended in 500 µL TE buffer (10 mM Tris-Cl; 0.1 mM EDTA, pH 8). Recovered cells were transferred to a microtube, centrifugation as previously described, and the supernatant was removed. The pellet was disrupted by vortexing. Cells were suspended in a lysis buffer (2% Triton X-100, 1% SDS, 100 mM NaCl, 10 mM Tris-HCl, pH 8, 1 mM EDTA pH 8) with approximately 300 μL of glass beads. A volume of 200 µL of phenol-chloroform-isoamyl alcohol solution was added, and the sample was vortex for 3 min at 3000 rpm (Disruptor Genie), followed by the addition of 200 µL of TE buffer and centrifugation for 5 min at 21000 × g. The upper layer was transferred to a new microtube and mixed by inversion with 1 mL of absolute ethanol. The tube was centrifuged for 3 min at 21000 × g, the supernatant removed, and the pellet suspended in 400 µL of TE buffer. Then, 30 µL of DNAse-free RNAse A was added, and the solution was mixed and incubated for 5 min at 37°C. The DNA was precipitated by adding 10 µL of 4 M ammonium acetate and 1 mL of absolute ethanol, mixing by inversion, and centrifuging for 3 min at 21000 × g. The supernatant was discarded and the pellet was dried for approximately 20 min at room temperature. Genomic DNA was suspended in 100 µL of TE buffer and stored at -20°C.

The genome was sequenced by Illumina NextSeq 550 using the paired-end method. Libraries were created with the Illumina DNA Prep kit for WGS (https://www.illumina.com/products/by-type/sequencing-kits/library-prep-kits/nextera-dna-flex.html). The initial quality analysis and cleaning steps were performed with fastp v. 0.20.1 (CHEN et al., 2018). The genome was assembled with SPAdes v. 3.15.5 (GUREVICH et al., 2013) with multiple k-mers (21, 31, 41, 51, 61, 71, and 81). The blobtools v. 3.5.2 software (LAETSCH; BLAXTER, 2017) was used to check contaminants. Briefly, Blastn 2.13.0+ (ALTSCHUL et al., 1990) was used to search for structures across the entire NCBI nt database; minimap2 v. 2.24-r1122 (LI, 2018) and samtools v.1.13 (LI et al., 2009) were used to obtain sequencing coverage. The blobtools visualization was used to ascertain the results. The rDNA and ITS regions were annotated with ITSx v. 1.3 (BENGTSSON-PALME et al., 2013), which was compared to an amplicon previously submitted to NCBI for this isolate (accession: ON493969) as part of the contamination verification steps. Final assembly statistics were obtained with QUAST v. 5.2.0 (BANKEVICH et al., 2012). Completeness was checked using BUSCO v. 5.4.4 (SEPPEY et al., 2019) with the saccharomycetes_odb10 dataset (2,137 BUSCOs). The genetic prediction was performed using Braker v. 2.1.6 (BRŮNA et al., 2021) [which includes Augustus v. 3.4.0 (STANKE; MORGENSTERN, 2005), ProtHint v. 2.6.0 and GeneMark-ES v. 4.71 (LOMSADZE et al. 2005)] with OrthoDB v. 10 methods (KRIVENTSEVA et al., 2019). InterProScan v. 5.60 (Jones et al., 2014) was used for functional annotation of predicted proteins.

### 4.8. Genome mining for toxin-encoding regions

To identify toxin-encoding chromosomal regions, a search using proteins predicted from whole genome sequencing was performed with BLASTP 2.9.0, using a protein sequence database of known killer toxins. Results with query/hit alignments greater than 100 were selected. An open reading frame similar to the Klus toxin (28% identity, with 151 aa alignment length and an e-value of 1.56E^-05^) was found and selected for further analysis. Polymerase chain reactions (PCR) were performed to amplify the region of interest using primers designed with Primer3 software. 4.1.0 (https://primer3.ut.ee/). The primers were designed to target Ksino untranslated regions (961F: 5’-TTAACGACTTTCGTCTTCGCTATCC-3’ and 961R: 5’-ATTGAGATCAGGTGGCCTGTGTAGC-3’) followed by nested-PCR to amplify the open reading frame using the previous PCR product as the template (961NF: 5’-CGATCACCTAGCCCAAAATGC-3’ and 961NR: 5’-AAAGTGTTGGCCAAGGACACG-3’). Both of the amplification reactions used Phusion high fidelity master mix (New England Biolabs, Ipswich, MA) with the amplification conditions of 98°C for 3 min, 30 cycles at 98°C for 30 s, 63°C for 30 s, 72°C for 30 s, and a final extension step at 72°C for 5 min. Amplicons were sequenced to confirm the identity of the region of interest.

### 4.9. Prediction of tertiary structure and comparative analysis of proteins

After confirming the genomic coding region of Ksino was the correct size using PCR, the sequence was used to predict protein structure via the AlphaFold2 software (Jumper et al., 2021) with default parameters. The B factor columns of each unrelaxed structure were used to create per residue pLDDT graphs. Secondary structure was assigned using PyMol to create a linear representation of the protein and aligned to per residue graphs. The top-scoring relaxed model from AlphaFold2 was used for molecular dynamics simulation using the GROMACS package (Spoel et al., 2005). Briefly, the system was set up using the AMBER99SB-ILDN force field with TIP3P water, a dodecahedral box with 1.0 nm padding, and 0.15 M NaCl. The system was then subjected to energy minimization by steepest descent, followed by NVT and NPT equilibration with a pressure of 1.0 atm and temperature of 300 K. Pressure and temperature were maintained constant using arrinello-Rahman with isotropic coupling and V-rescale with Protein Non-protein as the coupling groups, respectively. This setup was chosen to match standard cellular conditions closely: 26.85°C, 0.15 M NaCl, pH 7, and 1 atm. Simulations were run for 100 ns with a 2.0 fs timestep, and snapshots were saved every 2 fs. The RMSD was calculated as a function of time using the resulting trajectories. In addition, the final snapshot of each trajectory was saved as a PDB structure file and compared to relaxed and unrelaxed AlphaFold2 models using the SWISS structure assessment tool (Studer et al., 2019).

FoldX was used to predict the ΔΔG_Folding_ of L186S (Delgado et al., 2019). Amber relaxed Ksino L186 was used as the starting structure, and FoldX PDBrepair was carried out six times to optimize the structure. Then, L186 in the optimized structure was mutated to every other residue using PositionScan. ΔΔG_Folding_ was taken from the output file.

### 4.10. Bioinformatic discovery of Ksino homologs in fungi

A search for similar sequences in NCBI (https://www.ncbi.nlm.nih.gov/) was also carried out with Blastp, with the results filtered by alignment length (above 109), identity (above 30%), and e-value (less than 0.03). The protein sequences were aligned with MUSCLE (https://www.ebi.ac.uk/jdispatcher/msa/muscle), with the alignment consisting of 86 sequences and 426 positions. Amino acid substitution models were generated according to the Bayesian information criterion (BIC), with WAG+R4 being the chosen model. Maximum likelihood (ML) phylogenetic trees were reconstructed in IQTREE2 (Nguyen et al., 2015), with 10,000 ultrafast bootstrap replicates (tree completed after 163 iterations, LogL: -18525.485). The final trees were edited in FigTree v.1.4.3 (RAMBAUT, 2016).

### 4.11. Cloning and transformation in *Escherichia coli*

To study the coding region, a “TA” terminal was added to the amplicon in a reaction that included 1 μL Taq polymerase, 19 μL of PCR product, 1 μL of 10 μM dNTPs, 5 μL of 10×buffer, and 24 µL of sterile ultrapure water. The reaction was incubated at 72°C for 20 min. For cloning, the pCR™8/GW/TOPO™ TA Cloning kit (Invitrogen, Waltham, MA) was used, adding 1 μL of amplicons with TA terminals, 0.25 μL of saline solution from the kit, and 0.25 μL of the pCR™8/GW/TOPO™ vector. The reaction was incubated at room temperature for 2 h, followed by adding 25 μL of competent *E. coli* cells (One-Shot TOP10 Competent Cells, Invitrogen, Waltham, MA). The reaction was kept on ice for 30 min, followed by heat shock at 42°C for 30 s and then ice bath for 2 min. Subsequently, 250 μL of Super Optimal broth (SOC: 20 g L^-1^ Tryptone, 5 g L^-1^ Yeast extract, 0.58 g L^-1^ NaCl, 0.19 g L^-1^ KCl, 2, 03 g L^-1^ MgCl_2_, 1.20 g L^-1^ MgSO_4_, and 3.6 g L^-1^ dextrose) preheated to 37°C was added, and incubated at 37°C for 1 h. Then, 250 μL were plated on Luria bertani medium (LB: 10 g L^-1^ of tryptone, 5 g L^-1^ of yeast extract, 10 g L^-1^ of NaCl, and 15 g L^-1^ of agar) supplemented with 100 µg mL-1 of spectinomycin. Plasmids were extracted using the High-Speed Mini Plasmid kit (IBI Scientific, Dubuque, IA), following manufacturer recommendations.

Due to the transcriptional diversity of CUG-Leu codons in some yeasts (Krassowski et al., 2018), the vectors containing the region of interest were subjected to targeted mutations (SDM: Site-Directed Mutagenesis) in a CUG codon starting at position 556 of the DNA sequence (position 186 in the protein sequence). Thus, the mutants consist of the alternatives identified for CUG in other yeasts, being called L186A and L186S, with L being the original amino acid, 186 the position, and “S” the mutation for Ser. To obtain the mutants, we design primers targeting position 186 as follows: 961GCG: 5’-CAGCTGTGGCGCGAACCATAGCGGG-3’, 961TCG: 5’-CAGCTGTGGCTCGAACCATAGCGGG-3’ (forward), and 961SNM: 5’-AGTCCCGCAGGCCAATCC-3’ (reverse). Amplification reactions included 12.5 μL of Phusion High Fidelity Master Mix, 1 μL 10 μM forward primer, 1 μL 10 μM reverse primer, 9.5 μL sterile ultrapure water, 1 μL 1:9 diluted DNA (i.e., pCR™8 plasmid with the region of interest). Amplification conditions were as follows: 98°C for 30 min, 25 cycles at 98°C for 5 s, 67°C for 10 s, 72°C for 50 s, and a final extension step at 72°C for 5 min. To remove the template DNA from the reaction, a Dpnl treatment was performed by mixing 22 μL of PCR product, 2 μL Dpnl, 3 μL CutSmart buffer, and 3 μL sterile ultrapure water. The reaction was incubated at 37°C for 1 day. For the addition of 5’-P and 3’-OH, we performed a PNK treatment: 30 μL of the treated product, 5 μL PNK buffer, 1 μL PNK, 5 μL 10 mM ATP, 1 μL 0.1 M DTT, 8 μL of sterile ultrapure water. The reactions were incubated for 30 min at 37°C and inactivated at 65°C for 10 min. The product was purified using the Monarch DNA & PCR Cleanup kit (New England Biolabs, Ipswich, MA), following the manufacturer’s protocols. To re-ligate the plasmids, the following reaction was performed: 2 μL of purified product, 1 μL T4 DNA ligase, 2 μL 10× binding buffer, 15 μL sterile ultrapure water. The reaction was incubated for 2 h at room temperature, and then 25 μL of competent *E. coli* cells (One-Shot TOP10 Competent Cells, Invitrogen, Waltham, MA) were added. Transformed cultures were selected in LB supplemented with 100 µg mL^-1^ spectinomycin, plasmids extracted with the High-Speed Mini Plasmid kit (IBI Scientific, Dubuque, IA), and mutations checked by sequencing (ELIM Biopharmaceuticals, Hayward, CA).

The vectors were subjected to a Gateway™ recombination from the vectors in pCR8 to pAG306-GAL-ccdB. For recombination, the reaction was conducted with 0.5 μL of the source vector, 0.5 μL of the destination vector, 1 μL of sterile ultrapure water, and 0.5 μL of LR Clonase II (New England Biolabs, Ipswich, MA). The reaction was incubated for 4 h at room temperature, followed by adding 0.25 μL of Proteinase K (New England Biolabs, Ipswich, MA) and incubating at 37°C for 10 min. Finally, the *E. coli* transformation was performed as previously described, and cultures were selected in LB supplemented with 100 μg mL^-1^ of chloramphenicol and ampicillin.

### 4.12. Transformation and heterologous expression of killer toxins by*Saccharomyces cerevisiae*

For yeast transformation, the integrative vector containing a killer toxin gene was linearized with the NcoI enzyme by cutting at the *URA3* locus, followed by inactivation of the enzyme at 65°C for 30 min. The product was purified using the Monarch DNA & PCR Cleanup kit (New England Biolabs, Ipswich, MA), following the manufacturer’s recommendations. *S. cerevisiae* CRY1 cells were grown in YPD until an optical density of 1.3 was obtained, then the cells were washed with sterile water and suspended in 10 mL of TE buffer supplemented with 100 mM LiAc, and incubated at 30°C for 30 min under 130 rpm agitation. Cells were recovered by centrifugation at 1500 × g for 5 min at 4°C and suspended in 5 mL of ice-cold 1M sorbitol, followed by centrifugation and suspended in 550 μL of ice-cold 1M sorbitol. For transformation, 80 μL of the suspended cells were added to 5 to 10 μg of the linearized plasmids and transferred to electroporation cuvettes. The cuvettes were incubated on ice for 10 min and pulsed with GenePulser (200 Ω, 1.5 kV, and 25 μF). To recover the transformed cultures, the cells were grown at 30°C for 2 days in CM medium without uracil (complete media: 2.5 g L^-1^ of the drop-out mix without uracil, 1.7 g L^-1^ of yeast nitrogen base YNB, 5 g L^-1^ ammonium sulfate, 20 g L^-1^ dextrose). For killer toxin expression was induced by overnight growth in YPGal (yeast extract 10 g L^-1^, peptone 20 g L^-1^, galactose 20 g L^-1^) followed by plating on either YPD or YPGal with citrate-phosphate 29.9 g L^-1^, supplemented with 0.003% w/v methylene blue and pH adjusted to 4.6 with HCl and 15 g L^-1^ agar seeded with 10^5^ cells mL^-1^ of a susceptible lawn strain. *C. sinolaborantium* and *S. cerevisiae* CRY1 were used as positive and negative controls for killer activity, respectively.

### 4.13. Detection of killer toxin expression by RT-PCR

An overnight culture of Ksino was grown in YPD and harvested at OD600 0.9-1.2. Speroplasting was performed by incubating the harvested cells at 30°C for 30 min in buffer Y1 (1M sorbitol, 0.1 M EDTA, pH 7.4, B-ME 0.1%, and 0.1% v/v zymolase 20T (10 mg/mL)). Spheroplasts were used as input for a Qiagen RNeasy procedure. The RNA quality was checked using a spectrophotometer, and 10 μg of total RNA was treated with 1 μL of DNase I (ThermoFisher Scientific) in the presence of 10 μL of DNase reaction buffer at 37°C for 10 min, and then inactivated by heat at 75°C for 10 min. Transcripts encoding Ksino or actin were detected by RT-PCR (Superscript IV) following the manufacturer’s instructions using the primers 5’-GGCCTTGAGATACCCCATCG-3’ and 5’-CCACGTGAGTAACACCGTCA-3’ (for *ACT1*) and 5’-GATCTACGACAGCAGCCTAGA-3’ and 5’-GTGGTGTACATCGTTGACTCC-3’ (for Ksino). Briefly, the RNA template is denatured with a reverse primer mix at 65°C for 5 min and incubated on ice for at least 1 min. The reverse transcription reaction mix was added to the denatured RNA and incubated for 10 minutes at 55°C and then inactivated at 80°C for 10 minutes. A 20 L PCR was performed with the cycling instructions as follows (98°C for 30 s, then 30 cycles of 98°C for 10 s, 60°C for 30 s, 72°C for 10 s followed by one cycle of 72°C for 5 min). PCR products were analyzed by agarose gel electrophoresis.

