## Supplementary material for "The Prevalence of Killer Yeasts in the Gardens of Fungus-Growing Ants and the Discovery of a Novel Killer Toxin named Ksino": Supp Tables

**Supplementary Material 1**

**Table S1.** Strains and species of yeasts used in the current study.

| Strain | Species | Family | Order | Ant source | Nest ID | City (Brazil) |
|---|---|---|---|---|---|---|
| LESF 1481 | *Blastobotrys proliferans* | Trichomonascaceae | Dipodascales | *Acromyrmex coronatus* | RB190919-01 | Rio Claro-SP |
| LESF 1442 | *Candida albicans* | Debaryomycetaceae | Serinales | *Acromyrmex coronatus* | RB200104-03 | Mogi-Guaçu-SP |
| LESF 1417 | *Candida blattae* | Debaryomycetaceae | Serinales | *Mycetophylax* aff. *auritus* | AR210126-05 | Itatiaia-RJ |
| LESF 1440 | *Candida blattae* | Debaryomycetaceae | Serinales | *Mycetophylax* aff. *auritus* | AR210126-05 | Itatiaia-RJ |
| LESF 1401 | *Candida blattae* | Debaryomycetaceae | Serinales | *Mycetophylax* aff. *auritus* | AR210126-05 | Itatiaia-RJ |
| LESF 1494 | *Candida blattae* | Debaryomycetaceae | Serinales | *Apterostigma goniodes* | AR210127-02 | Itatiaia-RJ |
| LESF 1450 | *Candida margitis* | Debaryomycetaceae | Serinales | *Acromyrmex coronatus* | RB190909-01 | Rio Claro-SP |
| LESF 1451 | *Candida margitis* | Debaryomycetaceae | Serinales | *Acromyrmex coronatus* | RB190909-01 | Rio Claro-SP |
| LESF 1453 | *Candida margitis* | Debaryomycetaceae | Serinales | *Acromyrmex coronatus* | RB190909-01 | Rio Claro-SP |
| LESF 1377 | *Candida margitis* | Debaryomycetaceae | Serinales | *Acromyrmex coronatus* | RB190909-01 | Rio Claro-SP |
| LESF 1378 | *Candida margitis* | Debaryomycetaceae | Serinales | *Acromyrmex coronatus* | RB190909-01 | Rio Claro-SP |
| LESF 1381 | *Candida margitis* | Debaryomycetaceae | Serinales | *Acromyrmex coronatus* | RB190909-01 | Rio Claro-SP |
| LESF 1364 | *Candida orthopsilosis* | Debaryomycetaceae | Serinales | *Acromyrmex coronatus* | RB191007-01 | Rio Claro-SP |
| LESF 1394 | *Candida orthopsilosis* | Debaryomycetaceae | Serinales | *Acromyrmex coronatus* | RB191007-01 | Rio Claro-SP |
| LESF 1402 | *Candida orthopsilosis* | Debaryomycetaceae | Serinales | *Acromyrmex coronatus* | RB191007-01 | Rio Claro-SP |
| LESF 1403 | *Candida orthopsilosis* | Debaryomycetaceae | Serinales | *Acromyrmex coronatus* | RB191007-01 | Rio Claro-SP |

| | | | | | | |
|---|---|---|---|---|---|---|
| LESF 1477 | *Candida orthopsilosis* | Debaryomycetaceae | Serinales | *Acromyrmex coronatus* | RB191007-01 | Rio Claro-SP |
| LESF 1363 | *Candida orthopsilosis* | Debaryomycetaceae | Serinales | *Acromyrmex coronatus* | RB191007-01 | Rio Claro-SP |
| LESF 1407 | *Candida orthopsilosis* | Debaryomycetaceae | Serinales | *Acromyrmex coronatus* | RB191007-01 | Rio Claro-SP |
| LESF 1373 | *Candida orthopsilosis* | Debaryomycetaceae | Serinales | *Acromyrmex coronatus* | RB191007-01 | Rio Claro-SP |
| LESF 1392 | *Candida orthopsilosis* | Debaryomycetaceae | Serinales | *Acromyrmex coronatus* | RB191007-01 | Rio Claro-SP |
| LESF 1374 | *Candida orthopsilosis* | Debaryomycetaceae | Serinales | *Acromyrmex coronatus* | RB191007-01 | Rio Claro-SP |
| LESF 1380 | *Candida orthopsilosis* | Debaryomycetaceae | Serinales | *Acromyrmex coronatus* | RB191007-01 | Rio Claro-SP |
| LESF 1390 | *Candida orthopsilosis* | Debaryomycetaceae | Serinales | *Acromyrmex coronatus* | RB191007-01 | Rio Claro-SP |
| LESF 1371 | *Candida orthopsilosis* | Debaryomycetaceae | Serinales | *Acromyrmex coronatus* | RB191007-01 | Rio Claro-SP |
| LESF 1370 | *Candida orthopsilosis* | Debaryomycetaceae | Serinales | *Acromyrmex coronatus* | RB191007-01 | Rio Claro-SP |
| LESF 1347 | *Candida parapsilosis* | Debaryomycetaceae | Serinales | *Mycocepurus goeldii* | RB210430-03 | Anhembi-SP |
| LESF 1333 | *Candida parapsilosis* | Debaryomycetaceae | Serinales | *Mycocepurus goeldii* | RB210501-02 | Anhembi-SP |
| LESF 1384 | *Candida railenensis* | Debaryomycetaceae | Serinales | *Acromyrmex coronatus* | RB200104-02 | Mogi-Guaçu-SP |
| LESF 1386 | *Candida railenensis* | Debaryomycetaceae | Serinales | *Acromyrmex coronatus* | RB200104-02 | Mogi-Guaçu-SP |
| LESF 1387 | *Candida railenensis* | Debaryomycetaceae | Serinales | *Acromyrmex coronatus* | RB200104-02 | Mogi-Guaçu-SP |
| LESF 1456 | *Candida railenensis* | Debaryomycetaceae | Serinales | *Acromyrmex coronatus* | RB200104-02 | Mogi-Guaçu-SP |
| LESF 1388 | *Candida railenensis* | Debaryomycetaceae | Serinales | *Acromyrmex coronatus* | RB200104-02 | Mogi-Guaçu-SP |
| LESF 1455 | *Candida railenensis* | Debaryomycetaceae | Serinales | *Acromyrmex coronatus* | RB200104-02 | Mogi-Guaçu-SP |
| LESF 1393 | *Candida railenensis* | Debaryomycetaceae | Serinales | *Acromyrmex coronatus* | RB200104-02 | Mogi-Guaçu-SP |
| LESF 1341 | *Candida railenensis* | Debaryomycetaceae | Serinales | *Acromyrmex coronatus* | RB200104-02 | Mogi-Guaçu-SP |
| LESF 1437 | *Candida railenensis* | Debaryomycetaceae | Serinales | *Acromyrmex coronatus* | RB200104-03 | Mogi-Guaçu-SP |
| LESF 1449 | *Candida railenensis* | Debaryomycetaceae | Serinales | *Acromyrmex coronatus* | RB200104-03 | Mogi-Guaçu-SP |
| LESF 1349 | *Candida railenensis* | Debaryomycetaceae | Serinales | *Mycocepurus goeldii* | RB210501-06 | Anhembi-SP |
| LESF 1354 | *Candida railenensis* | Debaryomycetaceae | Serinales | *Mycocepurus goeldii* | RB210501-06 | Anhembi-SP |
| LESF 1467 | *Candida sinolaborantium* | Debaryomycetaceae | Serinales | *Mycetophylax* aff. *auritus* | AR210126-03 | Itatiaia-RJ |
| LESF 1395 | *Candida* sp. (closest to *C. intermedia*) | Debaryomycetaceae | Serinales | *Acromyrmex coronatus* | RB191007-01 | Rio Claro-SP |

| | | | | | | |
|---|---|---|---|---|---|---|
| LESF 1359 | *Candida* sp. (closest to *C. intermedia*) | Debaryomycetaceae | Serinales | *Acromyrmex coronatus* | RB191007-01 | Rio Claro-SP |
| LESF 1433 | *Candida* sp. (closest to *C. vrieseae*) | Debaryomycetaceae | Serinales | *Mycetophylax* aff. *auritus* | AR210126-04 | Itatiaia-RJ |
| LESF 1461 | *Candida* sp. (closest to *C. temnochilae*) | Debaryomycetaceae | Serinales | *Mycetophylax* aff. *auritus* | AR210126-05 | Itatiaia-RJ |
| LESF 1372 | *Candida* sp. (closest to *C. temnochilae*) | Debaryomycetaceae | Serinales | *Mycetophylax* aff. *auritus* | PK210128-01 | Itatiaia-RJ |
| LESF 1413 | *Candida* sp. (closest to *C. auris*) | Debaryomycetaceae | Serinales | *Mycetophylax* aff. *auritus* | PK210128-01 | Itatiaia-RJ |
| LESF 1493 | *Candida* sp. (closest to *C. vrieseae*) | Debaryomycetaceae | Serinales | *Apterostigma goniodes* | AR210127-02 | Itatiaia-RJ |
| LESF 1438 | *Cyberlindnera* sp. | Phaffomycetaceae | Phaffomycetales | *Mycetophylax* aff. *auritus* | AR210126-02 | Itatiaia-RJ |
| LESF 1405 | *Debaryomyces hansenii* | Debaryomycetaceae | Serinales | *Acromyrmex coronatus* | RB191007-01 | Rio Claro-SP |
| LESF 1446 | *Diutina catenulata* | Debaryomycetaceae | Serinales | *Acromyrmex coronatus* | RB200104-03 | Mogi-Guaçu-SP |
| LESF 1439 | *Diutina catenulata* | Debaryomycetaceae | Serinales | *Acromyrmex coronatus* | RB200104-03 | Mogi-Guaçu-SP |
| LESF 1396 | *Diutina catenulata* | Debaryomycetaceae | Serinales | *Acromyrmex coronatus* | RB200104-03 | Mogi-Guaçu-SP |
| LESF 1416 | *Diutina catenulata* | Debaryomycetaceae | Serinales | *Acromyrmex coronatus* | RB200104-03 | Mogi-Guaçu-SP |
| LESF 1501 | *Diutina catenulata* | Debaryomycetaceae | Serinales | *Acromyrmex coronatus* | RB200104-03 | Mogi-Guaçu-SP |
| LESF 1419 | *Diutina catenulata* | Debaryomycetaceae | Serinales | *Acromyrmex coronatus* | RB200104-03 | Mogi-Guaçu-SP |
| LESF 1345 | *Grigorovia* sp. | Saccharomycetaceae | Saccharomycetales | *Mycocepurus goeldii* | RB210501-06 | Anhembi-SP |
| LESF 1470 | *Hanseniaspora opuntiae* | Saccharomycodaceae | Saccharomycodales | *Acromyrmex coronatus* | RB200104-03 | Mogi-Guaçu-SP |
| LESF 1488 | *Hanseniaspora opuntiae* | Saccharomycodaceae | Saccharomycodales | *Acromyrmex coronatus* | RB200104-03 | Mogi-Guaçu-SP |
| LESF 1484 | *Hanseniaspora opuntiae* | Saccharomycodaceae | Saccharomycodales | *Acromyrmex coronatus* | RB200104-03 | Mogi-Guaçu-SP |
| LESF 1297 | *Hanseniaspora thailandica* | Saccharomycodaceae | Saccharomycodales | *Acromyrmex coronatus* | RB200104-02 | Mogi-Guaçu-SP |
| LESF 1298 | *Hanseniaspora thailandica* | Saccharomycodaceae | Saccharomycodales | *Acromyrmex coronatus* | RB200104-02 | Mogi-Guaçu-SP |
| LESF 1483 | *Hanseniaspora thailandica* | Saccharomycodaceae | Saccharomycodales | *Acromyrmex coronatus* | RB200104-03 | Mogi-Guaçu-SP |
| LESF 1435 | *Hanseniaspora thailandica* | Saccharomycodaceae | Saccharomycodales | *Acromyrmex coronatus* | RB200104-03 | Mogi-Guaçu-SP |
| LESF 1486 | *Hanseniaspora thailandica* | Saccharomycodaceae | Saccharomycodales | *Acromyrmex coronatus* | RB200104-04 | Mogi-Guaçu-SP |
| LESF 1503 | *Hyphopichia burtonii* | Debaryomycetaceae | Serinales | *Mycetomoellerius tucumanus* | AR201207-02 | Rio Claro-SP |
| LESF 1505 | *Hyphopichia burtonii* | Debaryomycetaceae | Serinales | *Mycetomoellerius tucumanus* | AR201207-02 | Rio Claro-SP |
| LESF 1504 | *Hyphopichia burtonii* | Debaryomycetaceae | Serinales | *Mycetomoellerius tucumanus* | AR201207-02 | Rio Claro-SP |

| | | | | | | |
|---|---|---|---|---|---|---|
| LESF 1499 | *Hyphopichia burtonii* | Debaryomycetaceae | Serinales | *Mycetomoellerius tucumanus* | RB201207-01 | Rio Claro-SP |
| LESF 1344 | *Hyphopichia* sp. | Debaryomycetaceae | Serinales | *Mycetophylax* aff. *auritus* | AR210126-04 | Itatiaia-RJ |
| LESF 1468 | *Kazachstania spencerorum* | Saccharomycetaceae | Saccharomycetales | *Mycetophylax* aff. *auritus* | RB210126-02 | Itatiaia-RJ |
| LESF 1474 | *Kodamaea ohmeri* | Debaryomycetaceae | Serinales | *Mycetophylax* aff. *auritus* | PK210128-01 | Itatiaia-RJ |
| LESF 1421 | *Kurtzmaniella quercitrusa* | Debaryomycetaceae | Serinales | *Mycetophylax* aff. *auritus* | AR210126-01 | Itatiaia-RJ |
| LESF 1489 | *Kurtzmaniella quercitrusa* | Debaryomycetaceae | Serinales | *Apterostigma goniodes* | RB210126-03 | Itatiaia-RJ |
| LESF 1398 | *Kurtzmaniella quercitrusa* | Debaryomycetaceae | Serinales | *Apterostigma goniodes* | RB210126-03 | Itatiaia-RJ |
| LESF 1400 | *Kurtzmaniella quercitrusa* | Debaryomycetaceae | Serinales | *Apterostigma goniodes* | AR210127-02 | Itatiaia-RJ |
| LESF 1498 | *Kurtzmaniella quercitrusa* | Debaryomycetaceae | Serinales | *Apterostigma goniodes* | AR210127-02 | Itatiaia-RJ |
| LESF 1399 | *Kurtzmaniella quercitrusa* | Debaryomycetaceae | Serinales | *Apterostigma goniodes* | AR210127-02 | Itatiaia-RJ |
| LESF 1473 | *Kurtzmaniella* sp. | Debaryomycetaceae | Serinales | *Mycetophylax* aff. *auritus* | PK210126-03 | Itatiaia-RJ |
| LESF 1491 | *Kurtzmaniella* sp. | Debaryomycetaceae | Serinales | *Mycetophylax* aff. *auritus* | PK210128-01 | Itatiaia-RJ |
| LESF 1500 | *Limtongozyma cylindracea* | Debaryomycetaceae | Serinales | *Mycetophylax* aff. *auritus* | AR210126-02 | Itatiaia-RJ |
| LESF 1404 | *Limtongozyma* sp. | Debaryomycetaceae | Serinales | *Mycetophylax* aff. *auritus* | PK210126-03 | Itatiaia-RJ |
| LESF 1492 | *Limtongozyma* sp. | Debaryomycetaceae | Serinales | *Mycetophylax* aff. *auritus* | PK210128-01 | Itatiaia-RJ |
| LESF 1368 | *Lodderomyces elongisporus* | Debaryomycetaceae | Serinales | *Acromyrmex coronatus* | RB191007-01 | Rio Claro-SP |
| LESF 1360 | *Lodderomyces elongisporus* | Debaryomycetaceae | Serinales | *Acromyrmex coronatus* | RB191007-01 | Rio Claro-SP |
| LESF 1362 | *Lodderomyces elongisporus* | Debaryomycetaceae | Serinales | *Acromyrmex coronatus* | RB191007-01 | Rio Claro-SP |
| LESF 1365 | *Lodderomyces elongisporus* | Debaryomycetaceae | Serinales | *Acromyrmex coronatus* | RB191007-01 | Rio Claro-SP |
| LESF 1422 | *Metschnikowia koreensis* | Metschnikowiaceae | Serinales | *Mycetomoellerius tucumanus* | AR201207-03 | Rio Claro-SP |
| LESF 1343 | *Metschnikowia* sp. | Metschnikowiaceae | Serinales | *Mycetomoellerius tucumanus* | LV201207-01 | Rio Claro-SP |
| LESF 1415 | *Metschnikowia* sp. | Metschnikowiaceae | Serinales | *Mycetomoellerius tucumanus* | LV201207-01 | Rio Claro-SP |
| LESF 1383 | *Metschnikowia* sp. | Metschnikowiaceae | Serinales | *Mycetophylax* aff. *auritus* | PK210128-01 | Itatiaia-RJ |
| LESF 1355 | *Metschnikowia* sp. | Metschnikowiaceae | Serinales | *Mycocepurus goeldii* | RB210501-05 | Anhembi-SP |
| LESF 1332 | *Metschnikowia* sp. | Metschnikowiaceae | Serinales | *Mycocepurus goeldii* | RB210501-05 | Anhembi-SP |
| LESF 1375 | *Meyerozyma caribbica* | Debaryomycetaceae | Serinales | *Acromyrmex coronatus* | RB191007-01 | Rio Claro-SP |

| | | | | | | |
|---|---|---|---|---|---|---|
| LESF 1382 | *Meyerozyma caribbica* | Debaryomycetaceae | Serinales | *Acromyrmex coronatus* | RB191007-01 | Rio Claro-SP |
| LESF 1443 | *Meyerozyma caribbica* | Debaryomycetaceae | Serinales | *Acromyrmex coronatus* | RB200104-03 | Mogi-Guaçu-SP |
| LESF 1452 | *Meyerozyma caribbica* | Debaryomycetaceae | Serinales | *Acromyrmex coronatus* | RB200104-03 | Mogi-Guaçu-SP |
| LESF 1389 | *Meyerozyma carpophila* | Debaryomycetaceae | Serinales | *Acromyrmex coronatus* | RB191007-01 | Rio Claro-SP |
| LESF 1406 | *Meyerozyma guilliermondii* | Debaryomycetaceae | Serinales | *Acromyrmex coronatus* | RB191007-01 | Rio Claro-SP |
| LESF 1444 | *Meyerozyma guilliermondii* | Debaryomycetaceae | Serinales | *Acromyrmex coronatus* | RB191007-01 | Rio Claro-SP |
| LESF 1379 | *Meyerozyma guilliermondii* | Debaryomycetaceae | Serinales | *Acromyrmex coronatus* | RB191007-01 | Rio Claro-SP |
| LESF 1411 | *Pichia kudriavzevii* | Pichiaceae | Pichiales | *Acromyrmex coronatus* | RB200104-03 | Mogi-Guaçu-SP |
| LESF 1391 | *Priceomyces* sp. | Debaryomycetaceae | Serinales | *Acromyrmex coronatus* | RB200507-03 | Rio Claro-SP |
| LESF 1445 | Saccharomycetales sp. | Debaryomycetaceae | Serinales | *Acromyrmex coronatus* | RB190919-01 | Rio Claro-SP |
| LESF 1457 | *Scheffersomyces* sp. | Debaryomycetaceae | Serinales | *Mycetophylax* aff. *auritus* | AR210126-05 | Itatiaia-RJ |
| LESF 1448 | *Scheffersomyces* sp. | Debaryomycetaceae | Serinales | *Mycetophylax* aff. *auritus* | RB210126-02 | Itatiaia-RJ |
| LESF 1408 | *Scheffersomyces* sp. | Debaryomycetaceae | Serinales | *Mycetophylax* aff. *auritus* | PK210126-03 | Itatiaia-RJ |
| LESF 1464 | *Scheffersomyces* sp. | Debaryomycetaceae | Serinales | *Mycetophylax* aff. *auritus* | PK210126-03 | Itatiaia-RJ |
| LESF 1497 | *Schwanniomyces vanrijiae* | Debaryomycetaceae | Serinales | *Mycetophylax* aff. *auritus* | AR210126-04 | Itatiaia-RJ |
| LESF 1475 | *Schwanniomyces vanrijiae* | Debaryomycetaceae | Serinales | *Mycetophylax* aff. *auritus* | AR210126-05 | Itatiaia-RJ |
| LESF 1335 | *Schwanniomyces vanrijiae* | Debaryomycetaceae | Serinales | *Mycetophylax* aff. *auritus* | RB210126-02 | Itatiaia-RJ |
| LESF 1482 | *Schwanniomyces vanrijiae* | Debaryomycetaceae | Serinales | *Mycetophylax* aff. *auritus* | RB210126-02 | Itatiaia-RJ |
| LESF 1480 | *Schwanniomyces vanrijiae* | Debaryomycetaceae | Serinales | *Mycetophylax* aff. *auritus* | RB210126-02 | Itatiaia-RJ |
| LESF 1478 | *Schwanniomyces vanrijiae* | Debaryomycetaceae | Serinales | *Mycetophylax* aff. *auritus* | RB210126-02 | Itatiaia-RJ |
| LESF 1479 | *Schwanniomyces vanrijiae* | Debaryomycetaceae | Serinales | *Mycetophylax* aff. *auritus* | RB210126-02 | Itatiaia-RJ |
| LESF 1521 | *Schwanniomyces vanrijiae* | Debaryomycetaceae | Serinales | *Apterostigma goniodes* | AR210127-02 | Itatiaia-RJ |
| LESF 1472 | *Starmerella bombicola* | Trichomonascaceae | Dipodascales | *Acromyrmex coronatus* | RB190909-01 | Rio Claro-SP |
| LESF 1376 | *Starmerella bombicola* | Trichomonascaceae | Dipodascales | *Acromyrmex coronatus* | RB190909-01 | Rio Claro-SP |
| LESF 1476 | *Starmerella bombicola* | Trichomonascaceae | Dipodascales | *Acromyrmex coronatus* | RB190909-01 | Rio Claro-SP |
| LESF 1471 | *Starmerella bombicola* | Trichomonascaceae | Dipodascales | *Acromyrmex coronatus* | RB190909-01 | Rio Claro-SP |

| | | | | | | |
|---|---|---|---|---|---|---|
| LESF 1487 | *Starmerella etchellsii* | Trichomonascaceae | Dipodascales | *Acromyrmex coronatus* | RB200507-01 | Rio Claro-SP |
| LESF 1299 | *Starmerella etchellsii* | Trichomonascaceae | Dipodascales | *Acromyrmex coronatus* | RB200507-01 | Rio Claro-SP |
| LESF 1300 | *Starmerella etchellsii* | Trichomonascaceae | Dipodascales | *Acromyrmex coronatus* | RB200507-01 | Rio Claro-SP |
| LESF 1485 | *Starmerella etchellsii* | Trichomonascaceae | Dipodascales | *Acromyrmex coronatus* | RB200507-01 | Rio Claro-SP |
| LESF 1301 | *Starmerella etchellsii* | Trichomonascaceae | Dipodascales | *Acromyrmex coronatus* | RB200507-01 | Rio Claro-SP |
| LESF 1304 | *Starmerella etchellsii* | Trichomonascaceae | Dipodascales | *Acromyrmex coronatus* | RB200507-01 | Rio Claro-SP |
| LESF 1302 | *Starmerella etchellsii* | Trichomonascaceae | Dipodascales | *Acromyrmex coronatus* | RB200507-01 | Rio Claro-SP |
| LESF 1303 | *Starmerella etchellsii* | Trichomonascaceae | Dipodascales | *Acromyrmex coronatus* | RB200507-01 | Rio Claro-SP |
| LESF 1508 | *Starmerella etchellsii* | Trichomonascaceae | Dipodascales | *Acromyrmex coronatus* | RB200507-01 | Rio Claro-SP |
| LESF 1496 | *Starmerella meliponinorum* | Trichomonascaceae | Dipodascales | *Acromyrmex coronatus* | RB200507-01 | Rio Claro-SP |
| LESF 1420 | *Starmerella meliponinorum* | Trichomonascaceae | Dipodascales | *Acromyrmex coronatus* | RB200507-01 | Rio Claro-SP |
| LESF 1418 | *Starmerella meliponinorum* | Trichomonascaceae | Dipodascales | *Acromyrmex coronatus* | RB200507-01 | Rio Claro-SP |
| LESF 1293 | *Starmerella* sp. | Trichomonascaceae | Dipodascales | *Acromyrmex coronatus* | RB190909-01 | Rio Claro-SP |
| LESF 1454 | *Starmerella* sp. | Trichomonascaceae | Dipodascales | *Acromyrmex coronatus* | RB190919-01 | Rio Claro-SP |
| LESF 1294 | *Starmerella* sp. | Trichomonascaceae | Dipodascales | *Acromyrmex coronatus* | RB190919-01 | Rio Claro-SP |
| LESF 1295 | *Starmerella* sp. | Trichomonascaceae | Dipodascales | *Acromyrmex coronatus* | RB190919-01 | Rio Claro-SP |
| LESF 1431 | *Starmerella* sp. | Trichomonascaceae | Dipodascales | *Acromyrmex coronatus* | RB200507-01 | Rio Claro-SP |
| LESF 1465 | *Sugiyamaella bonitensis* | Trichomonascaceae | Dipodascales | *Mycetophylax* aff. *auritus* | AR210126-01 | Itatiaia-RJ |
| LESF 1366 | *Sugiyamaella boreocaroliniensis* | Trichomonascaceae | Dipodascales | *Mycocepurus goeldii* | RB210430-06 | Anhembi-SP |
| LESF 1463 | *Sugiyamaella carassensis* | Trichomonascaceae | Dipodascales | *Mycetophylax* aff. *auritus* | AR210126-01 | Itatiaia-RJ |
| LESF 1338 | *Sugiyamaella carassensis* | Trichomonascaceae | Dipodascales | *Mycetophylax* aff. *auritus* | AR210126-01 | Itatiaia-RJ |
| LESF 1356 | *Sugiyamaella carassensis* | Trichomonascaceae | Dipodascales | *Mycocepurus goeldii* | RB210501-06 | Anhembi-SP |
| LESF 1459 | *Sugiyamaella ligni* | Trichomonascaceae | Dipodascales | *Mycetophylax* aff. *auritus* | AR210126-01 | Itatiaia-RJ |
| LESF 1414 | *Sugiyamaella ligni* | Trichomonascaceae | Dipodascales | *Mycetophylax* aff. *auritus* | AR210126-01 | Itatiaia-RJ |
| LESF 1462 | *Sugiyamaella ligni* | Trichomonascaceae | Dipodascales | *Mycetophylax* aff. *auritus* | AR210126-01 | Itatiaia-RJ |
| LESF 1351 | *Sugiyamaella smithiae* | Trichomonascaceae | Dipodascales | *Mycocepurus goeldii* | RB210430-06 | Anhembi-SP |

| | | | | | | |
|---|---|---|---|---|---|---|
| LESF 1352 | *Sugiyamaella smithiae* | Trichomonascaceae | Dipodascales | *Mycocepurus goeldii* | RB210430-06 | Anhembi-SP |
| LESF 1353 | *Sugiyamaella smithiae* | Trichomonascaceae | Dipodascales | *Mycocepurus goeldii* | RB210430-06 | Anhembi-SP |
| LESF 1334 | *Sugiyamaella smithiae* | Trichomonascaceae | Dipodascales | *Mycocepurus goeldii* | RB210430-06 | Anhembi-SP |
| LESF 1357 | *Sugiyamaella smithiae* | Trichomonascaceae | Dipodascales | *Mycocepurus goeldii* | RB210430-06 | Anhembi-SP |
| LESF 1348 | *Sugiyamaella smithiae* | Trichomonascaceae | Dipodascales | *Mycocepurus goeldii* | RB210430-06 | Anhembi-SP |
| LESF 1369 | *Sugiyamaella smithiae* | Trichomonascaceae | Dipodascales | *Mycocepurus goeldii* | RB210430-06 | Anhembi-SP |
| LESF 1367 | *Sugiyamaella smithiae* | Trichomonascaceae | Dipodascales | *Mycocepurus goeldii* | RB210430-06 | Anhembi-SP |
| LESF 1361 | *Sugiyamaella smithiae* | Trichomonascaceae | Dipodascales | *Mycocepurus goeldii* | RB210430-06 | Anhembi-SP |
| LESF 1350 | *Sugiyamaella smithiae* | Trichomonascaceae | Dipodascales | *Mycocepurus goeldii* | RB210430-06 | Anhembi-SP |
| LESF 1337 | *Sugiyamaella smithiae* | Trichomonascaceae | Dipodascales | *Mycocepurus goeldii* | RB210430-06 | Anhembi-SP |
| LESF 1336 | *Sugiyamaella smithiae* | Trichomonascaceae | Dipodascales | *Mycocepurus goeldii* | RB210430-06 | Anhembi-SP |
| LESF 1340 | *Sugiyamaella smithiae* | Trichomonascaceae | Dipodascales | *Mycocepurus goeldii* | RB210430-06 | Anhembi-SP |
| LESF 1342 | *Sugiyamaella smithiae* | Trichomonascaceae | Dipodascales | *Mycocepurus goeldii* | RB210430-06 | Anhembi-SP |
| LESF 1346 | *Sugiyamaella smithiae* | Trichomonascaceae | Dipodascales | *Mycocepurus goeldii* | RB210430-06 | Anhembi-SP |
| LESF 1296 | *Wickerhamiella occidentalis* | Trichomonascaceae | Dipodascales | *Acromyrmex coronatus* | RB200104-01 | Mogi-Guaçu-SP |
| LESF 1502 | *Wickerhamiella occidentalis* | Trichomonascaceae | Dipodascales | *Mycetophylax* aff. *auritus* | PK210126-03 | Itatiaia-RJ |
| LESF 1506 | *Wickerhamiella* sp. | Trichomonascaceae | Dipodascales | *Acromyrmex coronatus* | RB200507-02 | Rio Claro-SP |
| LESF 1441 | *Wickerhamiella* sp. | Trichomonascaceae | Dipodascales | *Acromyrmex coronatus* | RB200507-02 | Rio Claro-SP |
| LESF 1495 | *Wickerhamiella* sp. | Trichomonascaceae | Dipodascales | *Mycocepurus goeldii* | RB210430-06 | Anhembi-SP |
| LESF 1397 | *Wickerhamomyces ciferrii* | Wickerhamomycetaceae | Phaffomycetales | *Mycetomoellerius tucumanus* | AR201207-02 | Rio Claro-SP |
| LESF 1436 | *Wickerhamomyces ciferrii* | Wickerhamomycetaceae | Phaffomycetales | *Mycetomoellerius tucumanus* | AR201207-02 | Rio Claro-SP |
| LESF 1409 | *Wickerhamomyces ciferrii* | Wickerhamomycetaceae | Phaffomycetales | *Mycetomoellerius tucumanus* | AR201207-02 | Rio Claro-SP |
| LESF 1410 | *Wickerhamomyces ciferrii* | Wickerhamomycetaceae | Phaffomycetales | *Mycetomoellerius tucumanus* | AR201207-02 | Rio Claro-SP |
| LESF 1339 | *Wickerhamomyces ciferrii* | Wickerhamomycetaceae | Phaffomycetales | *Mycetomoellerius tucumanus* | AR201207-02 | Rio Claro-SP |
| LESF 1466 | *Wickerhamomyces ciferrii* | Wickerhamomycetaceae | Phaffomycetales | *Mycetomoellerius tucumanus* | RB201207-01 | Rio Claro-SP |
| LESF 1460 | *Wickerhamomyces* sp. | Wickerhamomycetaceae | Phaffomycetales | *Mycetophylax* aff. *auritus* | AR210126-02 | Itatiaia-RJ |

| | | | | | | |
|---|---|---|---|---|---|---|
| LESF 1458 | *Wickerhamomyces* sp. | Wickerhamomycetaceae | Phaffomycetales | *Mycetophylax* aff. *auritus* | AR210126-02 | Itatiaia-RJ |
| LESF 1412 | *Yamadazyma paraaseri* | Debaryomycetaceae | Serinales | *Mycetophylax* aff. *auritus* | AR210126-02 | Itatiaia-RJ |
| LESF 1434 | *Yamadazyma* sp. | Debaryomycetaceae | Serinales | *Mycetophylax* aff. *auritus* | AR210126-03 | Itatiaia-RJ |
| LESF 1490 | *Yamadazyma* sp. | Debaryomycetaceae | Serinales | *Mycetophylax* aff. *auritus* | AR210126-03 | Itatiaia-RJ |
| LESF 1507 | *Yamadazyma* sp. | Debaryomycetaceae | Serinales | *Apterostigma goniodes* | AR210127-02 | Itatiaia-RJ |
| LESF 1385 | *Yarrowia lipolytica* | Incertis sedis | Dipodascales | *Acromyrmex coronatus* | RB191007-01 | Rio Claro-SP |
| LESF 1358 | *Yarrowia lipolytica* | Incertis sedis | Dipodascales | *Acromyrmex coronatus* | RB191007-01 | Rio Claro-SP |
| LESF 1447 | *Yarrowia lipolytica* | Incertis sedis | Dipodascales | *Acromyrmex coronatus* | RB191007-01 | Rio Claro-SP |

**Table S2.** The effect of isolation source on killer yeasts activity.

| **Statistics of interactions with 69 lawn strains** | |
|---|---|
| Total of interactions | 12,420 |
| **Statistics of interactions with 50 susceptible strains** | |
| Total of interactions | 9,000 |
| Killed phenotypes (same source) | 6 |
| Killed phenotypes (different source) | 301 |
| Not killed phenotypes (same source) | 641 |
| Not killed phenotypes (different source) | 8,052 |

Pearson's Chi-squared test, $\chi2 = 13.05$, df = 1, p-value = $3.03E^{-04}$.

**Table S3.** The effect of pH on the growth inhibition of *Kodamaea ohmeri* LESF 1474 and *Candida castyelli* by *Candida sinolaborantium* LESF 1467.

| | *Kodamaea ohmeri* | *Candida castelli* |
|---|---|---|
| Treatment | Area in cm² (Mean ± SE) | |
| pH 3.5 | 1.03 ± 0.04 | 6.65 ± 0.33 |
| pH 4.0 | 1.18 ± 0.04 | 8.61 ± 0.65 |
| pH 4.5 | 2.35 ± 0.06 | 8.15 ± 0.92 |
| pH 5.0 | 3.25 ± 0.09 | 5.74 ± 1.58 |
| pH 5.5 | 0.81 ± 0.03 | 2.08 ± 0.38 |
| pH 6.0 | 0 | 1.90 ± 0.16 |
| pH 6.5 | 0 | 1.57 ± 0.11 |

**Table S4.** The effect of temperature on the growth inhibition of *Kodamaea ohmeri* LESF 1474 and *Candida castyelli* by *Candida sinolaborantium* LESF 1467.

| | *Kodamaea ohmeri* | *Candida castelli* |
|---|---|---|
| Treatment | Area in cm² (Mean ± SE) | |
| 17°C | 2.03 ± 0.04 | 8.15 ± 0.73 |
| 20°C | 2.42 ± 0.08 | 8.42 ± 0.47 |
| 25°C | 2.15 ± 0.05 | 8.02 ± 0.49 |
| 30°C | 1.01 ± 0.03 | 5.39 ± 0.10 |

**Table S5.** The ratio between elongated and normal *Kodamaea ohmeri* LESF 1474 cells challenged with killer toxins.

| Treatment | Value (Mean ± SE) |
|---|---|
| Control | 0.03 ± 0.00 |
| LESF1467 | 0.24 ± 0.02 |
| Klus | 0.49 ± 0.07 |

LESF1467 × Control: Welch Two Sample t-test, $t = -12.18$, $df = 9.26$, p-value = $5.194E^{-07}$

Klus × Control: Welch Two Sample t-test, t = -6.78, df = 9.02, p-value = $8.034E^{-05}$

**Table S6.** Cell survival of *Kodamaea ohmeri* LESF 1474 cells against killer toxins.

| Treatment | CFUs (Mean ± SE) |
|---|---|
| Control | $1.47E^{+02} \pm 1.31E^{+01}$ |
| LESF1467 | $7.00E^{+01} \pm 6.33E^{+00}$ |
| Klus | $1.30E^{+02} \pm 1.02E^{+01}$ |

LESF1467 × Control: Welch Two Sample t-test, t = 5.30, df = 12.99, p-value = $1.45E^{-04}$
Klus × Control: Two Sample t-test, t = 1.02, df = 18, p-value = 0.3194

**Table S7.** Statistics of the assembly, prediction, and annotation of the *Candida sinolaborantium* LESF1467 genome.

| **Statistics of cleaning steps** | | |
|---|---|---|
| Category | Before filtering | After filtering |
| Total of reads | 5,240,040 | 4,203,746 |
| Total of bases | 786,006,000 | 608,646,389 |
| Ratio Q30 | 69.15% | 75.61% |
| GC contente | 46.15% | 44.46% |
| **BUSCO and QUAST assembly statistics** | | |
| Category | Value | |
| Total BUSCO groups searched (saccharomycetes_odb10) | 2,137 | |
| Complete BUSCOs | 2,097 (98.2%) | |
| Complete and simple copy BUSCOs | 2,099 (98.1%) | |
| Complete and duplicates BUSCOs | 2 (0.1%) | |
| Fragmented BUSCOs | 1.00% | |
| Absent BUSCOs | 0.80% | |
| Contigs > 200 bp | 270 | |

| | |
|---|---|
| Contigs > 1,000 bp | 163 |
| Total length (>= 0 bp) | 11,214,429 |
| Total length (>= 1,000 bp) | 11,175,526 |
| Largest contig | 543,051 |
| GC (%) | 48.76 |
| N50 | 155,979 |
| N's per 100 kbp | 29.07 |
| **Gene prediction and annotation statistics** | |
| Category | Value |
| Number of protein-coding genes | 5,951 |

**Table S8.** Killer toxins used as queries for the BLASTp search on *Candida sinolaborantium* LESF1467 genome.

| Toxin | Organism | GenBank accession |
|---|---|---|
| HM-1 | *Cyberlindnera mrakii* | P10410 |
| SMK | *Millerozyma farinosa* | P19972 |
| K1 | *Saccharomyces cerevisiae* | AII19505 |
| K2 | *Saccharomyces cerevisiae* | CAA39941 |
| KHR | *Saccharomyces cerevisiae* | P22313 |
| KHS | *Saccharomyces cerevisiae* | P39690 |
| Klus | *Saccharomyces cerevisiae* | ADG64740 |
| K1L | *Saccharomyces paradoxus* | QQX23408 |
| K21 | *Saccharomyces paradoxus* | ATN38270 |
| K28 | *Saccharomyces paradoxus* | ATN38273 |
| K28 | *Saccharomyces paradoxus* | AII19507 |
| K45 | *Saccharomyces paradoxus* | ATN38271 |
| K62 | *Saccharomyces paradoxus* | ATN38496 |

| | | |
|---|---|---|
| K66 | *Saccharomyces paradoxus* | AYN80721 |
| K74 | *Saccharomyces paradoxus* | ATN38272 |
| Kbarr-1 | *Torulaspora delbrueckii* | ALI93476 |
| KP4 | *Ustilago maydis* | Q90121 |
| zygocin | *Zygosaccharomyces bailii* | AAM54023 |

**Table S9.** Blastp hits of killer toxins towards *Candida sinolaborantium* LESF1467 genome.

| Query | % Identity | Alignment length | Mismatches | e-value | Bit score |
|---|---|---|---|---|---|
| K2 (CAA39941) | 25.15 | 167 | 118 | 3.16E-07 | 49.3 |
| K21 (ATN38270) | 27.35 | 117 | 60 | 1.40E+00 | 28.5 |
| K21 (ATN38270) | 24.62 | 130 | 90 | 5.20E+00 | 26.9 |
| K28 (AII19507) | 23.59 | 106 | 69 | 7.60E+00 | 26.2 |
| K28 (ATN38273) | 24.38 | 160 | 99 | 4.30E+00 | 27.3 |
| K45 (ATN38271) | 28.21 | 117 | 57 | 9.30E+00 | 26.2 |
| K66 (AYN80721) | 26.52 | 132 | 85 | 8.70E+00 | 26.2 |
| K74 (ATN38272) | 25.52 | 145 | 76 | 9.10E+00 | 26.2 |
| KHR (P22313) | 23.53 | 187 | 101 | 7.30E+00 | 26.2 |
| KHS (P39690) | 26.98 | 126 | 78 | 3.50E+00 | 28.9 |
| KHS (P39690) | 24.00 | 100 | 75 | 5.90E+00 | 28.1 |
| Klus (ADG64740) | 27.82 | 151 | 94 | 1.56E-05 | 42.4 |

**Table S10.** SWISS-MODEL structure assessment scores of Ksino top unrelaxed model and Klus top unrelaxed model compared to models after 100 ns of molecular dynamics simulation. Global pLDDT score is also included in the last row.

| | Ksino unrelaxed | Ksino relaxed | Ksino 100ns | Klus unrelaxed | Klus relaxed | Klus 100ns |
|---|---|---|---|---|---|---|
| MolProbity score (version 4.4) | 3.44 | 2.39 | 1.54 | 3.51 | 2.15 | 1.86 |

| | | | | | | |
|---|---|---|---|---|---|---|
| Clash score | 32.64 | 5.43 | 1.03 | 37.61 | 2.97 | 1.1 |
| Ramachandran favoured | 77.00% | 87.50% | 91.16% | 72.92% | 82.08% | 89.04% |
| Ramachandran outliers | 15.50% | 5.50% | 0.55% | 17.50% | 7.08% | 1.83% |
| Rotamer outliers | 7.50% | 4.38% | 1.99% | 6.86% | 2.94% | 4.19% |
| Bad bonds | 64/1545 | 0/1546 | 28/1534 | 57/1936 | 0/1937 | 34/1925 |
| Bad angles | 119/2094 | 14/2096 | 77/2063 | 119/2651 | 275273 | 122/2619 |
| Twisted non-proline | 22/196 | 4/196 | 1/196 | 12/235 | 4/235 | 2/235 |
| Global pLDDT | 63645 | - | - | 65086 | - | - |

**Table S11.** Statistics of Blastp hits of Ksino homologs.

| GenBank accession | % Identity | Alignment length | e-value | Species | Family | Class | Phylum |
|---|---|---|---|---|---|---|---|
| XP_022390945 | 33.33 | 144 | 2.34E-08 | *Aspergillus bombycis* | Aspergillaceae | Eurotiomycetes | Ascomycota |
| XP_031921194 | 35.17 | 145 | 2.88E-09 | *Aspergillus caelatus* | Aspergillaceae | Eurotiomycetes | Ascomycota |
| KAB8067042 | 31.97 | 147 | 1.30E-08 | *Aspergillus leporis* | Aspergillaceae | Eurotiomycetes | Ascomycota |
| KAE8410747 | 35.17 | 145 | 1.86E-09 | *Aspergillus pseudocaelatus* | Aspergillaceae | Eurotiomycetes | Ascomycota |
| XP_031911149 | 40.00 | 110 | 1.61E-10 | *Aspergillus pseudotamarii* | Aspergillaceae | Eurotiomycetes | Ascomycota |
| XP_041552517 | 33.03 | 109 | 7.00E-03 | *Aspergillus puulaauensis* | Aspergillaceae | Eurotiomycetes | Ascomycota |
| KAE8167611 | 32.65 | 147 | 1.26E-08 | *Aspergillus tamarii* | Aspergillaceae | Eurotiomycetes | Ascomycota |
| TEY44878 | 36.49 | 148 | 1.88E-10 | *Botryotinia calthae* | Sclerotiniaceae | Leotiomycetes | Ascomycota |
| KAF7887508 | 37.01 | 154 | 8.86E-12 | *Botryotinia globosa* | Sclerotiniaceae | Leotiomycetes | Ascomycota |
| KAF7906842 | 35.14 | 148 | 1.94E-10 | *Botryotinia globosa* | Sclerotiniaceae | Leotiomycetes | Ascomycota |
| KAF7958093 | 33.56 | 149 | 1.61E-08 | *Botrytis aclada* | Sclerotiniaceae | Leotiomycetes | Ascomycota |
| XP_038810106 | 34.87 | 152 | 3.70E-10 | *Botrytis deweyae* | Sclerotiniaceae | Leotiomycetes | Ascomycota |
| XP_037198077 | 32.20 | 118 | 1.00E-03 | *Botrytis fragariae* | Sclerotiniaceae | Leotiomycetes | Ascomycota |
| CAK7894508 | 37.24 | 196 | 2.03E-34 | *Candida anglica* | Debaryomycetaceae | Pichiomycetes | Ascomycota |
| CAK7918227 | 37.24 | 196 | 8.83E-35 | *Candida anglica* | Debaryomycetaceae | Pichiomycetes | Ascomycota |

| | | | | | | | |
|---|---|---|---|---|---|---|---|
| XP_044657970 | 35.95 | 153 | 2.04E-10 | *Cercospora kikuchii* | Mycosphaerellaceae | Dothideomycetes | Ascomycota |
| TDZ15916 | 32.65 | 147 | 4.82E-07 | *Colletotrichum orbiculare* | Mycosphaerellaceae | Dothideomycetes | Ascomycota |
| TDZ36561 | 32.65 | 147 | 2.88E-07 | *Colletotrichum spinosum* | Mycosphaerellaceae | Dothideomycetes | Ascomycota |
| TDZ71680 | 33.33 | 147 | 4.78E-08 | *Colletotrichum trifolii* | Mycosphaerellaceae | Dothideomycetes | Ascomycota |
| XP_047769465 | 31.91 | 141 | 9.70E-09 | *Fulvia fulva* | Mycosphaerellaceae | Dothideomycetes | Ascomycota |
| KAG5657935 | 36.09 | 133 | 2.72E-11 | *Fusarium avenaceum* | Nectriaceae | Sordariomycetes | Ascomycota |
| KAF5988856 | 32.99 | 194 | 1.76E-09 | *Fusarium coicis* | Nectriaceae | Sordariomycetes | Ascomycota |
| PTD03240 | 30.77 | 156 | 9.14E-10 | *Fusarium culmorum* | Nectriaceae | Sordariomycetes | Ascomycota |
| KAF4961169 | 30.38 | 158 | 6.92E-08 | *Fusarium gaditjirri* | Nectriaceae | Sordariomycetes | Ascomycota |
| CZS79277 | 31.41 | 156 | 9.22E-10 | *Fusarium graminearum* | Nectriaceae | Sordariomycetes | Ascomycota |
| KAK6720909 | 31.41 | 156 | 8.76E-10 | *Fusarium graminearum* | Nectriaceae | Sordariomycetes | Ascomycota |
| WXC44248 | 31.41 | 156 | 7.59E-10 | *Fusarium graminearum* | Nectriaceae | Sordariomycetes | Ascomycota |
| XP_011319547 | 31.41 | 156 | 7.20E-10 | *Fusarium graminearum* | Nectriaceae | Sordariomycetes | Ascomycota |
| CAF3522722 | 31.17 | 154 | 1.15E-08 | *Fusarium graminearum* | Nectriaceae | Sordariomycetes | Ascomycota |
| KAI6763338 | 31.17 | 154 | 1.09E-08 | *Fusarium graminearum* | Nectriaceae | Sordariomycetes | Ascomycota |
| KAJ4009560 | 31.41 | 156 | 7.84E-10 | *Fusarium irregulare* | Nectriaceae | Sordariomycetes | Ascomycota |
| RGP71720 | 32.05 | 156 | 1.64E-10 | *Fusarium longipes* | Nectriaceae | Sordariomycetes | Ascomycota |
| RGP78867 | 35.95 | 153 | 1.11E-11 | *Fusarium longipes* | Nectriaceae | Sordariomycetes | Ascomycota |
| KAF5724847 | 30.06 | 163 | 2.08E-06 | *Fusarium mundagurra* | Nectriaceae | Sordariomycetes | Ascomycota |
| XP_044675207 | 35.50 | 169 | 2.36E-10 | *Fusarium musae* | Nectriaceae | Sordariomycetes | Ascomycota |
| KAF5567016 | 33.73 | 169 | 1.03E-10 | *Fusarium napiforme* | Nectriaceae | Sordariomycetes | Ascomycota |
| PNP74207 | 31.01 | 158 | 3.92E-08 | *Fusarium nygamai* | Nectriaceae | Sordariomycetes | Ascomycota |
| KAG7405468 | 31.79 | 195 | 8.58E-09 | *Fusarium oxysporum* | Nectriaceae | Sordariomycetes | Ascomycota |
| EXM19552 | 31.48 | 162 | 6.45E-09 | *Fusarium oxysporum* | Nectriaceae | Sordariomycetes | Ascomycota |
| KAF5558678 | 30.43 | 161 | 1.05E-04 | *Fusarium phyllophilum* | Nectriaceae | Sordariomycetes | Ascomycota |
| KAF5609400 | 31.01 | 158 | 5.77E-08 | *F. pseudoanthophilum* | Nectriaceae | Sordariomycetes | Ascomycota |
| KAF5592859 | 35.50 | 169 | 1.80E-09 | *Fusarium pseudocircinatum* | Nectriaceae | Sordariomycetes | Ascomycota |

| | | | | | | | |
|---|---|---|---|---|---|---|---|
| QPC77963 | 32.05 | 156 | 2.98E-10 | *F. pseudograminearum* | Nectriaceae | Sordariomycetes | Ascomycota |
| UZP36555 | 31.41 | 156 | 9.71E-10 | *F. pseudograminearum* | Nectriaceae | Sordariomycetes | Ascomycota |
| XP_009254827 | 31.41 | 156 | 7.67E-10 | *F. pseudograminearum* | Nectriaceae | Sordariomycetes | Ascomycota |
| XP_037208290 | 33.14 | 169 | 7.70E-10 | *Fusarium tjaetaba* | Nectriaceae | Sordariomycetes | Ascomycota |
| KAH7251039 | 33.33 | 147 | 3.24E-08 | *Fusarium tricinctum* | Nectriaceae | Sordariomycetes | Ascomycota |
| KAG5800341 | 31.79 | 195 | 1.28E-08 | *Fusarium xylarioides* | Nectriaceae | Sordariomycetes | Ascomycota |
| KAG5820874 | 32.99 | 194 | 6.82E-09 | *Fusarium xylarioides* | Nectriaceae | Sordariomycetes | Ascomycota |
| KAH7009605 | 33.78 | 148 | 5.84E-11 | *Ilyonectria destructans* | Nectriaceae | Sordariomycetes | Ascomycota |
| XP_046102592 | 34.46 | 148 | 1.34E-12 | *Ilyonectria robusta* | Nectriaceae | Sordariomycetes | Ascomycota |
| KAH6972811 | 36.04 | 111 | 5.15E-10 | *Ilyonectria* sp. | Nectriaceae | Sordariomycetes | Ascomycota |
| XP_003958467 | 39.90 | 198 | 1.70E-31 | *Kazachstania africana* | Saccharomycetaceae | Saccharomycetes | Ascomycota |
| XP_003957740 | 36.78 | 174 | 4.53E-27 | *Kazachstania africana* | Saccharomycetaceae | Saccharomycetes | Ascomycota |
| XP_003957274 | 33.06 | 124 | 1.12E-04 | *Kazachstania africana* | Saccharomycetaceae | Saccharomycetes | Ascomycota |
| KAJ8128189 | 31.90 | 116 | 6.00E-03 | *Lasiodiplodia mahajangana* | Botryosphaeriaceae | Dothideomycetes | Ascomycota |
| MCJ1271286 | 34.76 | 164 | 2.13E-11 | *Lobaria immixta* | Peltigerineae | Lecanoromycetes | Ascomycota |
| XP_003671032 | 31.97 | 122 | 1.81E-04 | *Naumovozyma dairenensis* | Saccharomycetaceae | Saccharomycetes | Ascomycota |
| XP_057019693 | 30.23 | 129 | 1.74E-04 | *Penicillium verhagenii* | Aspergillaceae | Eurotiomycetes | Ascomycota |
| MCJ1467775 | 31.10 | 164 | 7.55E-08 | *Pseudocyphellaria aurata* | Peltigerineae | Lecanoromycetes | Ascomycota |
| XP_023627751 | 33.94 | 109 | 4.76E-06 | *Ramularia collo_cygni* | Mycosphaerellaceae | Dothideomycetes | Ascomycota |
| XP_001383947 | 42.02 | 188 | 2.48E-35 | *Scheffersomyces stipitis* | Debaryomycetaceae | Pichiomycetes | Ascomycota |
| XP_001388021 | 37.06 | 143 | 7.55E-17 | *Scheffersomyces stipitis* | Debaryomycetaceae | Pichiomycetes | Ascomycota |
| XP_001588904 | 35.86 | 145 | 2.58E-08 | *Sclerotinia sclerotiorum* | Sclerotiniaceae | Leotiomycetes | Ascomycota |
| XP_020062392 | 35.08 | 191 | 4.32E-25 | *Suhomyces tanzawaensis* | Debaryomycetaceae | Pichiomycetes | Ascomycota |
| KAH6887094 | 32.21 | 149 | 2.98E-09 | *Thelonectria olida* | Nectriaceae | Sordariomycetes | Ascomycota |
| XP_056028500 | 33.56 | 146 | 1.12E-08 | *Trichoderma breve* | Hypocreaceae | Sordariomycetes | Ascomycota |
| KAK4063601 | 31.61 | 155 | 2.93E-08 | *Trichoderma harzianum* | Hypocreaceae | Sordariomycetes | Ascomycota |
| KKO99952 | 32.88 | 146 | 9.39E-08 | *Trichoderma harzianum* | Hypocreaceae | Sordariomycetes | Ascomycota |

| | | | | | | | |
|---|---|---|---|---|---|---|---|
| KAF3056720 | 32.88 | 146 | 1.34E-08 | *Trichoderma lentiforme* | Hypocreaceae | Sordariomycetes | Ascomycota |
| KAH0526984 | 33.56 | 146 | 1.18E-08 | *Trichoderma semiorbis* | Hypocreaceae | Sordariomycetes | Ascomycota |
| KAJ7348369 | 31.94 | 144 | 2.32E-07 | *Mycena albidolilacea* | Marasmiineae | Agaricomycetes | Basidiomycota |
| KAJ7036171 | 32.26 | 155 | 3.97E-07 | *Mycena alexandri* | Marasmiineae | Agaricomycetes | Basidiomycota |
| KAJ7048476 | 30.99 | 171 | 1.25E-07 | *Mycena amicta* | Marasmiineae | Agaricomycetes | Basidiomycota |
| KAJ7051285 | 31.25 | 144 | 2.43E-07 | *Mycena amicta* | Marasmiineae | Agaricomycetes | Basidiomycota |
| KAJ6568641 | 32.26 | 124 | 3.00E-03 | *Mycena capillaripes* | Marasmiineae | Agaricomycetes | Basidiomycota |
| KAJ7124264 | 32.64 | 144 | 4.13E-05 | *Mycena epipterygia* | Marasmiineae | Agaricomycetes | Basidiomycota |
| KAJ7463168 | 30.19 | 159 | 1.87E-08 | *Mycena latifolia* | Marasmiineae | Agaricomycetes | Basidiomycota |
| KAJ7436857 | 30.32 | 155 | 8.19E-06 | *Mycena latifolia* | Marasmiineae | Agaricomycetes | Basidiomycota |
| KAJ7478936 | 30.32 | 155 | 3.00E-03 | *Mycena latifolia* | Marasmiineae | Agaricomycetes | Basidiomycota |
| KAJ7453852 | 31.45 | 124 | 3.00E-02 | *Mycena latifolia* | Marasmiineae | Agaricomycetes | Basidiomycota |
| KAF7336346 | 30.97 | 155 | 1.16E-04 | *Mycena venus* | Marasmiineae | Agaricomycetes | Basidiomycota |
| KAJ6517861 | 30.28 | 142 | 2.48E-07 | *Mycena vulgaris* | Marasmiineae | Agaricomycetes | Basidiomycota |
| KAJ7614860 | 32.43 | 148 | 7.44E-07 | *Roridomyces roridus* | Marasmiineae | Agaricomycetes | Basidiomycota |

**Table S12.** Nucleotide sequences used in phylogenetic analysis of yeast species delimitation.

| **Yeast species** | **Isolate** | **GenBank accession** |
|---|---|---|
| *Babjeviella inositovora* | NRRL Y-12698 | JQ689053 |
| *Blastobotrys proliferans* | LESF 1481 | ON493984 |
| *Blastobotrys proliferans* | NRRL Y-17577 | DQ442684 |
| *Candida aaseri* | NRRL YB-3897 | U45802 |
| *Candida africana* | UPV-EHU 97135 | AM495238 |
| *Candida albicans* | LESF 1442 | ON493939 |
| *Candida albicans* | NRRL Y-12983 | U45776 |
| *Candida auris* | 143 | AB375773 |

| | | |
|---|---|---|
| *Candida blattae* | LESF 1417 | ON493963 |
| *Candida blattae* | ATCC MYA-4360 | FJ614695 |
| *Candida blattariae* | NRRL Y-27703 | AY640213 |
| *Candida dosseyi* | NRRL Y-27950 | DQ655681 |
| *Candida endomychidarum* | NRRL Y-27708 | AY520330 |
| *Candida haemuloni* | CBS 5149 | KY106490 |
| *Candida insectorum* | NRRL Y-7787 | U45791 |
| *Candida intermedia* | NRRL Y-981 | U44809 |
| *Candida loeiensis* | NBRC 106731 | AB602835 |
| *Candida margitis* | LESF 1378 | ON493906 |
| *Candida margitis* | NYNU 15857 | KU128721 |
| *Candida membranifaciens* | NRRL Y-2089 | U45792 |
| *Candida nanaspora* | NRRL Y-17679 | U70187 |
| *Candida orthopsilosis* | LESF 1402 | ON493923 |
| *Candida orthopsilosis* | MCO456 | DQ213056 |
| *Candida oxycetoniae* | AS 2.3656 | EU402933 |
| *Candida parapsilosis* | LESF 1347 | ON493997 |
| *Candida parapsilosis* | NRRL Y-12969 | U45754 |
| *Candida powellii* | UWO(PS)99-325.3 | AF251554 |
| *Candida pseudoaaseri* | CBS 11170 | JN241689 |
| *Candida pseudointermedia* | CBS 6918 | KY106703 |
| *Candida quercuum* | NRRL Y-12942 | EF550292 |
| *Candida railenensis* | LESF 1437 | ON493931 |
| *Candida railenensis* | NRRL Y-17762 | U45800 |
| *Candida ruelliae* | MTCC 7739 | AM262326 |
| *Candida sinolaborantium* | LESF 1467 | ON493969 |

| | | |
|---|---|---|
| *Candida sinolaborantium* | ATCC MYA-4337 | FJ614676 |
| *Candida* sp. | BG01-7-23-017A-1-2 | AY242310 |
| *Candida* sp. | NRRL Y-27166 | EU011603 |
| *Candida* sp. | NRRL Y-12764 | EU011605 |
| *Candida* sp. | 05-7-186T | FJ537069 |
| *Candida* sp. | DMKU-WBL1-3 | MH744637 |
| *Candida* sp. (closest to *C. intermedia*) | LESF 1395 | ON493958 |
| *Candida* sp. (closest to *C. vrieseae*) | LESF 1433 | ON493964 |
| *Candida* sp. (closest to *C. auris*) | LESF 1413 | ON493965 |
| *Candida* sp. (closest to *C. temnochilae*) | LESF 1461 | ON493970 |
| *Candida* sp. (closest to *C. vrieseae*) | LESF 1493 | ON493972 |
| *Candida temnochilae* | CBS 9938 | AY242344 |
| *Candida vrieseae* | BI146 | EU200785 |
| *Cyberlindnera fabianii* | NRRL Y-1871 | EF550321 |
| *Cyberlindnera* sp. | LESF 1438 | ON493974 |
| *Cyberlindnera xylosilytica* | NRRL YB-2097 | EF550324 |
| *Debaryomyces hansenii* | LESF 1405 | ON493924 |
| *Debaryomyces hansenii* | NRRL Y-7426 | U45808 |
| *Debaryomyces prosopidis* | JCM 9913 | AB054993 |
| *Diutina catenulata* | LESF 1446 | ON493934 |
| *Diutina catenulata* | CBS 565 | KT336721 |
| *Grigorovia humatica* | IFO 10673 | AB040999 |
| *Grigorovia* sp. | LESF 1345 | ON493914 |
| *Grigorovia yakushimaensis* | IFO 1889 | AB041000 |
| *Hanseniaspora lachancei* | CBS 8818 | KY107804 |
| *Hanseniaspora opuntiae* | LESF 1470 | ON493938 |

| | | |
|---|---|---|
| *Hanseniaspora opuntiae* | CBS 8733 | AJ512453 |
| *Hanseniaspora thailandica* | ST-464 | DQ404527 |
| *Hanseniaspora thailandica* | LESF 1297 | ON493932 |
| *Hyphopichia burtonii* | LESF 1503 | ON493977 |
| *Hyphopichia burtonii* | CBS 2352 | KY107882 |
| *Hyphopichia fennica* | CBS 6027 | KY106445 |
| *Hyphopichia heimii* | NRRL Y-7502 | JQ689033 |
| *Hyphopichia lachancei* | CBS 5999 | MN645470 |
| *Hyphopichia pseudorhagii* | NRRL YB-2076 | AY789656 |
| *Hyphopichia rhagii* | CBS 618 | KY106723 |
| *Hyphopichia* sp. | LESF 1344 | ON493978 |
| *Issatchenkia orientalis* | NRRL Y-5396 | U76347 |
| *Kazachstania spencerorum* | LESF 1468 | ON493979 |
| *Kazachstania spencerorum* | NRRL Y-17920 | AY048162 |
| *Kluyveromyces hubeiensis* | AS 2.1536 | AH013031 |
| *Kodamaea ohmeri* | LESF 1474 | ON493980 |
| *Kodamaea ohmeri* | NRRL Y-1932 | U45702 |
| *Kurtzmaniella natalensis* | NRRL Y-17680 | U45818 |
| *Kurtzmaniella quercitrusa* | LESF 1421 | ON493967 |
| *Kurtzmaniella quercitrusa* | NRRL Y-5392 | U45831 |
| *Kurtzmaniella* sp | CLIB 1620 | LN870347 |
| *Kurtzmaniella* sp | LESF 1473 | ON493966 |
| *Kurtzmaniella* sp | LESF 1491 | ON493996 |
| *Limtongozyma cylindracea* | LESF 1500 | ON493917 |
| *Limtongozyma cylindracea* | NRRL Y-17506 | EU011644 |
| *Limtongozyma* sp. | LESF 1404 | ON493920 |

| | | |
|---|---|---|
| *Lodderomyces elongisporus* | LESF 1360 | ON493922 |
| *Lodderomyces elongisporus* | NRRL YB-4239 | LEU45763 |
| *Metschnikowia cibodasensis* | UICC Y-340 | AB236915 |
| *Metschnikowia koreensis* | LESF 1422 | ON493982 |
| *Metschnikowia koreensis* | KCTC 7828 | AF257272 |
| *Metschnikowia kunwiensis* | CBS 9679 | AJ716106 |
| *Metschnikowia reukaufii* | NRRL Y-7112 | U44825 |
| *Metschnikowia* sp. | LESF 1355 | ON493911 |
| *Metschnikowia* sp. | LESF 1343 | ON493912 |
| *Metschnikowia* sp. | LESF 1383 | ON493983 |
| *Meyerozyma caribbica* | LESF 1452 | ON493927 |
| *Meyerozyma caribbica* | NRRL Y-27274 | EU348786 |
| *Meyerozyma carpophila* | LESF 1389 | ON493918 |
| *Meyerozyma carpophila* | CBS 5256 | KY106386 |
| *Meyerozyma guilliermondii* | LESF 1444 | ON493925 |
| *Meyerozyma guilliermondii* | NRRL Y-2075 | JQ689047 |
| *Meyerozyma smithsonii* | BG02-7-13-007B-1-2 | AY518525 |
| *Pichia cecembensis* | NRRL Y-27985 | AM159112 |
| *Pichia kudriavzevii* | LESF 1411 | ON493941 |
| *Priceomyces fermenticarens* | NRRL Y-17321 | U45756 |
| *Priceomyces melissophilus* | NRRL Y-7585 | U45740 |
| *Priceomyces* sp. | LESF 1391 | ON493954 |
| *Saccharomycetales* sp. | LESF 1445 | ON493907 |
| *Scheffersomyces cryptocerci* | NRRL Y-48824 | KC702810 |
| *Scheffersomyces lignosus* | ATCC 58779 | HQ652022 |
| *Scheffersomyces shehatae* | NRRL Y-12858 | JQ025409 |

| | | |
|---|---|---|
| *Scheffersomyces* sp. | LESF 1457 | ON493988 |
| *Schizosaccharomyces pombe* | NRRL Y-12796 | JQ689077 |
| *Schwanniomyces vanrijiae* | LESF 1478 | ON493989 |
| *Schwanniomyces vanrijiae* | NRRL Y-7430 | U45842 |
| *Starmerella apicola* | NRRL Y-2481 | U45703 |
| *Starmerella bombicola* | LESF 1472 | ON493929 |
| *Starmerella bombicola* | NRRL Y-17069 | U45705 |
| *Starmerella etchellsii* | LESF 1300 | ON493946 |
| *Starmerella etchellsii* | NRRL Y-17084 | U45723 |
| *Starmerella floris* | UWO(PS)00-226.2 | AF313353 |
| *Starmerella ilheusensis* | UFMG-CM-Y596 | KR232374 |
| *Starmerella meliponinorum* | LESF 1496 | ON493947 |
| *Starmerella meliponinorum* | UWO(PS)00-227.1 | AF313354 |
| *Starmerella* sp. | LESF 1293 | ON493951 |
| *Starmerella* sp. | LESF 1454 | ON493995 |
| *Sugiyamaella americana* | NRRL YB-2067 | DQ438193 |
| *Sugiyamaella bonitensis* | LESF 1465 | ON493990 |
| *Sugiyamaella bonitensis* | UFMG-CM-Y608 | KT006004 |
| *Sugiyamaella boreocaroliniensis* | LESF 1366 | ON493916 |
| *Sugiyamaella boreocaroliniensis* | NRRL YB-1835 | DQ438221 |
| *Sugiyamaella carassensis* | LESF 1463 | ON493991 |
| *Sugiyamaella carassensis* | UFMG-CM-Y606 | KX550111 |
| *Sugiyamaella floridensis* | NRRL YB-3827 | DQ438222 |
| *Sugiyamaella ligni* | LESF 1459 | ON493992 |
| *Sugiyamaella ligni* | UFMG-CM-Y295 | KX550112 |
| *Sugiyamaella smithiae* | LESF 1336 | ON493915 |

| | | |
|---|---|---|
| *Sugiyamaella smithiae* | NRRL Y-17850 | DQ438218 |
| *Wickerhamiella azyma* | CBS 6826 | EF536346 |
| *Wickerhamiella occidentalis* | LESF 1296 | ON493930 |
| *Wickerhamiella occidentalis* | CBS 8452 | AF046037 |
| *Wickerhamiella* sp. | LESF 1495 | ON493913 |
| *Wickerhamiella* sp. | LESF 1506 | ON493952 |
| *Wickerhamomyces canadensis* | NRRL Y-1888 | EF550300 |
| *Wickerhamomyces ciferrii* | LESF 1397 | ON493994 |
| *Wickerhamomyces ciferrii* | NRRL Y-1031 | EF550339 |
| *Wickerhamomyces ochangensis* | N7a-Y2T | HM485464 |
| *W. psychrolipolyticus* | YS-2018 202-2 | LC333101 |
| *Wickerhamomyces siamensis* | DMKU-RK359 | AB714248 |
| *Wickerhamomyces* sp. | LESF 1460 | ON493968 |
| *Yamadazyma laniorum* | yHMH7 | KY588136 |
| *Yamadazyma paraaseri* | NYNU 1811114 | MK682805 |
| *Yamadazyma paraaseri* | LESF 1412 | ON493961 |
| *Yamadazyma* sp. | LESF 1490 | ON493971 |
| *Yamadazyma tenuis* | NRRL Y-1498 | U45774 |
| *Yamadazyma tumulicola* | T6517-9-5 | AB365463 |
| *Yarrowia lipolytica* | LESF 1385 | ON493926 |
| *Yarrowia lipolytica* | CBS 6124 | AM268450 |
| *Yarrowia lipolytica* | NRRL Y-2317 | U45793 |
| *Yarrowia yakushimensis* | CBS 10254 | AM268474 |

**Table S13.** The effect of pH on the growth inhibition of *Kodamaea ohmeri* LESF 1474 and *Candida castyelli* by Ksino expressed by *S. cerevisiae*.

| | *Kodamaea ohmeri* | *Candida castelli* |
|---|---|---|
| Treatment | Area in cm² (Mean ± SE) | |
| pH 3.5 | 0 | 3.65 ± 0.57 |
| pH 4.0 | 0 | 8.42 ± 0.70 |
| pH 4.5 | 0 | 7.29 ± 1.26 |
| pH 5.0 | 0 | 3.30 ± 0.06 |
| pH 5.5 | 0 | 0 |
| pH 6.0 | 0 | 0 |
| pH 6.5 | 0 | 0 |

**Table S14.** The effect of temperature on the growth inhibition of *Kodamaea ohmeri* LESF 1474 and *Candida castyelli* by Ksino expressed by *S. cerevisiae*.

| | *Kodamaea ohmeri* | *Candida castelli* |
|---|---|---|
| Treatment | Area in cm² (Mean ± SE) | |
| 17°C | 0 | 2.18 ± 0.56 |
| 20°C | 0 | 7.35 ± 0.20 |
| 25°C | 0 | 7.53 ± 0.72 |
| 30°C | 0 | 1.75 ± 0.17 |
| 35°C | 0 | 0 |

**Table S15.** Yeasts used in killer assays as suceptible or killer positive standards.

| **Yeast species** | **Isolate** | **Use in the study** |
|---|---|---|
| *Blastobotrys proliferans* | LESF 1481 | Lawn strain |
| *Candida blattae* | LESF 1417 | Lawn strain |
| *Candida railenensis* | LESF 1437 | Lawn strain |

| | | |
|---|---|---|
| *Candida railenensis* | LESF 1551 | Lawn strain |
| *Candida sinolaborantium* | LESF 1467 | Lawn strain |
| *Cyberlindnera* sp. | LESF 1556 | Lawn strain |
| *Cyberlindnera* sp. | LESF 1555 | Lawn strain |
| *Debaryomyces hansenii* | LESF 1405 | Lawn strain |
| *Diutina catenulata* | LESF 1446 | Lawn strain |
| *Hanseniaspora opuntiae* | LESF 1470 | Lawn strain |
| *Hanseniaspora thailandica* | LESF 1297 | Lawn strain |
| *Issatchenkia orientalis* | LESF 1411 | Lawn strain |
| *Kazachstania spencerorum* | LESF 1468 | Lawn strain |
| *Kodamaea ohmeri* | LESF 1474 | Lawn strain |
| *Kurtzmaniella quercitrusa* | LESF 1489 | Lawn strain |
| *Limtongozyma cylindracea* | LESF 1500 | Lawn strain |
| *Metschnikowia koreensis* | LESF 1422 | Lawn strain |
| *Meyerozyma caribbica* | LESF 1452 | Lawn strain |
| *Meyerozyma caribbica* | LESF 1545 | Lawn strain |
| *Meyerozyma carpophila* | LESF 1389 | Lawn strain |
| *Meyerozyma guilliermondii* | LESF 1406 | Lawn strain |
| *Ogataea* sp. | LESF 1525 | Lawn strain |
| *Schwanniomyces vanrijiae* | LESF 1521 | Lawn strain |
| *Starmerella bombicola* | LESF 1472 | Lawn strain |
| *Starmerella etchellsii* | LESF 1300 | Lawn strain |
| *Starmerella meliponinorum* | LESF 1496 | Lawn strain |
| *Sugiyamaella bonitensis* | LESF 1465 | Lawn strain |
| *Sugiyamaella carassensis* | LESF 1463 | Lawn strain |
| *Sugiyamaella ligni* | LESF 1459 | Lawn strain |
| *Wickerhamomyces ciferrii* | LESF 1397 | Lawn strain |

| | | |
|---|---|---|
| *Yamadazyma paraaseri* | LESF 1412 | Lawn strain |
| *Yamadazyma riverae* | LESF 1541 | Lawn strain |
| *Yamadazyma riverae* | LESF 1538 | Lawn strain |
| *Yarrowia lipolytica* | LESF 1385 | Lawn strain |
| *Candida castellii* | Y-17070 | Lawn strain |
| *Saccharomyces arboricolus* | 2.3317 | Lawn strain |
| *Kazachstania africana* | Y-2729 | Lawn strain |
| *Saccharomyces cerevisiae* | Y-2046 | Lawn strain |
| *Saccharomyces cerevisiae* | MSY"99" | Lawn strain |
| *Saccharomyces cerevisiae* | Y-5509 | Lawn strain |
| *Saccharomyces cerevisiae* | Y-1891 | Lawn strain |
| *Saccharomyces cerevisiae* | BY 4741 | Lawn strain |
| *Saccharomyces mikatae* | NBRC 1815 | Lawn strain |
| *Saccharomyces paradoxus* | YB - 4565 | Lawn strain |
| *Saccharomyces paradoxus* | Y8.5 | Lawn strain |
| *Saccharomyces eubayanus* | CBS 12357 | Lawn strain |
| *Saccharomyces kudriavzevii* | NBRC/IFO 10990 | Lawn strain |
| *Naumovozyma castellii* | NCYC2898 | Lawn strain |
| *Nakaseomyces delphensis* | Y-2379 | Lawn strain |
| *Saccharomyces cerevisiae* | YJM1307 | K1 toxin-producing strain |
| *Saccharomyces cerevisiae* | CYC1172 | K2 toxin-producing strain |
| *Saccharomyces cerevisiae* | DSM-70459 | Klus toxin-producing strain |
| *Saccharomyces cerevisiae* | OS169 | K62 toxin-producing strain |
| *Saccharomyces cerevisiae* | OS78 | K45 toxin-producing strain |
| *Saccharomyces cerevisiae* | OS294 | K74 toxin-producing strain |
| *Saccharomyces cerevisiae* | MSC300c | K28 toxin-producing strain |
| *Saccharomyces cerevisiae* | OS40 | K21 toxin-producing strain |

| | | |
|---|---|---|
| *Saccharomyces paradoxus* | Y-63717 | K1L toxin-producing strain |
| *Candida albicans* | Y-12983 | Pathogenic lawn strain |
| *Candida auris* | AR0381 | Pathogenic lawn strain |
| *Candida auris* | AR0382 | Pathogenic lawn strain |
| *Candida auris* | AR0383 | Pathogenic lawn strain |
| *Candida auris* | AR0384 | Pathogenic lawn strain |
| *Candida auris* | AR0385 | Pathogenic lawn strain |
| *Candida auris* | AR0386 | Pathogenic lawn strain |
| *Candida auris* | AR0387 | Pathogenic lawn strain |
| *Candida auris* | AR0388 | Pathogenic lawn strain |
| *Candida auris* | AR0389 | Pathogenic lawn strain |
| *Candida auris* | AR0390 | Pathogenic lawn strain |
| *Candida castellii* | Y-17070 | Pathogenic lawn strain |
| *Candida glabrata* | AR0323 | Pathogenic lawn strain |
| *Candida glabrata* | AR0317 | Pathogenic lawn strain |
| *Candida glabrata* | ATCC2001 | Pathogenic lawn strain |
| *Candida humilis* | Y-17074 | Pathogenic lawn strain |
| *Candida kefyr* | AR0588 | Pathogenic lawn strain |
| *Candida nivarensis* | Y-48269 | Pathogenic lawn strain |
| *Candida pararugose* | AR0587 | Pathogenic lawn strain |
| *Saccharomyces cerevisiae* | AR0400 | Pathogenic lawn strain |
