## Supplementary figures and images for "The Prevalence of Killer Yeasts in the Gardens of Fungus-Growing Ants and the Discovery of a Novel Killer Toxin named Ksino"

### Figures S1-7

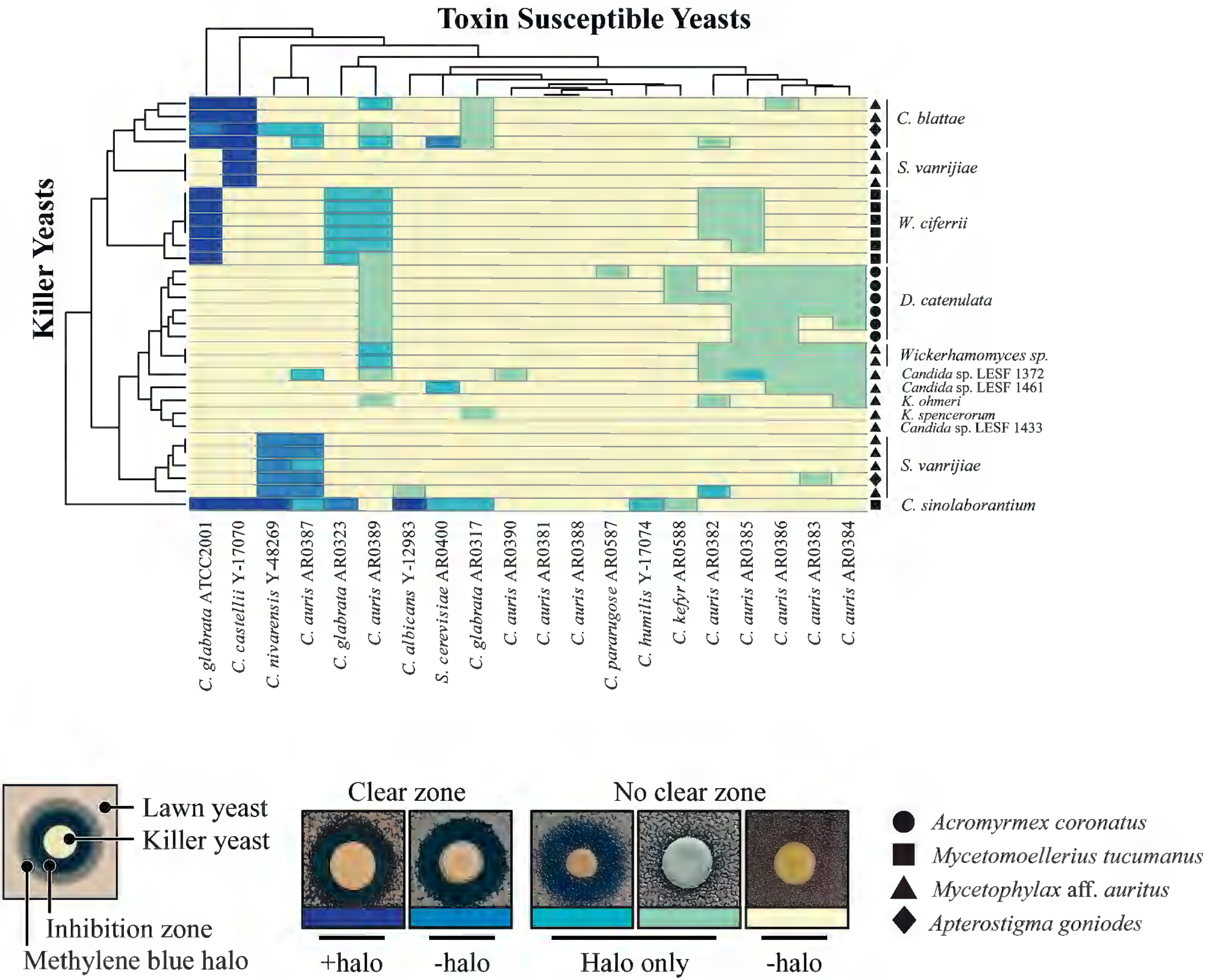

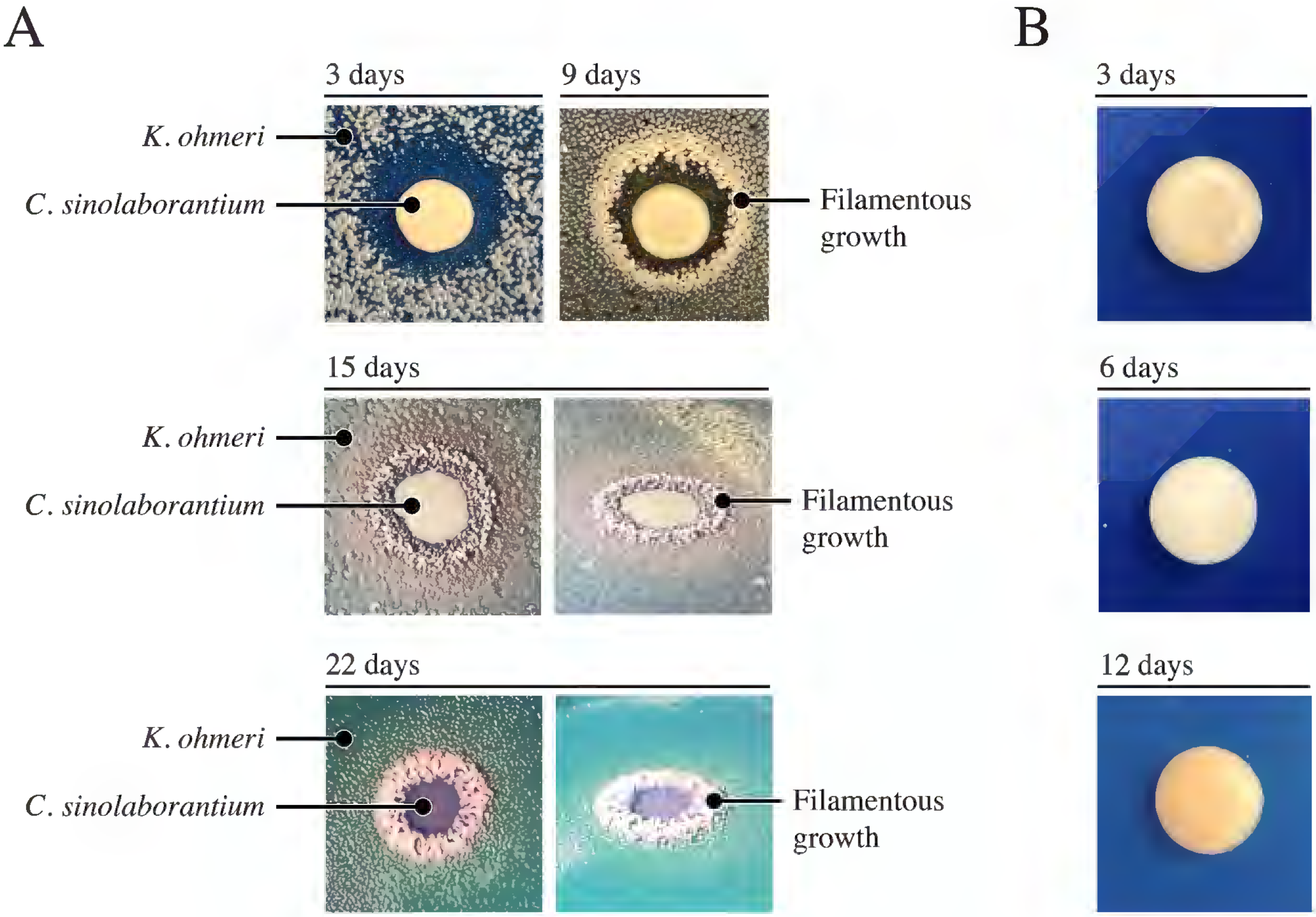

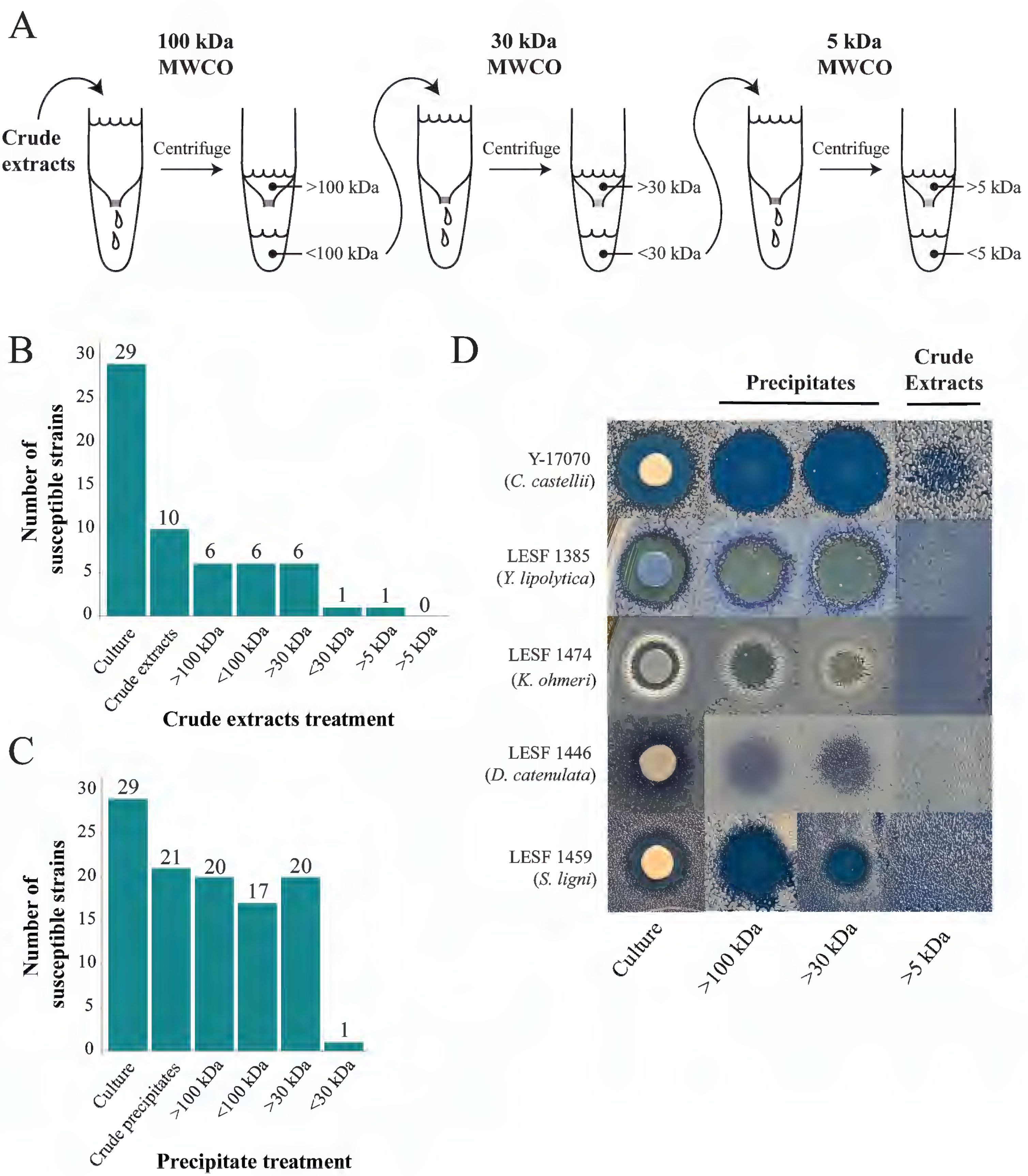

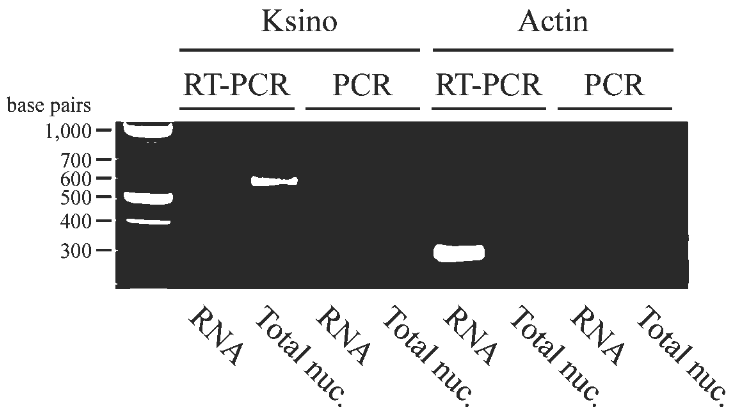

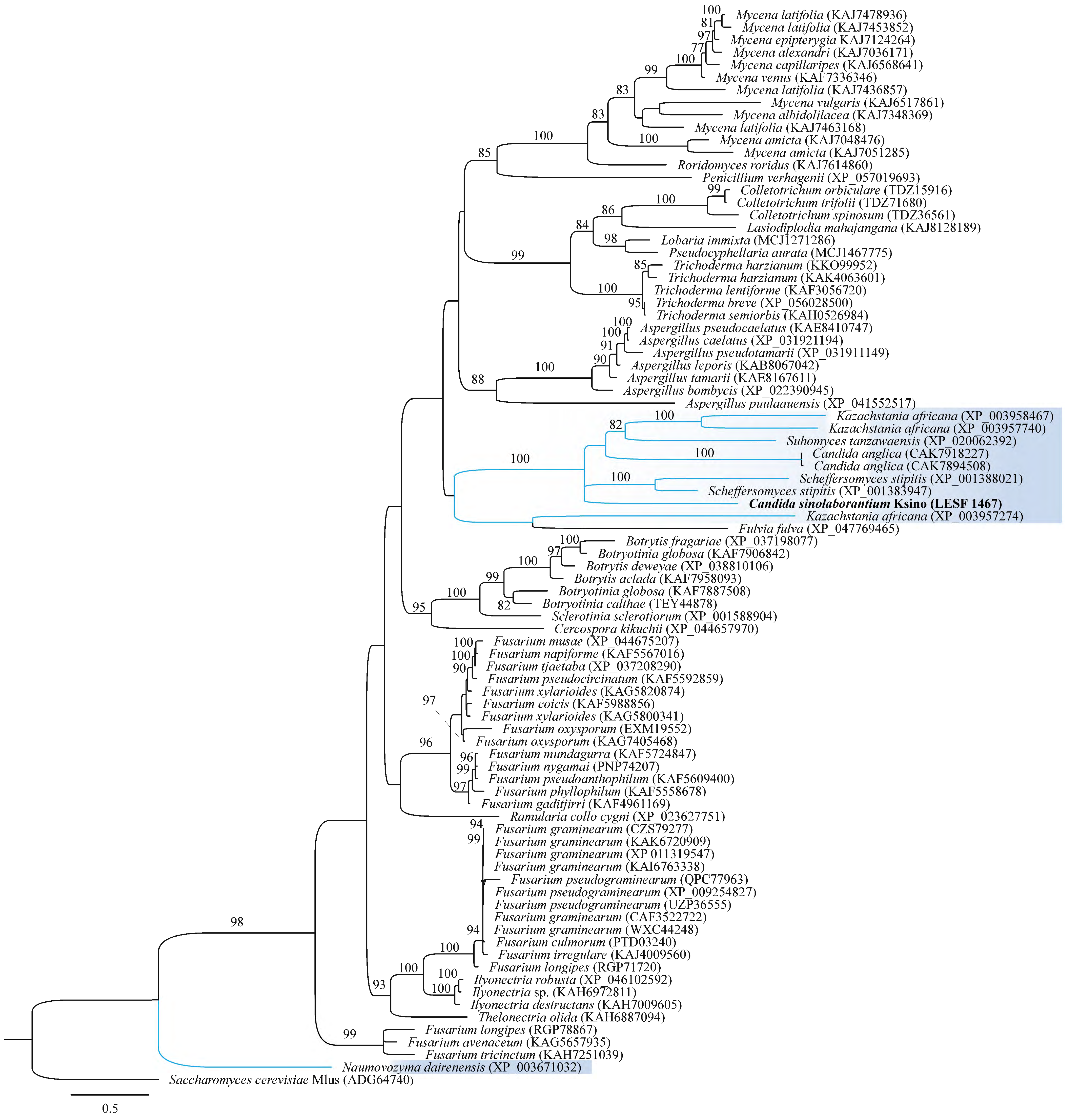

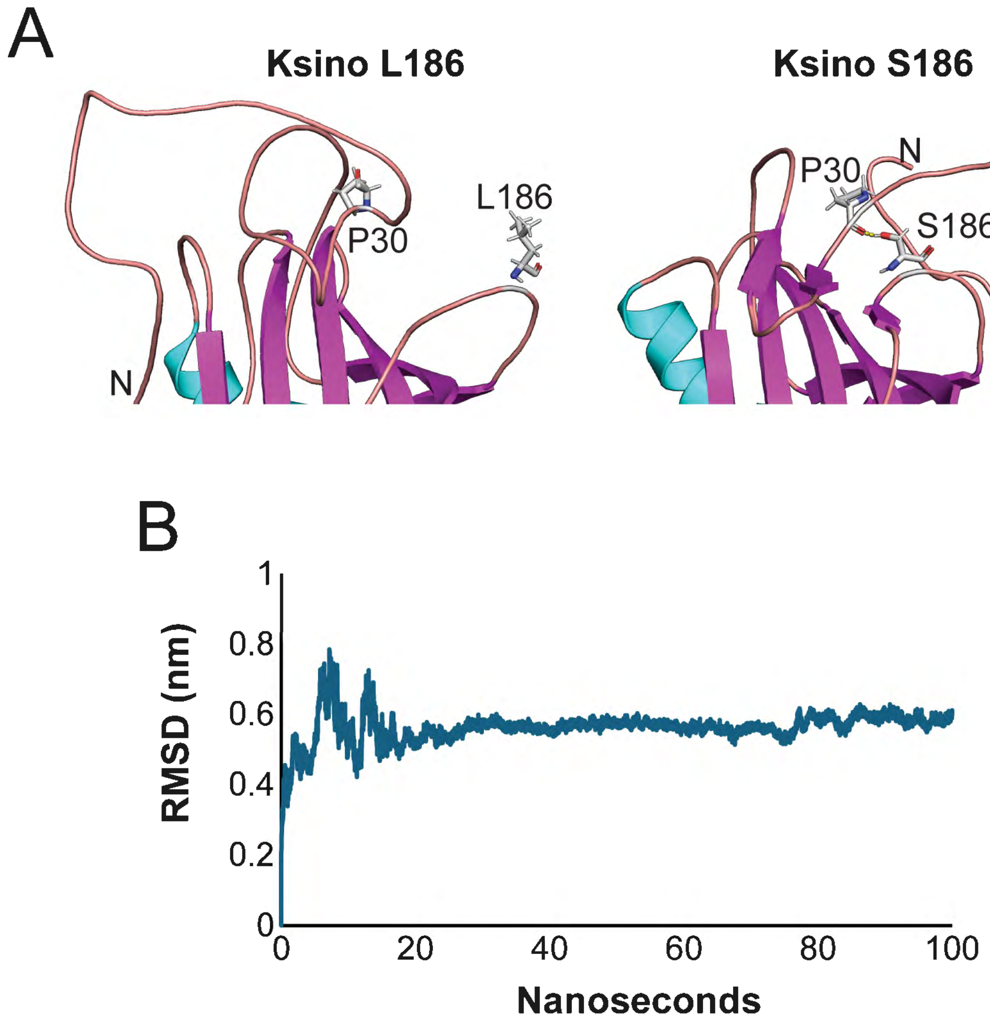

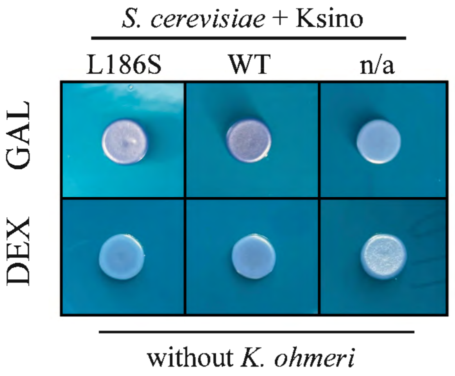
